# Reassessing Choice Probability: What 59 Macaque Studies Tell Us About Decision-Related Activity in Visual Cortex

**DOI:** 10.64898/2026.06.08.730961

**Authors:** Anton Pletenev, Richard T. Born, Gregory C. DeAngelis, Raymond Doudlah, Ichiro Fujita, Joshua I. Gold, Robbe L. T. Goris, Alexander C. Huk, Kristine Krug, Pooya Laamerad, Richard D. Lange, Aaron J. Levi, Hendrikje Nienborg, Christopher C. Pack, Andrew J. Parker, Ari Rosenberg, Mehdi Sanayei, Takanori Uka, Xuefei Yu, Adam Zaidel, Corey M. Ziemba, Ralf M. Haefner

## Abstract

Choice probability (CP) quantifies the trial-by-trial covariation between a sensory neuron’s response and perceptual reports. Despite extensive interest in CP as a window into how perception is linked to the activity of sensory neurons, the usefulness of CP remains debated. On one hand, reported CP magnitudes vary widely across seemingly equivalent studies, questioning its utility as a metric. On the other hand, the absence of clear patterns in CP variability has made it difficult to use it to adjudicate between competing models of visual perception. Here, we performed a meta-analysis of 150 CP estimates from 59 macaque neurophysiology studies to identify factors that systematically influence CP. We confirmed the positive relationship between CP and neuronal sensitivity both across and within individual studies. When controlled for sensitivity, we found remarkable consistency in CP across varying tasks and brain regions with two notable exceptions. First, CPs were higher in tasks involving bistable percepts, reinforcing the link between CP magnitude and subjective perception. Second, CPs were smaller in area V1, supporting prior suggestions about V1’s special role in visual processing. We further found a significant effect of stimulus duration on CP, providing evidence against strictly feedforward models and favoring models with substantial feedback and recurrent processing. Finally, we offer recommendations for future studies to enhance the cross-study comparability and theoretical utility of choice signals as the field transitions to large-scale population recordings. More broadly, our findings demonstrate the benefits of meta-studies that expose patterns across many different tasks and animals – yielding insights that complement large-scale population recordings.

## 1 Introduction

Perception arises from the activity of many sensory neurons, yet remarkably, the responses of *individual* neurons often covary with what an animal perceives or decides. This observation has long fueled the quest to identify whether and how the activity of a single neuron—amid many potential contributors—plays a causal role in shaping perceptual experience (Parker and Newsome, 1998). Understanding this link between single-neuron activity and behavior has therefore become a major focus of systems neuroscience. Because an animal’s subjective percept cannot be directly observed, researchers have instead examined how neuronal responses relate to both the sensory stimulus and the animal’s choices in tasks that rely on the perceptual judgment of the stimulus. If a neuron is informative about both the stimulus and the animal’s choice, it may contribute to perception (Parker and Newsome, 1998; Panzeri et al., 2017). The latter may be assessed by measuring choice correlations (Fig. 1a) – the correlation between a neuron’s responses and the subject’s choice across repeated presentations of an identical stimulus, thereby removing confounding effects of stimulus variability (Pitkow et al., 2015). Choice probability (CP) is conceptually similar to this correlation: it quantifies the probability of correctly predicting the animal’s choice such that CP = 0.5 corresponds to no correlation (Britten et al., 1996). Crucially, CP captures not just the strength of the choice signal, but also its alignment with the neuron’s stimulus tuning (Fig. 1a): CP *>* 0.5 if the response is higher preceding the choice corresponding to stimuli to which the neuron responds more.

**Figure 1:**
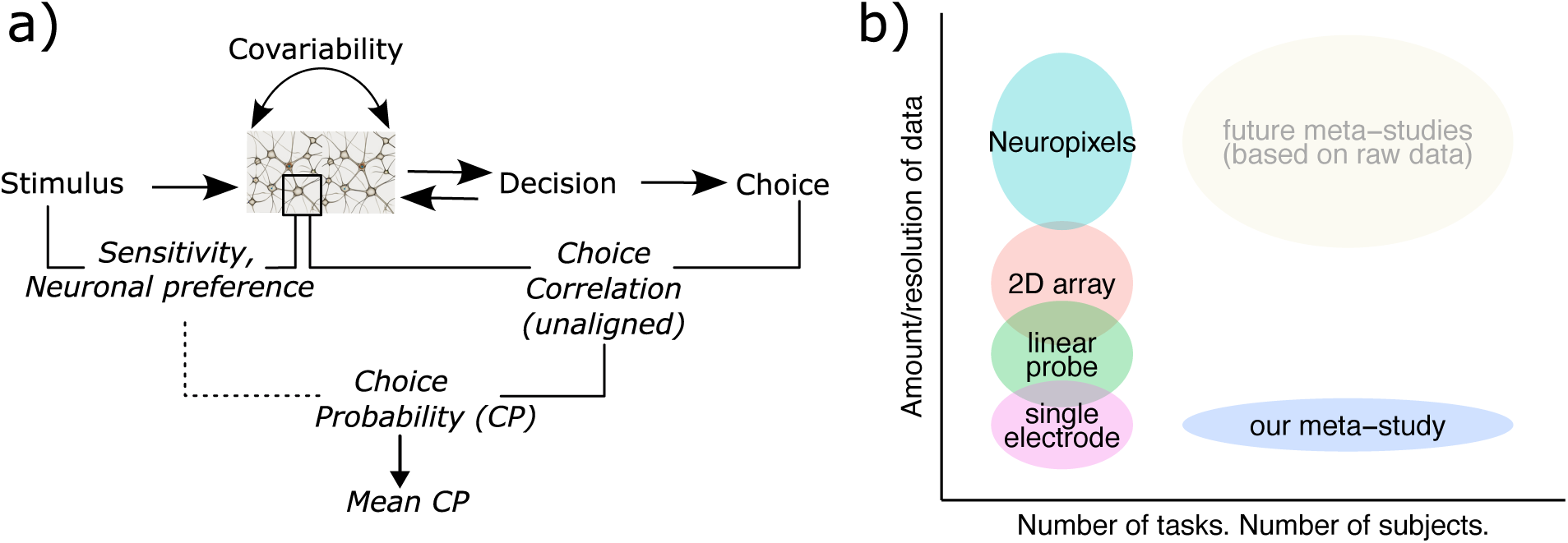
Conceptual framework of choice-related metrics and the rationale for a meta-analytic approach. **(a)** In perceptual decision tasks, sensory neuron responses reflect both the external stimulus and the animal’s choice. *Sensitivity* and *preference* characterize the neuron-stimulus relationship: sensitivity quantifies the magnitude of response modulation, while preference indicates the stimulus category eliciting higher firing rates (ideally both assessed when choice is held constant). *Choice Correlations* and *Choice Probability (CP)* quantify the relationship between neuronal activity and the subject’s choice for a fixed stimulus. Choice correlations measure the correlation between trial-by-trial neuronal firing rates and the subject’s binary choice (coded as +1 or -1), using a fixed sign convention for choice across the entire population. CP differs in two key respects: (1) it is calculated as an ROC area, and (2) the “positive” choice class is defined by the neuron’s own stimulus preference (its influence on CP is indicated by the dashed line). CP is shaped by *covariability* —the trial-by-trial correlation between neuronal responses to a fixed stimulus (Equation (1)). *Mean CP* —the core metric of our meta-analysis— is a population-level measure indicating how strongly, on average, neurons increase their firing when the subject selects the neuron’s preferred category. **(b)** Conceptual illustration of the complementary strengths of this meta-analysis versus individual experimental studies. The y-axis (Amount/resolution of data) reflects both the number of data points and the ability to capture simultaneous population dynamics. The x-axis represents experimental diversity, specifically the number of different behavioral tasks and individual subjects tested. Studies based on modern recording techniques offer superior data resolution but typically rely on few tasks and few animals (high y-axis, low x-axis). Conversely, our meta-analysis aggregates mean CP across 59 studies to span diverse tasks and subjects (high x-axis, low y-axis). Thus, while limited in the amount of data, it maximizes generalizability by overcoming subject-specific variance and enabling the isolation of experimental factors that impact the choice signal in sensory neurons. Future meta-studies may overcome these data limitations by pooling raw datasets rather than relying on summary statistics (see Section 3.7).

Interpreting CP as a perceptual metric requires ruling out the possibility that it reflects a purely motor signal—that is, a correlation driven by the physical behavioral response rather than the decision itself (Parker and Newsome, 1998). While such oculomotor planning signals likely dominate choice activity in higher-order sensorimotor areas like VIP and LIP, lower sensory areas appear less susceptible to this motor confound (Nichols and Newsome, 2002; Zaidel et al., 2017; Katz et al., 2016; Yu and Gu, 2018; but see Laamerad et al., 2024). Numerous macaque studies have reported CP values significantly above chance in various sensory cortical areas and behavioral tasks, attracting considerable interest in CP as a potential indicator of a neuron’s role in perception (Nienborg et al., 2012). The present study focuses exclusively on the macaque visual cortex, as this represents the most extensive and foundational body of CP research; however, significant CPs have also been documented in the sensory cortex of other sensory modalities (Romo et al., 2002; de Lafuente and Romo, 2006; Niwa et al., 2012), subcortical structures (Liu et al., 2013; Angelaki et al., 2004) and across various animal models (Yang et al., 2016; Steinfeld et al., 2024).

The view that CP is a meaningful measure of a neuron’s perceptual involvement is also supported by the observation that more task-relevant neurons tend to exhibit higher values of CP. Numerous studies have reported a positive correlation between CP and a neuron’s task sensitivity – that is, its ability to discriminate between task-relevant stimuli (Britten et al., 1996; Purushothaman and Bradley, 2005; Gu et al., 2008; Price and Born, 2010; Kang and Maunsell, 2020; Laamerad et al., 2024; Ghose and Harrison, 2009; Kim et al., 2022; Parker et al., 2002; Bosking and Maunsell, 2011; Shiozaki et al., 2012; Xu et al., 2014; Krug et al., 2016). Several studies have further shown that this relationship emerges over training (Law and Gold, 2008; Uka et al., 2012; Sanayei et al., 2018). These findings reinforce the idea that CP reflects a neuron’s contribution to perception by linking its informativeness about both the stimulus and the choice (Parker and Newsome, 1998). If this interpretation holds, CP should also depend on the animal’s task strategy – as supported by some evidence that CP selectively emerges in neurons carrying the specific sensory cues the animal subjectively relies upon, even when other neurons are objectively just as informative (Uka and DeAngelis, 2004; Nienborg and Cumming, 2007). Furthermore, microstimulation results obtained under the same experimental setups as CP measurements provide evidence for a causal link between recorded neurons and perception (Salzman et al., 1992; Uka and DeAngelis, 2006; Krug et al., 2013).

Several theoretical frameworks have been used to interpret CP findings. The foundational framework explains choice probabilities primarily in terms of feedforward connectivity, attributing them to both the causal effect of a neuron’s activity on perception, and its correlation with other neurons that contribute causally (Zohary et al., 1994; Britten et al., 1996; Shadlen et al., 1996; Liu and Newsome, 2005; Law and Gold, 2008; Cohen and Newsome, 2009; Rao et al., 2012; Haefner et al., 2013). Determining the true origin of CP, therefore, requires elucidating the mechanisms by which these correlations are generated (Nienborg et al., 2012). Within a strictly feedforward framework, such covariability is presumed to arise bottom-up from shared afferent inputs to sensory populations and subsequent dynamics within local lateral networks (Shadlen and Newsome, 1998; Kohn et al., 2016). Alternatively, these correlations may originate from top-down modulation (Ecker et al., 2014), where downstream areas provide choice-informative feedback to sensory neurons (Dodd et al., 2001; Nienborg et al., 2012; Chicharro et al., 2021). Some researchers have proposed that attention or attention-like fluctuations may be the main source of such feedback (Nienborg and Cumming, 2009, 2010; Ecker et al., 2016; Krug, 2004). The Bayesian brain hypothesis and associated hierarchical approximate inference models assume that the sensory cortex represents posterior beliefs, where top-down modulations communicate prior beliefs from higher order areas (Lange and Haefner, 2022; Haefner et al., 2016; Festa et al., 2020; Bányai et al., 2019).

However, a substantial disconnect remains between theoretical predictions and empirical results. Reported CP values vary widely across studies, and it is unclear whether such variability can be reconciled with existing theoretical models. Some studies have failed to detect significant CPs or found no reliable link with neuronal sensitivity (Goris et al., 2017; Clery et al., 2017; Lange et al., 2023; Laamerad et al., 2024). Even when significant effects are observed, their magnitudes differ markedly. This variability raises the question of whether all studies are indeed probing the same underlying CP phenomenon, as theoretical models implicitly assume. Heterogeneity in CP findings likely reflects systematic differences in experimental design across studies – factors whose influence remains insufficiently understood, leaving many current interpretations largely speculative. Additionally, some variability may stem from the small number of animals typically used per study (often just two).

Understanding the dependence of CP on key experimental variables is essential for bridging the gap between theory and empirical findings. A central question is which neurons are actually read out by decision-making circuits or targeted by top-down feedback or a combination of both. Normative models based on optimality predict that these neurons are the most informative about the stimulus (Haefner et al., 2013; Lange and Haefner, 2022), but this assumption may be violated by architectural constraints in the brain or by incomplete learning. Examining how reported CP varies with factors such as brain area, task type, stimulus duration, or behavioral performance offers a way to test some of these constraints.

Although developed during the era of single-neuron recordings, the CP metric remains highly relevant today because its widespread adoption provides a universal standard for cross-study comparisons. Recent breakthroughs in population recording techniques in nonhuman primates have significantly increased data resolution, allowing researchers to investigate choice-related signals in simultaneously recorded populations at a level of detail that traditional CP cannot capture (Bondy et al., 2018; Zhao et al., 2020; Levi et al., 2023). However, the relatively small number of such studies and, more importantly, the lack of a standardized metric for quantifying choice signals in simultaneously recorded populations—hinders broader generalization. A subtle yet critical complication is that, unlike the simple averaging of individual CPs, metrics derived from simultaneous population activity – such as the accuracy of the population choice decoder – are inherently sensitive to the number of recorded neurons, a variable that fluctuates widely across experiments without an established method for cross-study normalization. Furthermore, despite their increased resolution, these modern studies share a fundamental limitation with classic experiments: they typically rely on a single behavioral task and a restricted number of subjects. Consequently, aggregating CP results across decades of literature currently remains the only viable approach for comparing diverse tasks and subjects on a large scale, even if such analysis inevitably sacrifices the fine-grained, neuron-level details available at the study level (Fig. 1b).

The present study leverages the extensive body of research on choice probability to identify systematic patterns linking CP estimates to specific experimental variables. We compiled 150 CP values from 59 studies (Table 1). Across these studies, we found substantial variability in both the magnitude and the statistical significance of reported CPs. Using a meta-analytic approach, we evaluated how differences in experimental design contribute to this heterogeneity. Based on a comprehensive literature review, we pre-identified candidate variables and assessed their impact across studies. Some factors were identified as potential confounds that may bias CP estimates, while others offered insights into the underlying causes of the CP phenomenon. Our analysis revealed four key variables influencing empirical CPs: a consistent positive relationship with neuronal sensitivity, preserved across most task types and brain areas, a positive relationship with stimulus duration, consistently lower CPs in V1 relative to extrastriate cortex, and higher CPs in tasks involving bistable stimuli compared to classic coarse or fine-discrimination tasks. These findings provide empirical constraints on theoretical models of CP and suggest directions for future experiments to causally manipulate key variables and test their predicted influence on choice-related neural activity.

### Glossary Box

**Choice Correlation**: The correlation between a sensory neuron’s response and the subject’s choice across repeated presentations of an identical stimulus.

**Choice Probability (CP)**: Classic metric to quantify choice correlations in terms of a probability by comparing the neuron’s response distributions preceding each choice. Crucially, CP aligns the choice correlation with the neuron’s tuning preference by assigning the positive label to the choice corresponding to the neuron’s preferred stimulus category.

**Neurometric Threshold**: A measure of a single neuron’s sensitivity, defined as the stimulus strength required for the neuron to discriminate between categories at a specific performance level (e.g., 82% correct). It is the inverse of neural sensitivity and is expressed in experiment-specific parameters like stimulus coherence or contrast.

**Normalized Sensitivity**: A measure of neuronal sensitivity relative to the animal’s behavioral sensitivity (performance), allowing for comparisons across different studies and tasks. It can be calculated as the ratio of neuronal to behavioral *d^′^*, or as the inverse neurometric to psychometric *threshold* ratio.

**Noise Correlations (Covariability)**: The trial-to-trial correlations (covariability) in the responses of pairs of sensory neurons while keeping the external stimulus fixed. These correlations (specifically those aligned with the task-relevant signal, called “differential correlations”) are closely related to Choice Probability in large neural populations.

## 2 Results

### 2.1 Conceptual understanding of Choice probabilities

Choice probability (CP) quantifies the relationship between a sensory neuron’s activity and the subject’s choice in a task, while keeping the stimulus constant (Britten et al., 1996; Parker and Newsome, 1998; Nienborg et al., 2012, Fig. 1a). It can be viewed as an ROC-based analogue of a partial correlation measure: it is approximately proportional to the correlation between neural response and choice controlling for the sensory input (stimulus) (Britten et al., 1996; Haefner et al., 2013). This correlation is also called “choice correlation” (Pitkow et al., 2015). In this paper, we use choice correlation for unaligned correlation where the assignment of positive and negative labels to choices is arbitrary, making only the magnitude of the correlation meaningful to interpret (Fig. 2, Glossary Box). CP is instead computed relative to a neuron’s stimulus preference, assigning the positive label to the choice corresponding to the stimulus which elicits the higher response (sometimes called the neuron’s “preferred stimulus category”, or “preferred choice”). Like any correlation-based metric, CP itself is agnostic to the causal link between neural activity and choice.

**Figure 2:**
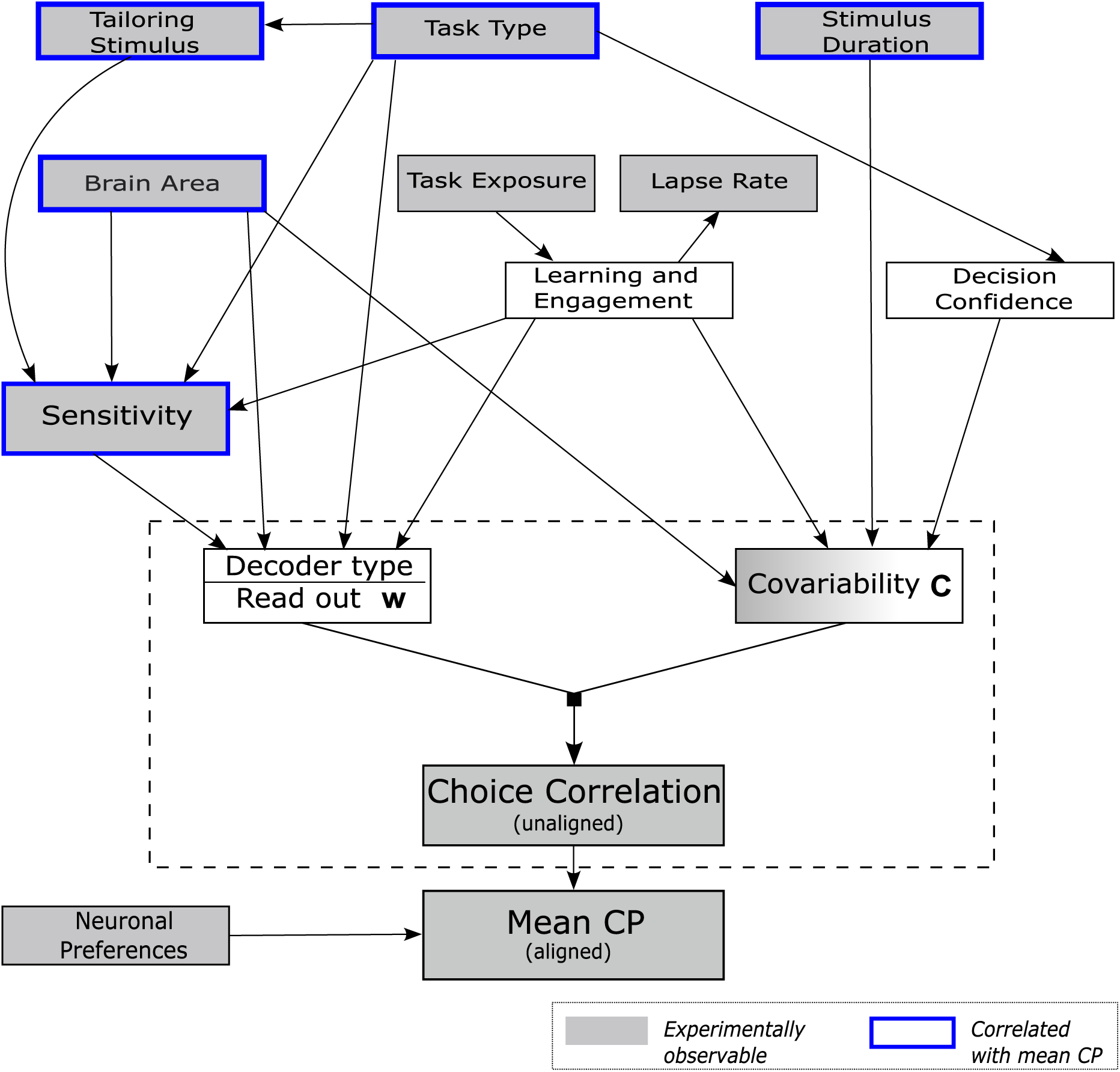
Schematic representation of factors hypothesized to influence Choice Probability (CP). Arrows denote putative causal relationships between experimental design choices, intermediate variables, and core factors influencing CP (see Tables 2 and 3 for details). Interaction effects between variables are not shown. Two central variables –decoder type/readout **w** and covariability **C** (enclosed by the dashed outline), linking CP to perception through Equation (1) – mediate the effect of all other factors on choice correlations. Gray boxes indicate variables that can be experimentally observed. Thick blue borders highlight variables that showed a statistically significant relationship with mean CP in our meta-analysis. For a more complete schematic incorporating additional factors (see Fig. S1).

We organized previously proposed factors that may influence CP magnitude according to distinct hypotheses about the underlying nature of CP (Fig. 2). Most efforts in the field have focused on understanding CP as a signature of the perceptual decision-making process (Shadlen et al., 1996) – either as a feedforward influence of sensory neurons on decisions, or as the result of decision-related signals propagating back to sensory areas (Nienborg et al., 2012). Although these theoretical frameworks differ conceptually, they agree in their prediction that two key factors shape choice probabilities: correlations between neurons and the structure of the readout profile (Chicharro et al., 2021).

In the classic feedforward paradigm, the output of a linear decoder, reading out activity from the sensory population, forms the decision. If the activity of sensory neurons were independent of each other (uncorrelated) then a CP value greater than 0.5 would imply that the corresponding neuron is causally contributing to the subject’s choice. But in reality where neural activity is correlated, the structure of these correlations critically contributes to the measured CP (Nienborg et al., 2012; Haefner et al., 2013). In this case, the choice probability (CP*_i_*) of a neuron *i* is only weakly determined by its own read-out weight (*w_i_*) and its stimulus preference (sign(*f^′^_i_*)). Instead, it is the aggregate result of the neuron’s correlations with other neurons from the decision pool (*C_ij_*) and those neurons’ read-out weights (*w_j_*) Haefner et al. (2013):

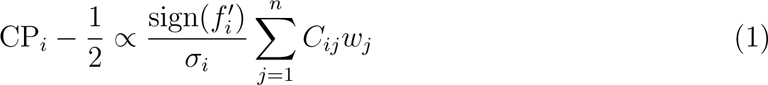

Thus, individual CPs within the readout population are determined by the interaction between the readout weight vector, **w**, and the covariance matrix, **C** (Fig. 2). Empirically, interneuronal covariability exhibits a highly specific, non-random structure, and this structure is critical for the observed patterns in CPs (Nienborg et al., 2012; Haefner et al., 2013; Kohn et al., 2016).

Under a wide range of scenarios, readout weights are expected to depend on the task sensitivities of the readout neurons – i.e., how well those neurons discriminate between stimulus categories in a given task (Fig. 2). In the case of an optimal linear decoder, a neuron’s CP becomes proportional to its own sensitivity (Haefner et al., 2013). But even when the decoder is suboptimal or nonlinear, some degree of alignment between sensitivity and readout weights is predicted provided that decoder performance remains substantially above chance (Haefner et al., 2013).

In models incorporating feedback, the dependencies of CP on readout weights, correlations, and sensitivity remain qualitatively similar to those in feedforward models as long as feedback signals are approximately aligned with the feedforward read-out (Chicharro et al., 2021). While the assumption of alignment between feedback and readout has been called into question by empirical studies (Zhao et al., 2020; Levi et al., 2023; Quinn et al., 2021), as long as there is some degree of alignment, a similar – though likely attenuated – relationship between CP and sensitivity can be expected.

In a hierarchical Bayesian inference framework, CPs arise from dependencies in the joint posterior over latent variables. In the context of a perceptual decision-making task, sensory neurons encoding latent statistical features of the inputs (Olshausen and Field, 1996) become correlated with higher-order latent variables representing choices or actions (Langlois et al., 2025). These dependencies are the result of feedforward signals that reflect bottom-up sensory evidence and feedback signals reflecting prior beliefs (Haefner et al., 2016; Lange and Haefner, 2022; Liu et al., 2026). Variability in these top-down belief signals – shared among sensory neurons – generates covariability in their responses that, after learning, aligns with the task structure, leading to the emergence of CPs and their relationship with neural sensitivity (Lange and Haefner, 2022). Importantly, Equation (1) still holds for the hierarchical models when evaluated using empirically estimated effective read-out weights, *w_j_*, justifying the organization of factors shown in Fig. 2 (Lange and Haefner, 2022).

Empirical evidence indicates that real neural communication is far more complex than the simplified interactions proposed by current theoretical models. Recent large-scale recordings in rodents have revealed rich, brain-wide dynamics during perceptual decision-making, and there is little reason to believe that these dynamics are any simpler in primates. These processes involve intricate interactions between excitatory and inhibitory sub-networks (Najafi et al., 2020; Esmaeili et al., 2022; Kuan et al., 2024), giving rise to synchronized activity and non-trivial temporal evolution of shared signals—particularly choice-related signals across multiple brain areas (Steinmetz et al., 2019; Ebrahimi et al., 2022; Orlandi et al., 2023; Bondy et al., 2025; Angelaki et al., 2025). While such complexity complicates the interpretation of summary statistics like choice probability (CP), it does not undermine their utility. However, the precise interpretation of CP – especially with respect to causal contributions – depends critically on the underlying modeling assumptions, such as whether one adopts a feedforward encoding–decoding perspective or a probabilistic inference framework.

In practice, neither the full structure of neural covariability nor the readout mechanism is directly observable. However, they differ in how accessible they are to empirical investigation. Correlations among neurons can be partially estimated from multi-electrode recordings, but such measurements capture only a small – and often biased – subset of the broader population. Most CP studies, however, rely on single-electrode recordings heavily restricting the estimation of inter-neuronal correlations. In contrast, the readout mechanism is fundamentally unobservable: because readout weights aggregate the effects of multiple synapses and entire neural circuits, they cannot be measured directly and must instead be indirectly inferred by modeling the relationship between neural responses and behavior.

However, several experimental variables that are expected to influence both neural correlations and readout weights are directly observable (indicated as gray in Fig. 2). For instance, many CP studies assess the task sensitivities of neurons, which are expected to strongly shape the readout profile in the case of optimal decoding or inference (Section 2.4). The recorded brain area – and its position within the cortical hierarchy – may influence the strength of its feedforward contribution to the decision and the strength of shared feedback modulation (Section 2.5). Other relevant factors – such as decision confidence, task engagement, and the extent of task or perceptual learning – are not directly observed, but can be inferred from behavioral measures or task design features (Fig. 2).

Fig. 2 offers a simplified conceptual overview of the key dimensions along which experimental and analytical choices are expected to influence CP values (for a more complete overview see Fig. S1). The arrows represent hypothesized relationships informed by theoretical considerations, some of which are supported by partial, sometimes controversial, empirical evidence (see Tables 2 and 3 for details). Two central variables in this diagram have direct links to Equation (1): (1) the properties of the decision decoder and its readout profile (Decoder type/Read out **w**) and (2) inter-neuronal correlations (Covariability **C**). Most decisions made in the design of experiments ultimately affect CP values through their influence on these central factors, while other potential pathways involve saccade planning or confounding biases (Fig. S1). Our meta-analysis examines the evidence for or against these hypothesized relationships – represented by the arrows in Fig. 2 – using observed experimental variables whenever possible.

### 2.2 Approach

We compiled a complete^1^ dataset of CP values and associated experimental variables to examine their influence on CP. We focused on sensory areas of the macaque monkey visual cortex, generally excluding sensory-motor regions (e.g. VIP and LIP) where motor-related signals, with their direct link to choice, are more prevalent (Zaidel et al., 2017; Katz et al., 2016; Yu and Gu, 2018) (see Section 7.1). When feasible, we analyzed data on a per-monkey basis – typically two monkeys per study – although in some cases only combined data were reported. In total, we reviewed 59 studies and extracted 150 mean CP values (some studies included more than two monkeys or tested multiple brain areas or tasks) spanning studies from 1994 until 2024. First, we extracted information from published studies reporting choice probabilities and then contacted authors to obtain missing details. We used the mean CP across the entire recorded population as our primary metric, as it is the most commonly reported measure across studies. Wherever possible, we used CP values computed with the most widely adopted estimation method (grand CP, stimuli with random noise; see Section 8.3), and for the 43% of data points where this method was not reported, we selected the most similar available alternative (see Section 7). While some studies have raised concerns regarding potential confounds in this widely adopted metric (Nienborg and Cumming, 2009; Kang and Maunsell, 2012; Wimmer et al., 2015), we found no evidence that the choice of estimation method significantly influenced our results (see Section 8.3).

We performed regressions between mean CP values and each experimental variable to test the hypothesized relationships. We adopted a nested approach, beginning with neuronal sensitivity as the primary predictor of CP. Subsequent analyses stepwise incorporated additional experimental dimensions. This strategy allowed us to prioritize more fundamental predictors while systematically assessing the contribution of more complex or indirect factors.

### 2.3 Data overview

CP studies have reported substantial variability in results. The average mean CP across all data points was 0.55, with substantial variability (95% central range: 0.47-0.66, *N* = 150). Of the 124 mean CP values for which statistical significance was reported, 87 (68%) were significantly above 0.5, consistent with theoretical expectations, while three were significantly below (Goris et al., 2017; Levi et al., 2023), in contradiction to any existing model. We also observed substantial variability in CP values between monkeys from the very same study, calculated separately within each study × brain area × task combination (Fig. S2). The average difference was 0.03 (median: 0.02; inter-quartile range: 0.01-0.04) – amounting to 58% of the average difference between any two points in the dataset (Fig. S2).

Interestingly, the published mean CP values declined across years of publication, reminiscent of the well-known “decline effect” observed across scientific disciplines (Schooler, 2011; Pietschnig et al., 2019). We found a pronounced negative relationship between mean CP and year of publication (slope: *β* = −0.0019, *p <* 0.001; Fig. 3a). This trend was evident across most brain areas (Fig. 3a). It was only partly attributable to the growing use of multi-electrode recordings – which typically yield lower CP estimates – as the decline effect persisted even when limiting the analysis to single-electrode ^2^ studies (Fig. S3). The decline effect is commonly attributed to publication bias, selective reporting, changes in experimental rigor or methodological changes (Schooler, 2011; Pietschnig et al., 2019). We did observe that studies with lower statistical power tend to report higher CP values, consistent with potential publication bias (Fig. S4). However, when controlling for changes in neuronal sensitivity, brain area, stimulus duration, and task type, the effects of both publication year and statistical power disappeared (Fig. 3b; Supplement 8.4). In fact, the sensitivity of recorded neurons contributed most strongly to explaining both the decline effect and the relationship to statistical power as neural sensitivity decreased with year of publication (Fig. 3c) while sample sizes increased (Fig. S27). These results support an alternative explanation for the decline in CP values: methodological shifts toward a less effective tailoring of the stimulus to the recorded neurons.

**Figure 3:**
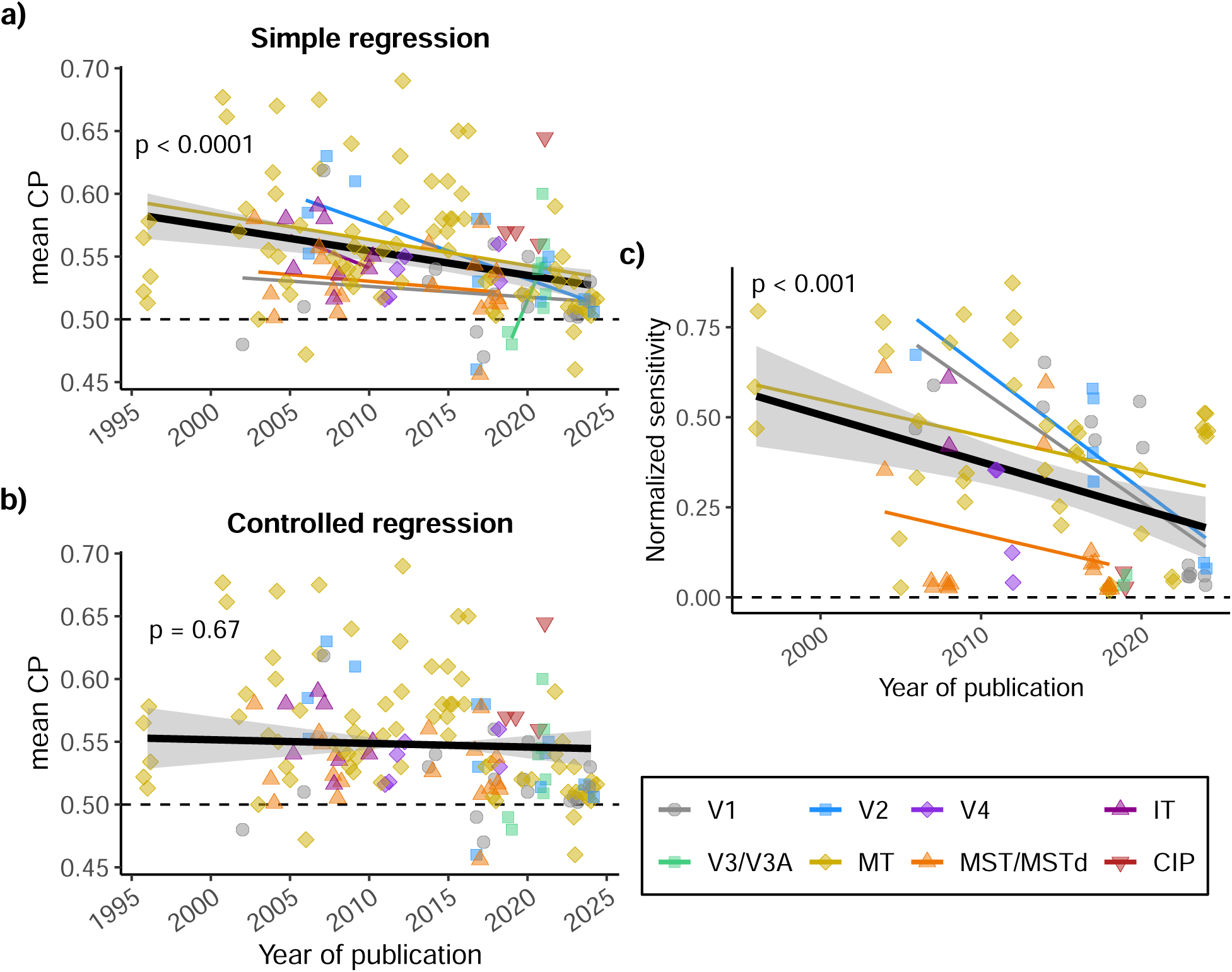
Mean CP values decline over time, primarily due to reduced neuronal sensitivity in later studies. **(a)** Relationship between mean CP and year of publication. Each point represents a study-level mean (typically per monkey, brain area, and task). Points are slightly jittered horizontally for visibility. The thick black line shows the overall regression (*β* = −0.0019, *p <* 0.001, *N* = 150); colored lines show within-area regressions. **(b)** Same as (a), but showing the partial regression after controlling for neuronal sensitivity, brain area, stimulus duration, and task type. No significant relationship remains, regardless of how we control for sensitivity (either via stimulus tailoring, black line, *p* = 0.68, or the inverse N/P ratio, not shown, *p* = 0.55, also see Section 2.4). **(c)** Relationship between normalized neuronal sensitivity and year of publication. Sensitivity is approximated by the inverse of the mean neurometric-to-psychometric threshold ratio (inverse N/P ratio; see Section 2.4.1). A strong decrease in sensitivity is observed, likely reflecting methodological shifts toward less stimulus tailoring to individual neurons.

In the following sections, we examine how mean CP depends on the experimental variables outlined in Fig. 2.

### 2.4 Sensitivity of the recorded neurons to the task

#### 2.4.1 Measures of neuronal sensitivity

Comparing neuronal sensitivity across studies is difficult because standard measures depend on experiment-specific stimulus units (e.g. % coherence). To address this, we normalized neural sensitivity by the monkey’s behavioral performance, providing a common reference across studies. A second issue is that the commonly reported neurometric-to-psychometric threshold ratio (N/P) is not well suited for testing theoretical predictions as both feedforward and feedback frameworks predict a *linear* relationship between CP and sensitivity (Haefner et al., 2013; Pitkow et al., 2015; Haefner et al., 2016; Chicharro et al., 2021), translating into a *nonlinear* one between CP and threshold (Fig. 4a). We therefore use the psychometric-to-neurometric ratio (P/N) as a sensitivity measure. Because most studies do not report P/N, we approximate it as the inverse of the mean N/P ratio, noting that this underestimates the true P/N (see Supplement 8.6). Finally, to account for differences in how neurometric thresholds were computed, we adjusted reported values to standardize mean N/P ratios across studies (see Section 7).

**Figure 4:**
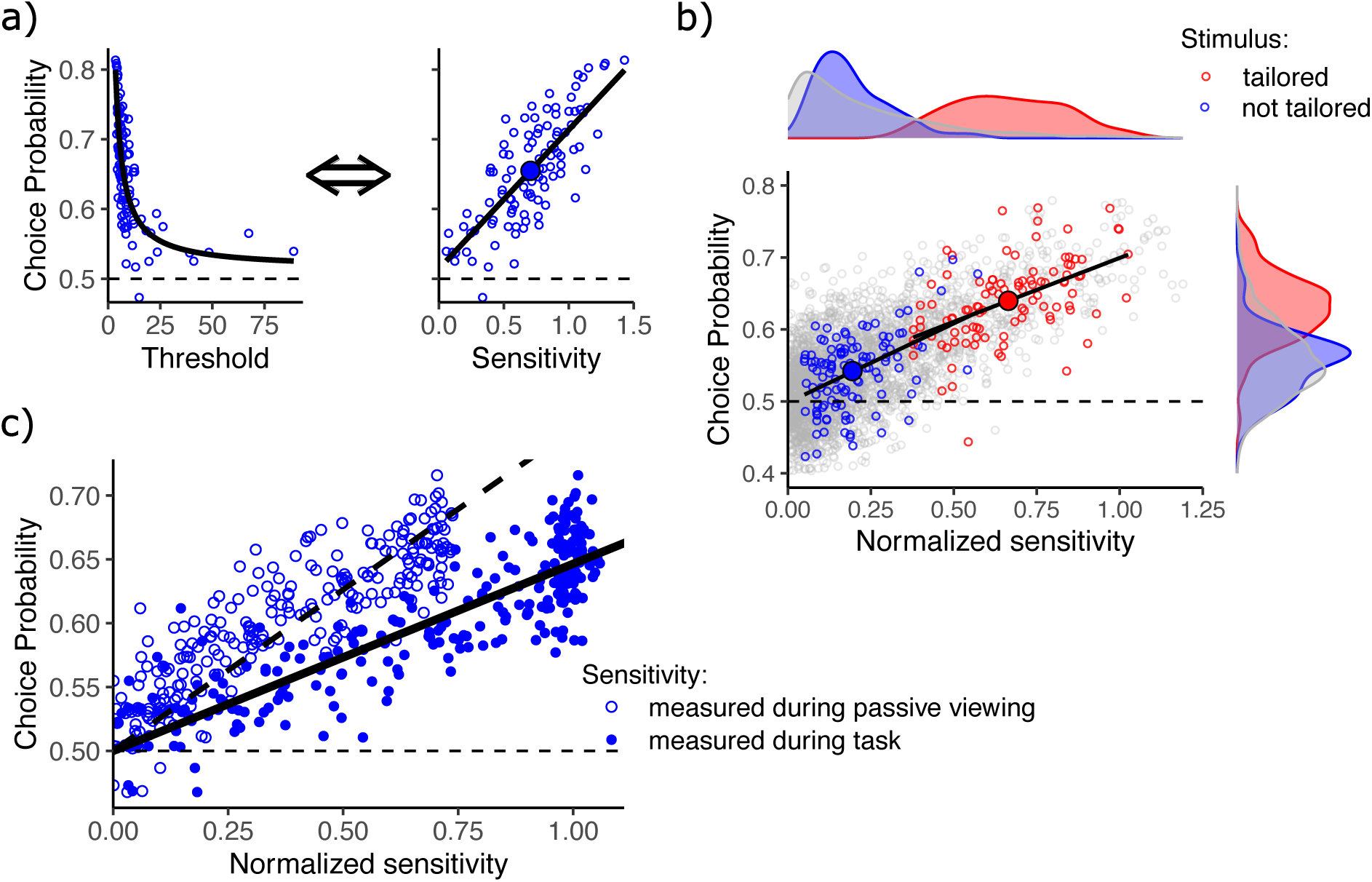
Choice of sensitivity metric and sampling strategy shapes the CP–sensitivity relationship. All panels present simulated data. **(a)** Toy simulation showing how neuronal informativeness metrics yield distinct relationships with CP. Left: CP vs. neurometric threshold. Right: CP vs. sensitivity (inverse threshold). Open circles are simulated neurons; large filled circles are study averages. Solid lines denote ground truth relationship. Theoretical models predict a linear CP–sensitivity relationship, but a hyperbolic relationship with threshold. **(b)** Toy simulation illustrating how stimulus tailoring impacts sampled sensitivity and CP. Grey points represent the full hypothetical neuronal population. Red points denote a simulated study tailoring stimuli to individual neurons (sampling the high-sensitivity tail); blue points denote a study without tailoring (random sample with a minimum response threshold). Despite sampling differences, both studies yield similar CP–sensitivity slopes (solid black lines) that closely match the line connecting their averages (large colored circles). Insets: marginal distributions. **(c)** Task-aligned feedback inflates sensitivity estimates. This effect is illustrated using simulations from the hierarchical inference model of Haefner et al. (2016). Normalized sensitivity is estimated either without feedback (resembling passive viewing data, open circles), or with feedback (resembling active task data, filled circles). CP values remain identical across conditions. However, feedback elevates estimated sensitivity—especially for already-sensitive neurons (rightward shift)—causing the CP–sensitivity slope to become shallower (solid vs. dashed line).

Variability in mean sensitivity across studies is largely explained by the degree to which stimuli were tailored to the tuning properties of recorded neurons. While it is standard practice to tailor stimulus parameters to maximize neuronal sensitivity to the task, studies vary considerably in how this tailoring is applied. We graded the degree of stimulus tailoring across three parameter categories: (1) stimulus size, (2) task-relevant parameter (e.g., orientation in orientation discrimination tasks), and (3) task-irrelevant parameters that still affect neuronal responsiveness (e.g., spatial frequency in orientation discrimination tasks). Together, these three tailoring factors accounted for approximately 57% of the variance in the inverse mean N/P ratio (*R*^2^_adj_ = 0.57), with significant contributions from task-relevant parameter and stimulus size (Fig. S5a and b). Differences in tailoring techniques across studies can be viewed as different sampling strategies: some preferentially target neurons from the high-sensitivity end of the distribution, whereas others approximate a random draw from the underlying population, which is dominated by lower-sensitivity neurons (Fig. 4b). Feedback signals can systematically increase sensitivity estimates in CP studies, when the feedback aligns with the stimulus. In standard practice, neuronal preferences (which stimulus is preferred) are measured during separate passive viewing sessions that minimize choice-related feedback, whereas sensitivities (magnitude of response difference) are typically estimated directly from task trials where responses likely reflect choice-related top-down modulation. To our knowledge all CP studies in our dataset that estimated N/P ratios followed this approach. This can affect sensitivity estimates in a systematic way (Zaidel et al., 2017; Liu et al., 2026). In models incorporating feedback, task-aligned top-down signals are expected to increase estimated sensitivity across almost all neurons they target (Fig. 4c, Lange and Haefner (2022); Liu et al. (2026)).

#### 2.4.2 Empirical relationship between CP and neuronal sensitivity within and across studies

Empirical studies generally support the theoretical prediction of a positive relationship between CP and neuronal sensitivity. Of the 49 data points (spanning 29 studies) that explicitly tested this link, 25 (51%) found a significant relationship in the theoretically predicted direc-tion—specifically, a negative correlation between CPs and neural thresholds and a positive correlation between CPs and sensitivities. Only one experiment – in V2 of a single monkey in Goris et al. (2017) – found the opposite effect, with a positive CP–threshold correlation (this same data point also had a mean CP significantly below 0.5). Across studies, the median correlation coefficient was 0.27 (interquartile range, (IQR): 0.12–0.37, N = 43). Intuitively, tighter CP–sensitivity relationships should correspond to higher mean CP values. Indeed, studies reporting stronger CP–sensitivity correlations also tended to report higher mean CP values: a 0.1 increase in the correlation coefficient was associated with a 0.01 increase in mean CP (*β* = 0.10, *p* = 0.001, *R*^2^_*adj*_ = 0.21; Fig. S6).

We next asked whether this relationship also holds across studies—that is, whether the mean CP of a sampled population correlates with its mean sensitivity across different cortical areas, tasks, and subjects. Importantly, such an across-study correlation does not necessarily follow from the observed within-study relationship. Two limiting scenarios can be considered (Fig. 5a). At one extreme (left panel), the slope of the mean CP–mean sensitivity relationship across studies matches the average of the within-study slopes. At the other extreme (right panel), no relationship is observed across study means. The first scenario would suggest a common underlying mechanism linking CP and sensitivity at both within-study and across-study levels, for example if differences in mean sensitivity across studies reflected only differences in neuronal sampling (Fig.4b). The second scenario would indicate that variability across neurons and across experiments arises from different sources. For instance, across-study differences in mean sensitivity may stem from the idiosyncrasies of individual subjects or lab-specific methodological nuances in recording and analysis. To distinguish between these scenarios, we estimated both the within- and across-study CP–sensitivity slopes using the P/N ratio as a standardized measure of neuronal sensitivity.

**Figure 5:**
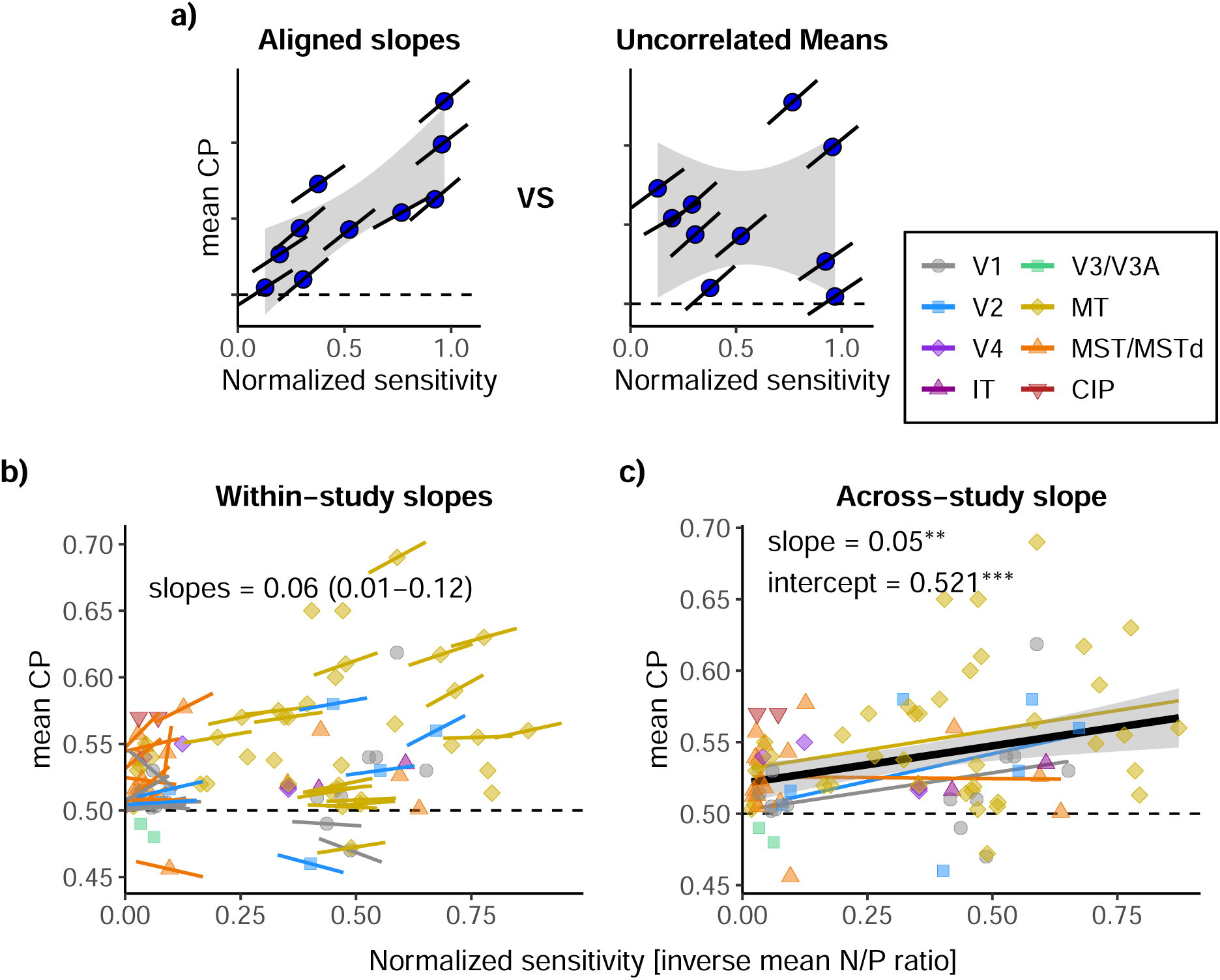
Positive relationship between CP and sensitivity both within and across studies, with a positive offset across studies. **(a)** Conceptual illustration of two edge cases. Left: the slope of the across-study mean CP–mean sensitivity relationship closely matches the average within-study slopes, suggesting a shared mechanism. Right: no relationship across studies, implying different sources of variability. Circles represent study-level means; shaded gray region – schematic 95% confidence interval of the across-study relationship; short thin lines – within-study slopes. **(b)** Empirical mean CP–mean sensitivity relationship across studies. Each point is a study-level mean. The x-axis shows normalized sensitivity approximated by the inverse of the mean neurometric-to-psychometric threshold ratio (inverse N/P ratio). Short colored segments passing through the points denote within-study slopes for the subset of studies where data were available (*N* = 42). The text insert reports the median and interquartile range (25%–75%) of these within-study slopes: 0.06 (0.01-0.12). **(c)** Same data as in (b), but overlaid with across-study relationships. The thick black line represents the across-study relationship with sensitivity as the sole predictor with slope of 0.05 (*p* = 0.003, *N* = 88). A multivariate model incorporating brain area, stimulus duration, and task type yielded a similar sensitivity slope of 0.04 (not shown; *p* = 0.02, *N* = 88). Thin colored lines show regressions within brain areas with at least 6 data points. Notably, the across-study slope estimate falls within the distribution of the within-study slopes.

We could extract within-study CP–P/N ratio slopes from 14 studies (yielding 42 data points). Notably, with the exception of Clery et al. (2017) and Nienborg and Cumming (2006), these slopes were not explicitly reported in the original publications and had to be estimated de novo from neuron-level data (see Methods). The slopes were predominantly positive, albeit with considerable variability in magnitude: 31% (*N* = 13) were significantly positive, whereas only two data points (from Goris et al. (2017) and Lange et al. (2023)) were significantly negative (Fig. 5b). The median (IQR) of the within-study slopes was 0.06 (0.01–0.12, *N* = 42).

We also found a positive relationship between mean CP and mean task sensitivity across studies (Fig. 5c). The slope of this relationship was 0.05 (*p* = 0.003, *N* = 88) and normalized sensitivity explains 10% of mean CP variance (*R*^2^_*adj*_ = 0.10). The relationship persisted after controlling for brain area, stimulus duration, and task type (*β*_sensitivity_ = 0.04, *p* = 0.02, *N* = 88). Importantly, the magnitude of this across-study slope is highly consistent with the within-study data, lying close to the median and well within the interquartile range. The mean CP – mean sensitivity relationship was not consistently observed within individual brain areas – possibly due to limited sample sizes (see Fig. 5c and Section 2.5).

The comparable magnitude of the within- and across-study CP–sensitivity slopes is consistent with a shared underlying mechanism linking CP and sensitivity at both the single-neuron and study levels (Fig. 5a, left panel). Our finding that variability in mean sensitivities across studies is largely explained by differences in tailoring technique (section 2.4.1) –a plausible proxy for neuronal sampling – further supports this interpretation.

We found that the intercept of the across-study CP-sensitivity regression was significantly greater than 0.5 – both when controlling for sensitivity alone (*β*_0_ = 0.521, *p <* 0.001, Fig. 5c) and within MT after additionally controlling for stimulus duration and task type (*β*_0_ = 0.512, *p <* 0.001). Similarly, the intercepts of the within-study CP–sensitivity relationships were skewed above 0.5, with a median of 0.51 (IQR: 0.49–0.53). From a theoretical standpoint, such offset is unexpected and suggests that even neurons with minimal task sensitivity still exhibit robust choice-related activity *aligned* with their preferences.

To ensure we did not discard 41% of our dataset lacking inverse mean N/P ratio data (*N* = 62*/*150), we conducted two analyses in parallel: one using the N/P ratios where available, and another one using the degree of stimulus tailoring (see Section 2.4.1) to approximate sensitivity across the entire dataset. Tailoring technique explained mean CP to a similar extent as the P/N ratio (*R*^2^_adj_ = 0.07, Fig. S5c and d). Thus, tailoring exhibits systematic relationships with both the inverse mean N/P ratio and mean CP. These findings support the use of tailoring technique as a coarse but practical proxy for neuronal sensitivity, enabling us to include the full dataset in analyses that control for sensitivity. In the following sections, we use two alternative regression models to control for sensitivity: one using the inverse mean N/P ratio (available for a subset of the data), and one using the three variables representing degree of tailoring: for stimulus size, for task-relevant parameter and for task-irrelevant parameters (applicable to the full dataset).

In conclusion, we observed a similar CP–sensitivity relationship at the single-neuron and at the per study level in agreement with theoretical models.

### 2.5 Brain area

Whether a neuron’s position in the visual hierarchy influences its CP depends on how directly it is engaged by the decision process – either as a feedforward input to the decision area or as a target of decision-related feedback. Two competing hypotheses make different predictions (Fig. 6a).

**Figure 6:**
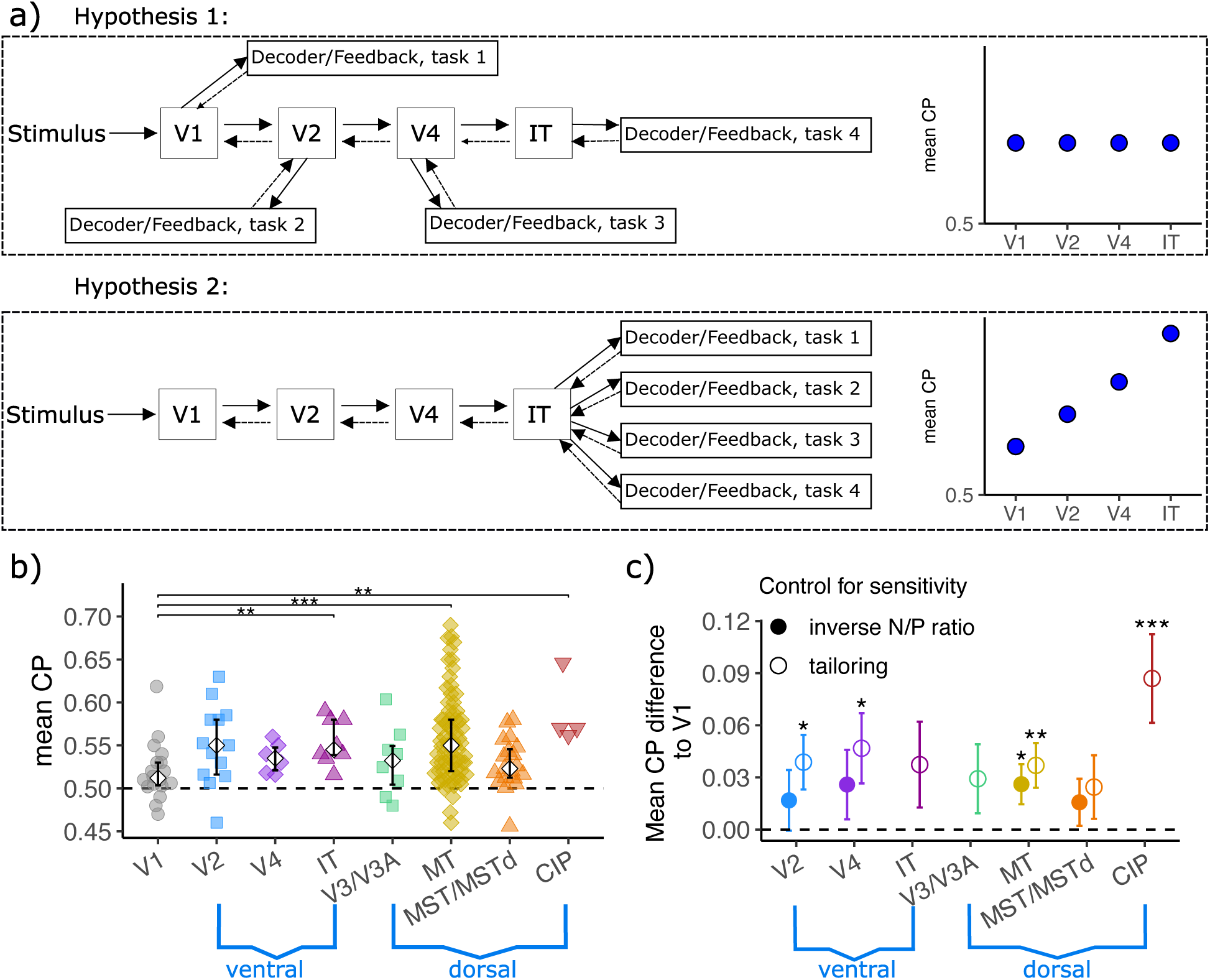
Lowest mean CP in V1 but no consistent increase with hierarchical level across brain areas. **(a)** Schematic illustration of two competing hypotheses about how CP varies across visual areas. Top: the decoder or feedback can target any sensory area, resulting in comparable CP values across areas if tasks are well matched to the recorded neurons. Bottom: the decoder or feedback accesses only the most downstream area, leading to increasing CP with hierarchical level. **(b)** Mean CP values across brain areas organized by hierarchical position across the ventral and dorsal streams. We assigned area V3/V3A to the dorsal stream, although the classification of its constituent parts remains a subject of ongoing debate (Felleman and Van Essen, 1991; Lyon and Kaas, 2002; Kravitz et al., 2011; Rosenberg et al., 2023). Diamonds and error bars represent medians and interquartile ranges for areas with more than five observations. Asterisks indicate significant differences between two brain areas based on Mann–Whitney tests (** *p <* 0.01, *** *p <* 0.001); *p <* 0.05 results are omitted as they would not survive correction for multiple comparisons. **(c)** Regression coefficients capturing the contribution of each brain area to mean CP, controlling for sensitivity (either inverse mean N/P ratio or tailoring technique), stimulus duration, and task type. V1 served as the baseline category and is not shown. Filled and open circles show regression coefficients using, respectively, the inverse N/P ratio and tailoring technique to control for sensitivity. Error bars indicate standard errors. Coefficients are only shown for brain areas with at least four data points. Asterisks denote significance relative to V1 (*V3/V3A (N = 8), MT (N = 74), MST/MSTd (N = 19), and CIP (N = 4). In our dataset, some areas—particularly IT, MT, and CIP, exhibited higher mean CPs than V1; however, beyond that, we found no evidence that later stages of the visual hierarchy are associated with higher CPs (Fig. 6b). Although a subset analysis of single-electrode recordings showed a similar overall pattern, it unexpectedly revealed lower mean CP values for MST compared to MT, despite MST occupying a higher level in the dorsal stream (Fig. S7). This surprising result can be explained by the lower task sensitivity in MST compared to MT (Fig. S9). Notably, V1 and MT had similar sensitivity, so the CP difference between them cannot be attributed to this factor.

Under the **flexible access hypothesis**, the decision system recruits whichever area is most informative for the current task, e.g. MT for motion discrimination, V1 for orientation discrimination, regardless of hierarchical level. Because feedforward and feedback connections link virtually all visual areas to higher-order regions (Kravitz et al., 2011, 2013), the decision area can read out from, and send feedback to, any sensory area directly. Since CP studies are typically designed to match the task to the recorded area, this view predicts no consistent relationship between CP and hierarchical position (Fig. 6a, top).

Under the **restricted access hypothesis**, the decision area draws input only from the most downstream sensory regions, in line with classic feedforward architectures. Consequently, CP is expected to increase monotonically along the sensory hierarchy: higher-order areas directly read out by decision circuits exhibit the strongest choice signals, while the choice signal in individual upstream neurons is progressively diluted. The same pattern is predicted for feedback contribution to CP: progressively weaker CP due to diminished top-down influences in lower sensory areas (Fig. 6a, bottom), in agreement with attention studies which typically find weaker effects in lower sensory areas Maunsell (2015).

Our meta-analysis revealed lower mean CP values for V1 – the first stage of the visual cortical hierarchy – but no consistent trend across higher areas (Fig. 6b,c). Visual processing in primates is classically divided into two cortical streams: the ventral “what” stream for object recognition and form processing, and the dorsal “where/how” stream for spatial and motion processing (Mishkin et al., 1983). Based on anatomical and functional studies in macaques, we grouped visual areas accordingly, though the hierarchical placement of some areas remains debated (Markov et al., 2014). V1 (N = 18), which feeds into both streams, sits at the base of the hierarchy and served as the baseline in our regression models. The ventral stream includes V2 (N = 13), V4 (N = 6), and IT (N = 8); the dorsal stream includes

Regression analyses controlling for task sensitivity, stimulus duration, and task type confirmed the main findings. We used V1 as the baseline, so the reported regression coefficients (*β*) represent the mean CP difference relative to V1 (Fig. 6c). MT showed significantly higher mean CP than V1, by about 0.03–0.04 on average across both regression models (inverse mean N/P ratio as sensitivity control: *β_MT_* = 0.03*, p* = 0.03; tailoring: *β_MT_* = 0.04*, p* = 0.005; Fig.6c). For other areas, conclusions were limited to the model using tailoring for the sensitivity control, as the small number of N/P ratio data points made coefficient estimates in the alternative model too uncertain (Fig. S9). In this model, three additional areas – V2 (*β_V_* _2_ = 0.04, *p* = 0.02), V4 (*β_V_* _4_ = 0.05, *p* = 0.02) and CIP (*β_CIP_* = 0.09, *p <* 0.001) – also showed significantly higher mean CP than V1 (Fig. 6c). While prominent oculomotor signals in CIP likely inflated its mean CP, motor-related signals have been found to be no more prevalent in CIP than V3A, despite CIP exhibiting a distinctly higher CP (Doudlah et al., 2022; Zhu et al., 2024).

Only a few empirical CP studies have recorded from multiple brain areas within the same experiment, and their findings remain inconclusive. In our dataset, we identified nine such studies, each involving recordings from exactly two areas in the same animals and same experimental context. These limited data reveal a consistent increase in mean CP along the dorsal, but not the ventral stream hierarchy (Fig. S8). It is important to note, however, that these pairwise comparisons do not control for differences in task sensitivity across brain areas, as the same task may be more appropriate for one area than another. Indeed, in four of the six studies where sensitivity data were available, the area with higher mean CP also showed higher task sensitivity (Goris et al., 2017; Kang and Maunsell, 2020; Nienborg and Cumming, 2006; Yu and Gu, 2018).

In summary, our analysis reveals a clear pattern: V1 exhibits lower mean CP compared to other visual areas, while there is no evidence for a systematic increase in CP with hierarchical level beyond V1. This suggests that V1 may be functionally distinct – either less directly accessed by decision-related signals or receiving weaker feedback – while the lack of a hierarchical trend beyond V1 supports the idea that the decision system flexibly targets the most task-informative neurons, irrespective of cortical level. Although our dataset is dominated by MT, V1, and MST recordings – highlighting the need for more data from other areas – the distinctiveness of V1 is robust and consistent. These findings are compatible with computational theories proposing a special status for V1 in sensory processing and learning (Crick and Koch, 1995; Li, 2002; Lee and Mumford, 2003).

### 2.6 Stimulus duration

Stimulus presentation duration varied widely across studies in our dataset, ranging from as short as 67 ms to over 2 seconds—a difference of more than 30-fold. In the majority of studies (84% of data points), the trial structure required the monkey to view the stimulus for a fixed duration, after which it was allowed to make a choice. The remaining studies used a reaction time paradigm, in which the monkey could respond at any time, resulting in variable stimulus durations; in these cases, we used the reported average stimulus duration for our analysis. For CP calculations, researchers typically used spike counts over the full stimulus presentation window (85% of data points) – accounting for response latency and excluding any activity prior to the motor response.

While the within-trial time course of instantaneous CP (calculated using short temporal bins) can offer mechanistic clues, it does not necessarily predict how overall CP scales with stimulus duration across studies. Furthermore, empirical reports of within-trial CP dynamics are inconsistent. Some studies observe a gradual increase in CP over the course of a trial (Uka and DeAngelis, 2004; Nienborg and Cumming, 2006, 2009; Bondy et al., 2018), whereas others report an abrupt initial rise followed by a sustained plateau (Britten et al., 1996; Liu and Newsome, 2005; Price and Born, 2010; Shiozaki et al., 2012; Krug et al., 2016). Crucially, how these within-trial dynamics translate into a relationship with stimulus duration – and specifically, how a given study’s CP trajectory would adapt if the exposure time were experimentally manipulated – remains an open question.

Feedforward models predict either no relationship or a negative relationship between mean CP and stimulus duration, depending on how decoding efficiency changes with exposure time (Fig. 7a, blue lines). Feedforward integration-to-bound models, such as the drift diffusion model (DDM), posit that a decision area accumulates momentary sensory evidence (i.e., spike counts of sensory neurons) until a decision threshold is reached (Gold and Shadlen, 2007). Within this framework, the predicted dependency of CP on stimulus duration depends on specific model assumptions:

- If integration spans the entire stimulus duration, CP does not change with duration. Because overall behavioral performance is usually matched across studies by adjusting signal level, the informativeness of the stimulus per unit time is reduced with stimulus duration (Nienborg et al., 2012). Thus, a neuron’s fractional contribution to the decision remains constant regardless of duration.
- If, on average, the ratio of active integration time to total stimulus duration remains constant across task designs, the expected fraction of choice-influencing spikes is unchanged, thereby preserving the CP magnitude.
- As attention and memory demands scale with time, longer durations may cause the subject to conserve resources by ignoring some part of the evidence. This increases effective non-decision time and the proportion of non-contributing spikes, and as a result reduces mean CP. Ignoring early evidence aligns with the robust primacy effect observed in empirical studies, where early sensory evidence impacts choices more strongly than later inputs (Kiani et al., 2008; Nienborg and Cumming, 2009; Wyart et al., 2012; Lange et al., 2021; Levi et al., 2023; Liu et al., 2026), although at least some component of the primacy effect is likely due to feedback (Lange et al. (2021), also see below). On the other hand, leaky integrator models (Usher and McClelland, 2001) or potential conservation of effort early in a trial both generate a recency effect, but still predict a negative relationship between mean CP and stimulus duration.

**Figure 7:**
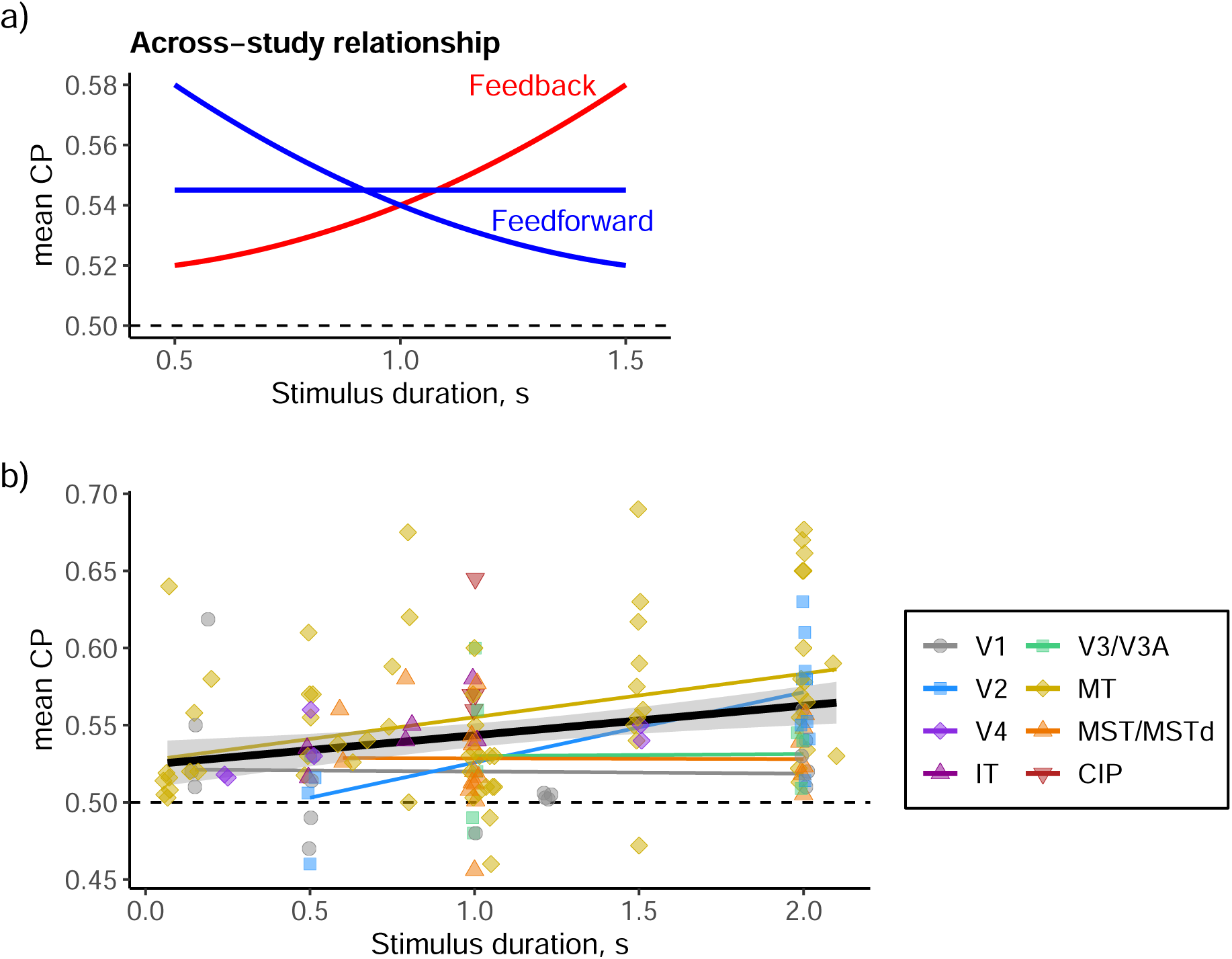
Mean CP increases with stimulus duration, aligning with feedback-based model predictions. **(a)** Schematic illustrating model predictions for the relationship between mean CP and stimulus duration. Feedforward models predict either no relationship (in classic integration-to-bound models) or a negative relationship (due to memory and attention constraints). Conversely, models incorporating feedback predict a positive relationship: hierarchical inference models predict this due to a larger number of samples acquired during inference, whereas models with post-decision feedback predict this due to increased post-decision time. **(b)** The empirical relationship between mean CP and stimulus duration across studies exhibits a positive slope in a simple linear regression with duration as the sole predictor (black line; slope = 0.019, *p* = 0.001). This positive trend remains robust in regressions controlling for sensitivity, brain area, and task type (not shown; inverse N/P ratio as sensitivity control: slope = 0.022, *p* = 0.001; tailoring as sensitivity control: slope = 0.019, *p* = 0.002). Data points are jittered horizontally for visibility. Thin colored lines show pairwise regressions within each brain area. Regression lines only shown for brain areas with more than 6 points.

Current feedback models predict a positive relationship between mean CP and stimulus duration (Fig. 7a, red lines):

- In hierarchical Bayesian models of approximate inference, the accumulation of decision-related beliefs generates feedback that progressively increases instantaneous CP over the course of a trial (Haefner et al., 2016; Lange and Haefner, 2022). Across different experimental designs of CP studies, longer stimulus durations typically entail a greater number of independent sensory samples (e.g., more stimulus frames) rather than simply stretching the duration of individual samples. This extended sampling period allows for prolonged belief accumulation and stronger overall feedback (Lange et al., 2021; Wen et al., 2022). In such a scenario these models predict a positive relationship between mean CP and stimulus duration across studies.
- Alternatively, an increase in the post-decision time, and hence, the task-aligned feedback that modulates sensory neurons after decision commitment could also drive a positive CP–duration relationship (Chicharro et al., 2017). Supporting the existence of such post-decision feedback, recent work has identified a discrete transition in neural population dynamics: a shift from an initial sensory-integration regime to a decision-commitment stage where the finalized choice is maintained in working memory (Charlton and Goris, 2024; Luo et al., 2025; Bondy et al., 2025).Choice signals are significantly more pronounced during this commitment phase than during the preceding integration stage (Levi et al., 2023; Charlton and Goris, 2024; Bondy et al., 2025). However, because some evidence suggests these post-decision signals may be misaligned with the neurons’ stimulus preferences (Zhao et al., 2020; Levi et al., 2023; Zahorodnii et al., 2025), whether the transition to a commitment regime definitively results in a higher CP remains an open question.

Our meta-analysis found a significant positive correlation between mean CP and stimulus duration, which remained robust after controlling for both neuronal sensitivity and other covariates (Fig. 7b). Regression analysis indicated that each additional second of stimulus presentation was associated with an increase of approximately 0.02 in mean CP both in a model with only stimulus duration (*β* = 0.019, *p* = 0.001), and after controlling for sensitivity, brain area and task type (inverse mean N/P ratio as sensitivity control: *β* = 0.022, *p* = 0.001; tailoring: *β* = 0.019, *p* = 0.002). The overall positive relationship between mean CP and stimulus duration was driven by MT and V2, the only areas in which the slope was individually statistically significant (model with stimulus duration only: MT *β* = 0.028, *p* = 0.003, *N* = 67; V2 *β* = 0.046, *p* = 0.007, *N* = 13; Fig. 7b). Excluding studies with reaction time tasks did not meaningfully alter any of the regression results. We also tested simple nonlinear transformations of stimulus duration (square root and logarithmic) to capture potential saturation effects; however, these transformations neither improved nor diminished the strength of the observed relationship.

This observed relationship cannot be explained by a simple increase in expected spike counts with stimulus duration. While the higher signal-to-noise ratio (SNR) for higher spike counts has been argued to increase CP measurements (Kang and Maunsell, 2012), this effect is likely too small to be observable for more than a few spikes per trial (Haefner et al., 2013). In line with this, Krug et al. (2016) empirically showed that manipulating task-irrelevant stimulus features to decrease a neuron’s firing rate by nearly half has no effect on CP estimates.

To conclude, the positive relationship between mean CP and stimulus duration provides some support for feedback-based accounts of the CP phenomenon. These results are consistent with mechanisms driven by either approximate hierarchical inference or post-decision choice signals. Notably, these two feedback-based models make divergent predictions for studies using reaction-time tasks: hierarchical inference continues to predict a positive relationship with stimulus duration, whereas models based on post-decision choice-related signals predict no such relationship, as these signals should be largely absent in reaction-time designs. Unfortunately, the small number of data points with reaction-time tasks in our dataset precludes a separate regression analysis to test these predictions directly (*N* = 15, model with only stimulus duration *p* = 0.65, Fig. S10).

### 2.7 Task type

We classified all tasks in CP studies into four categories: coarse-discrimination tasks, fine-discrimination tasks, detection tasks, and tasks involving bistable stimuli (Fig. 8a). In coarse-discrimination tasks, the two stimulus categories to be distinguished (e.g., orientation or direction of motion) are maximally separated – such as 90 degrees for orientation or 180 degrees for motion – but are made difficult to perceive due to added noise or low contrast. This is the most common paradigm in CP research, starting with Britten et al. (1996), and remains the most widely used (50% of the data points in our dataset). Fine-discrimination tasks (28% of data points) involve stimuli in which the task-relevant aspect is clearly visible, but the small difference between categories makes the task challenging, e.g. distinguishing between high contrast gratings of small orientation difference, or between high coherence motion stimuli with small differences in motion direction. Detection tasks (12% of data points) require the subject to report the presence or change of a specific feature in the stimulus and to withhold responses when no change or feature is present. While detection tasks vary in the amount of added noise and the magnitude of the feature change, we treat them here as a single group. The final category includes tasks involving bistable stimuli (10% of data points), where a single image gives rise to two distinct subjective percepts that alternate over time, typically on the order of a few seconds (Leopold and Logothetis, 1999; Sterzer et al., 2009; Krug et al., 2008). In contrast to the other task types, human experiments show that different choices to the same physical stimulus are associated with subjectively clear and distinct perceptual experiences (Brascamp et al., 2018).

**Figure 8:**
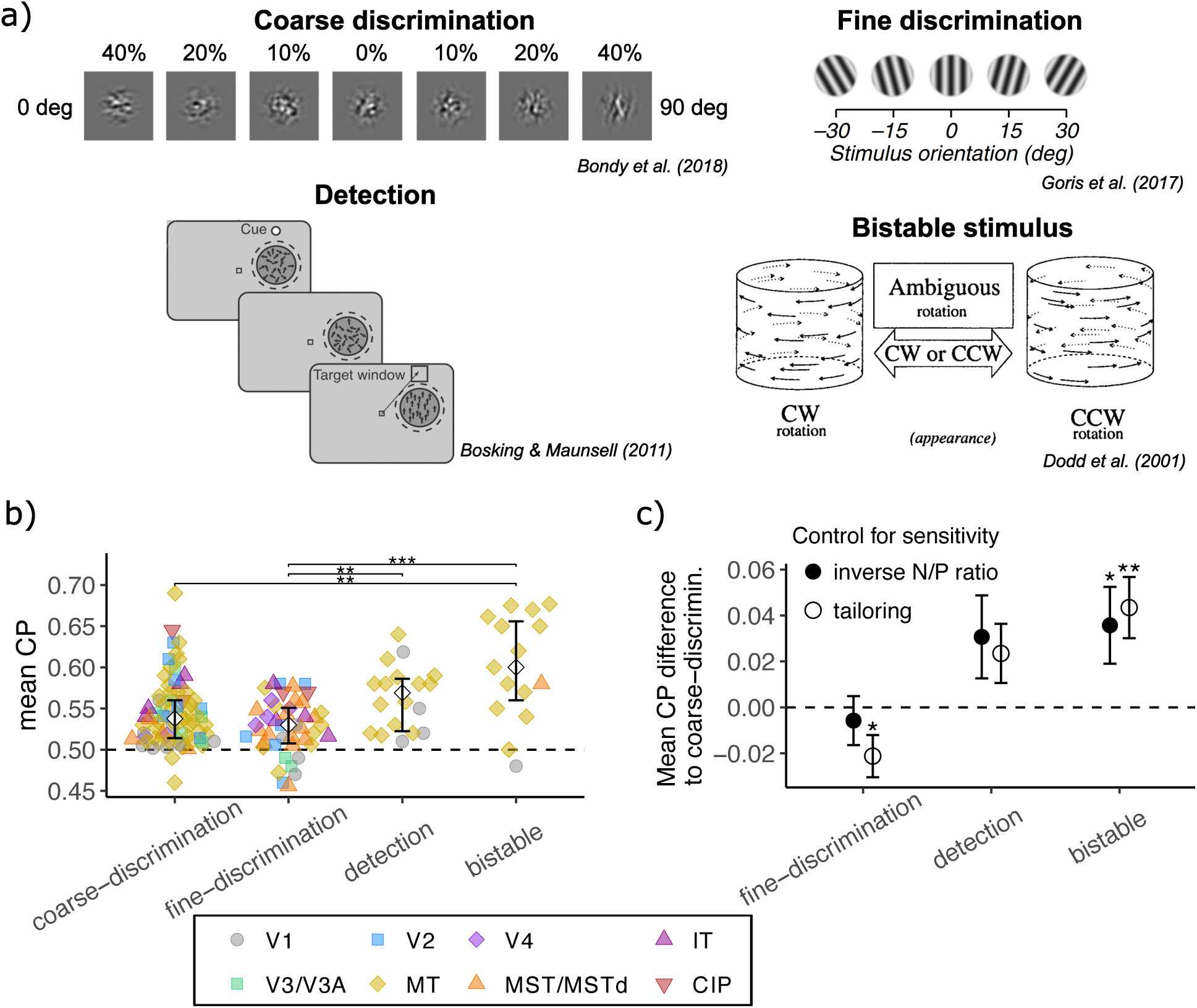
While mean CP varies systematically with task type, only bistable stimuli showed an independent effect. **(a)** Representative stimuli for coarse-discrimination, fine-discrimination, detection, and bistable stimulus tasks; in the latter, a single image evokes two distinct perceptions. **(b)** Mean CP values across four task categories. Diamonds and error bars represent medians and interquartile ranges. Asterisks indicate significant differences between two task types based on Mann–Whitney tests (** *p <* 0.01, *** *p <* 0.001); *p <* 0.05 results are omitted as they would not survive correction for multiple comparisons. While the difference in mean CP between coarse and fine-discrimination did not reach significance in the full dataset, it is significant within the subset of single-electrode studies (Fig. S13). **(c)** Regression coefficients estimating the effect of each task type on mean CP, controlling for sensitivity, brain area, and stimulus duration. Coarse-discrimination tasks serve as the baseline category and is not shown. Filled circles indicate coefficients from models using the inverse mean N/P ratio; open circles from models using tailoring technique. Error bars indicate standard errors. Asterisks denote statistical significance relative to coarse-discrimination tasks (* *p <* 0.05, ** *p <* 0.01). Only bistable tasks showed a consistently significant positive effect across both models. Conversely, the lower mean CP observed for fine discrimination under the tailoring control model is likely an artifact, as fine-discrimination tasks inherently exhibit smaller inverse *N/P* ratios than coarse-discrimination tasks even when stimulus tailoring is matched (see text).

Several individual studies have found that fine-discrimination tasks yield lower CPs and neuronal sensitivity than coarse-discrimination tasks even when targeting the same areas with similar stimuli (Purushothaman and Bradley, 2005; Uka and DeAngelis, 2006; Price and Born, 2010; Yu and Gu, 2018). Bistable stimuli, by contrast, tend to produce large CPs, most prominently in studies using a rotating cylinder stimulus in MT, where exceptionally high values were first reported by Dodd et al. (2001) and later replicated (Grunewald et al., 2002; Krug et al., 2004, 2016).

Our meta-analysis found that mean CP was lowest for fine-discrimination tasks and highest for bistable stimulus tasks (Fig. 8b and S13). Bistable tasks showed significantly higher CP than coarse and fine-discrimination tasks, which did not differ significantly from one another (Fig. 8b). Notably, when only comparing single-electrode studies, coarse-discrimination tasks showed higher mean CP than fine-discrimination tasks (Fig. S13). This suggests that the lack of a significant difference in mean CP between fine- and coarse-discrimination tasks (Fig. 8b) is driven by a higher proportion of multi-electrode studies in the coarse-discrimination group (41% vs. 14%) – a recording technique which yield lower CPs on average (Section 2.3).

Surprisingly, mean CP differences across task types were largely explained by sensitivity, brain area, and stimulus duration, with bistable stimuli being the only task type that still showed a consistently significant positive effect after accounting for these variables (Fig. 8c). We used coarse discrimination as the baseline, so the reported *β* for each task type represents the mean CP difference relative to coarse-discrimination tasks. Regression analyses showed no reliable difference for fine-discrimination tasks after controlling for brain area, stimulus duration and sensitivity using inverse mean N/P ratio (*β*_fine_ = − 0.006, *p* = 0.6; Fig.S11). Despite limited N/P ratio data (Fig. S15), bistable tasks showed significantly higher mean CP (*β*_bistable_ = 0.036, *p* = 0.04) while detection tasks did not (*β*_detection_ = 0.031, *p* = 0.09). Alternatively, when using tailoring as a proxy for sensitivity alongside brain area and duration covariates (Fig. 8c), the effect of bistable stimuli persisted (*β*_bistable_ = 0.043, *p* = 0.002) and detection remained non-significant (*β*_detection_ = 0.023, *p* = 0.07). Notably, fine discrimination showed a significantly lower mean CP under this model (*β*_fine_ = − 0.02, *p* = 0.02). However, this appears to be an artifact of the tailoring control, as fine-discrimination tasks inherently demonstrate smaller inverse N/P ratios than coarse-discrimination tasks even when tailoring is matched (see below).

#### 2.7.1 Fine-discrimination tasks

Arguably the most surprising finding comparing different tasks is the fact that the CP difference between coarse and fine-discrimination tasks can be entirely explained by the difference in neural sensitivity. Indeed, the median inverse N/P ratio in fine-discrimination tasks was several times smaller than for coarse tasks (0.08 compared to 0.47, Fig. S15). This large gap was partly explained by differences in tailoring technique (only for data points with available N/P ratios): stimulus parameters were substantially less often matched to neuronal tuning in fine-discrimination tasks – with task-relevant parameters not tailored in 71% of cases (vs. 23% in coarse), task-irrelevant parameters in 74% (vs. 18%), and stimulus size in 53% (vs. 8%). Yet even in the regression with controlling for tailoring and brain area, fine-discrimination tasks still showed an inverse N/P ratio that was on average 32% lower than in coarse-discrimination tasks (*p* = 0.007) with no other task type showing significant effects (detection: *p* = 0.4; bistable: *p* = 0.4).

It is unclear whether the low CP and low sensitivity observed in fine-discrimination tasks reflects systematic biases in experimental design or genuinely distinct processing mechanisms in the brain. We showed that stimuli in fine-discrimination tasks were less frequently tailored to neuronal tuning, possibly reflecting inherent difficulties in tailoring for such tasks. Because fine discrimination requires targeting the steepest slope of the tuning curve rather than its peak, this technically demanding process may leave greater mismatches between stimulus parameters and neuronal preferences, ultimately leading to reduced sensitivity and lower CP. Another possible bias could also occur if researchers have systematically targeted inappropriate brain areas specifically for fine-discrimination tasks. For example, it has been suggested that area MT is inherently less suited for fine discrimination due to its tuning properties (Purushothaman and Bradley, 2005; Uka and DeAngelis, 2006). Supporting this idea, one study found that MT microstimulation influenced choices in coarse, but not fine, depth-discrimination tasks (Uka and DeAngelis, 2006); however, another reported effects in both task types (Yu and Gu, 2018).

#### 2.7.2 Detection tasks

Although detection tasks exhibit a trend toward higher mean CP than coarse-discrimination tasks (*p* = 0.04, Fig. 8b), our regression analyses indicate that it is largely explained by differences in tailoring techniques. Studies employing detection tasks more frequently tailored stimuli to individual neurons, in part because they all used single-electrode recordings (100% vs. 59% for coarse discrimination). The mean CP difference between detection and coarse discrimination completely loses significance when compared within the single-electrode subset (*p* = 0.5, Fig. S13). Studies using detection tasks also disproportionately targeted MT (78% vs. 55% for coarse discrimination) but used shorter stimulus durations, partially offsetting the MT effect (Fig. S14). After controlling for sensitivity, brain area and duration, the difference in mean CP between detection and coarse-discrimination tasks largely disappears (Fig. 8c). This is surprising since there are substantial differences between detection and discrimination tasks that may influence CP. First, in most detection tasks, nearly all neurons prefer the same stimulus condition (stimulus presence or change), so unspecific global fluctuations, e.g. due to changes in alertness or eye movements, are expected to impact CP estimates for the whole neuronal population more than for other task types (Nienborg et al., 2012). Second, detection tasks in our dataset are strongly associated with reaction-time (RT) paradigms – 89% of detection tasks are RT tasks, and 67% of RT tasks are detection ones (Fig.S12). In feedforward models, this should enhance signal-to-noise ratio and increase CP (see Section 2.6). By contrast, models incorporating post-decision choice-related signals predict reduced CP in RT tasks due to diminished post-decision feedback. Our analysis leaves open the question whether these factors do not apply or cancel out.

#### 2.7.3 Bistable stimulus tasks

The most plausible explanation for the elevated CP values observed in bistable stimulus tasks is stronger collective neural dynamics, either driven by lateral or feedback signals, and associated with more vivid, and distinct subjective percepts (Brascamp et al., 2018). Initial efforts to explain high CPs in bistable tasks relied on winner-take-all feedforward models (Taylor and Aldridge, 1974; Dodd et al., 2001). However, without invoking top-down mechanisms or recurrency these models cannot adequately account for why some ambiguous stimuli evoke bistable perception (and high CP) while others do not (Leopold and Logothetis, 1999; Moreno-Bote et al., 2011; Dieter et al., 2016; Wasmuht et al., 2019). In humans, bistable perception recruits a broader network of higher-order brain regions and is accompanied by markedly stronger top-down and bottom-up influences (Wang et al., 2013; Lumer et al., 1998; Sterzer and Kleinschmidt, 2007; Weilnhammer et al., 2013; Brascamp et al., 2015). Attention-based fluctuations have been proposed to explain perceptual instability in binocular rivalry, where conflicting images are presented to each eye (Leopold and Logothetis, 1999; Dieter et al., 2016). Yet, only one study in our dataset (13% of data points of bistable stimulus) used a binocular rivalry paradigm (Maier et al., 2007). A key distinction between bistable and typical discrimination tasks is the subjective clarity of perception (Brascamp et al., 2018). Psychophysical studies in humans demonstrate that during a 1–2 second trial, the perceived direction of rotation of a bistable cylinder remains highly stable; spontaneous perceptual reversals generally occur only on the scale of tens of seconds (Krug et al., 2008). As in humans, the monkey in bistable tasks presumably experiences a vivid perceptual state even when the stimulus is ambiguous to an ideal observer, whereas typical CP tasks often involve low-confidence choices with weak sensory evidence. According to hierarchical inference models, decision confidence modulates the strength of belief signals fed back to sensory areas, thereby influencing CP (Haefner et al., 2016; Lange and Haefner, 2022). Hierarchical inference models thus predict that higher-confidence perceptual decisions – as might be expected in bistable tasks (Bainbridge et al., 2015) – should result in elevated mean CP. Wasmuht et al. (2019) demonstrated that bistable cylinders specifically increase long-timescale interneuronal correlations in MT, and that this network property directly predicts the elevated CP. While hierarchical inference models interpret such top-down influences as belief-induced correlations, Wasmuht et al. (2019) hypothesized that the feedback serves the functional role of stabilizing the dominant percept of the bistable stimulus.

Notably, 60% of bistable data points in our dataset came from studies using a rotating cylinder stimulus, which consistently reported exceptionally high mean CP values (median: 0.65; interquartile range (IQR): 0.60–0.66) (Dodd et al., 2001; Grunewald et al., 2002; Krug et al., 2004, 2016). In contrast, studies using other bistable stimuli (Grunewald et al., 2002; Williams et al., 2003; Maier et al., 2007; Clark and Bradley, 2022) in MT report more moderate CP values (median: 0.55; IQR: 0.54–0.62), similar to those observed for coarse-discrimination tasks in MT (median: 0.54; IQR: 0.52–0.57). As such, the finding of elevated mean CP in bistable stimulus tasks is predominantly supported by data from rotating cylinder studies; without them, the effect disappears (model with tailoring as sensitivity control: *β*_bistable_ = 0.020, *p* = 0.3).

In conclusion, we found that mean CP was largely universal across task types once adjusted for neuronal sensitivity. The remarkably low CPs observed in fine-discrimination tasks were directly driven by correspondingly low neuronal sensitivities, reinforcing the dependence of CP on sensory tuning. The notable exception was tasks employing bistable stimuli – particularly rotating cylinders – which produced genuinely elevated CPs even after controlling for sensitivity.

This concludes our description of the four factors that robustly predict mean CP across studies. Taken together, task type, stimulus duration, brain area, and sensitivity explained about 32-34% of the variance in mean CP across studies (model with inversed N/P ratio as sensitivity control: *R_adj_* = 0.32, *N* = 88, model with tailoring as sensitivity control: *R_adj_* = 0.34, *N* = 140, see Table 4 and 5 for model fitting results). For a discussion of the studies which deviated the most from our regression model see Supplementary Information 8.5.

We also examined several additional variables listed in Fig.2 and Fig. S1 – including task exposure, lapse rate, predictability of saccade targets, and methods of CP estimation – but found no significant relationship with mean CP. For the remaining variables (psychometric kernel, reward scheme, control of eye movements, stimulus and target locations) shown in Fig.S1, too little or no data were available to permit meaningful analyses. For more details see Supplementary Information 8.3.

### 2.8 Significant factors that do not have a good theoretical explanation

We also examined several variables that lack a strong theoretical basis for influencing mean CP but still may impact it as methodological artifacts.

The specific task-relevant stimulus feature – referred to here as the task parameter – explains substantial variability in mean CP across studies. In our dataset, the most commonly used task parameters were direction of motion (29% of data points), depth (23%), self-motion (11%), and orientation (9%). Task parameter alone accounted for 24% of the variance in mean CP (*R*^2^_adj_ = 0.24; Fig.S22). As expected, task parameter is strongly coupled with the recorded brain area. To disentangle these effects, we conducted a subset analysis restricted to MT, the most represented area in our dataset. Within MT, tasks involving depth judgments were associated with significantly higher mean CP than those involving direction of motion—both in pairwise regression (*β*_depth_ = 0.075, *p <* 0.001; Fig.S23) and after controlling for sensitivity, stimulus duration, and task type (inverse mean N/P ratio as sensitivity control: *β*_depth_ = 0.053, *p* = 0.005; tailoring: *β*_depth_ = 0.048, *p* = 0.002). Notably, this pattern persisted even after excluding depth-judgment studies that used the bistable rotating cylinder stimulus – a paradigm known to produce very high CP (pairwise: *β*_depth_ = 0.057, *p <* 0.001; inverse N/P ratio: *β*_depth_ = 0.052, *p* = 0.007; tailoring: *β*_depth_ = 0.048, *p* = 0.003). The effects of recording technique, publication year, and neuron sample size—which initially appeared to correlate with mean CP in pairwise regressions (see Section 2.3)—lost statistical significance after controlling for experimental variables; furthermore, we found no effect of stimulus eccentricity (for detailed results, see Supplement 8.4).

## 3 Discussion

Our study was motivated by the surprisingly large variability in empirical choice probability (CP) measurements across the literature. Whether this variability is simply the result of measurement noise or carries a meaningful signal has remained an open question. Through our meta-analysis, we identified four systematic drivers of this variability: neuronal sensitivity, brain area, stimulus duration, and task type. Taken together, our findings support CP as a functionally meaningful, universal metric of a neuron’s involvement in perceptual decision-making, while establishing critical empirical constraints for theoretical models.

First, confirming the positive relationship between CP and neuronal sensitivity across the literature validates one of the foundational observations that motivated the field to interpret CP as a signature of a neuron’s involvement in a perceptual decision. In our meta-analysis, we observed similar positive CP-sensitivity slopes both across and within individual studies. Second, our finding that CP appears invariant across most task types – once controlled for neural sensitivity – demonstrates the metric’s universality. The notable exception –higher CPs observed in tasks utilizing bistable stimuli – reinforces the link between CP magnitude and subjective percept, as such tasks generate particularly strong changes in perceptual state. Third, our results impose new constraints on our understanding of the cortical hierarchy during decision-making. The lower CP in V1, but no significant differences among other areas, support a unique role for V1 and suggests a flexible recruitment of higher-order areas during perception and decision formation. Fourth, the positive relationship between mean CP and stimulus duration, alongside the amplified CP for bistable stimuli, provides evidence in favor of recurrent architectures, as such a relationship is difficult to reconcile with purely feedforward models. Finally, we identified unexpectedly low neuronal sensitivities as the cause for lower CPs in fine-discrimination tasks, a phenomenon that warrants future investigation. Rather than representing a task-related anomaly, this tightly coupled reduction in both CPs and sensitivities further reinforces the universal dependence of choice signals on underlying sensory tuning.

### 3.1 Neuronal sensitivity

Our results support the view that a shared underlying mechanism links CP and neuronal sensitivity both within and across studies. The observed association with tailoring technique suggests that some studies more effectively targeted task-relevant neurons, leading to higher CP estimates. The differences in neuronal sensitivity observed within individual studies have been explained in a similar way: variability in how well the task aligns with each neuron’s tuning leads to some heterogeneity in the recorded population (Britten et al., 1992). Our findings empirically support the positive CP–sensitivity relationship predicted by current models, and extend them to the across-studies level.

Notably, the magnitude of the slope of the mean CP–mean sensitivity relationship observed across and within studies poses constraints on theoretical models and suggests the possible influence of nonsensory factors. Pitkow et al. (2015) demonstrated that in the absence of decision noise, choice correlations should scale with normalized sensitivity (P/N ratio) with a slope of approximately 0.45 which is roughly 10 times larger than the across-study slope and median within-study slope available in our dataset. Within a feedforward framework, this discrepancy implies substantial suboptimality or the strong influence of non-sensory factors on perceptual choices during threshold psychophysics (Pitkow et al., 2015), compatible with previous studies pointing to serial dependencies (Urai et al., 2019), exploratory strategies (Pisupati et al., 2021), or incomplete task learning (Law and Gold, 2008) as such sources. A reanalysis of the hierarchical inference model (Haefner et al., 2016) with task-aligned feedback but incomplete learning predicts a slope of ≈ 0.15, substantially closer to the empirical value, but still about three times as large (Fig. S24). In models including feedback, lower slopes can be explained by a substantial misalignment between the optimal readout weights and the top-down feedback signals as found by some studies (Katz et al., 2016; Clery et al., 2017; Zhao et al., 2020; Levi et al., 2023).

Our finding of a significant offset in the CP–sensitivity relationship (Fig. 5), both within and across studies, challenges current theoretical models. Such an offset indicates suboptimal readout or misaligned feedback – neither of which is surprising and has been proposed previously (Clery et al., 2017; Katz et al., 2016; Zhao et al., 2020; Levi et al., 2023). However, the offset implies that the suboptimality of read-out or misaligned feedback is structured in a way that systematically depends on neuronal preferences – even neurons with near-zero sensitivity are modulated according to their preferred stimulus, resulting in CP values above 0.5. In order to explain this result it is not enough to assume limited range correlations and a discontinuity in the read-out weights at the decision boundary (Haefner et al., 2013), e.g. as in a two pool model (Shadlen et al., 1996). It also requires a discontinuity in noise correlations (Haefner et al., 2013), making them task-specific and suggestive of a feedback source (Cohen and Newsome, 2008; Haefner et al., 2016; Bondy et al., 2018; Lange and Haefner, 2022; Liu et al., 2026). Alternatively, saccade planning signals could account for this offset by inverting the preferences of weakly tuned neurons (Laamerad et al., 2024, Fig. S7). The positive intercept in the relationship across studies, may also point to a publication bias, resulting from the systematic underreporting of experiments that yield both low neuronal sensitivities and mean CP values below 0.5.

### 3.2 Brain area

We showed that despite V1 and MT having comparable task sensitivities, MT exhibits significantly higher mean CP – on average 0.03–0.04 higher after controlling for other variables (Fig. 6c). Several other areas also tended to show higher mean CP than V1, though most comparisons did not reach statistical significance, likely due to limited sample sizes. The limited choice-related activity in V1 mirrors its characteristically weak top-down attentional modulation compared to higher sensory areas (McAdams and Maunsell, 1999; Buffalo et al., 2010; Maunsell, 2015). These observations suggest that V1 holds a special status within the visual hierarchy – serving as a general-purpose input layer whose output is distributed broadly across downstream areas but not directly involved in perception or decision mechanism (Kravitz et al., 2013). Although V1 receives rich top-down input from higher-order regions (Crick and Koch, 1995; Lee and Mumford, 2003; Liu et al., 2024), it may lack the direct task-aligned signals from frontoparietal areas seen in downstream regions such as V4 and MT (Gilbert and Li, 2013). Our results challenge the view of V1 as a ‘cognitive blackboard’ (Roelfsema and de Lange, 2016) or an active saliency map within a recurrent feedback loop (Zhaoping, 2019), as both hypotheses predict robust signatures of choice-related activity in the primary visual cortex.

The studies reporting relatively high mean CP in V1 argue that the decision circuitry can access this area directly, but only under specific conditions. Palmer et al. (2007) attributed their exceptionally high mean CP in V1 (0.62) to their neurons’ high sensitivity (mean N/P ratio of 1.2), yet other studies with similar sensitivities report much lower V1 CPs (Kang and Maunsell, 2020; Nienborg and Cumming, 2014) – a pattern confirmed in our dataset and not explained by differences in sensitivity, stimulus duration, or task type (Fig. 5c). Nienborg and Cumming (2014) proposed that the decision area selectively accesses neurons organized into task-relevant cortical columns: V1’s orientation columns but not disparity columns predict CP for orientation but not disparity discrimination (Nienborg and Cumming, 2006). Evidence remains mixed: among V1 studies using orientation tasks, two reported significant CP (Nienborg and Cumming, 2014; Bondy et al., 2018) while two others did not (Goris et al., 2017; Lange et al., 2023), despite comparable neuronal sensitivities.

The absence of a clear hierarchical trend beyond V1 could reflect methodological biases and limited statistical power rather than a functional flexibility in decision-related signaling. Because the functional properties of areas like V1 and MT are well understood, tasks targeting them are typically highly optimized. In contrast, studies in less-characterized regions may employ sub-optimal tasks, confounding fair cross-area CP comparisons. Furthermore, our dataset is severely unbalanced: MT accounts for nearly half of all data points, leaving other areas sparsely represented. This limits our statistical power to definitively resolve CP ordering across the visual hierarchy or to generalize the effects of stimulus duration and task type across all brain regions.

### 3.3 Stimulus duration

Our finding of a positive relationship between mean CP and stimulus duration provides evidence for a feedback contribution to the CP phenomenon beyond the evidence provided by the previously observed increase of instantaneous CP within a trial (Uka and DeAngelis, 2004; Nienborg and Cumming, 2006, 2009). This finding provides constraints on existing models regarding the scaling of feedback signals – whether beliefs or post-decision feedback – with trial length, suggesting feedback becomes stronger or longer *as a fraction of* stimulus duration. A promising next step would be to validate our findings within a single study by employing multiple stimulus durations in order to distinguish between different types of feedback signals. Belief-related signals which are more likely present during evidence accumulation, are predicted to be aligned with stimulus tuning and hence increase CPs (Haefner et al., 2016; Lange and Haefner, 2022). While task-aligned, late-trial post-decision signals could theoretically explain the positive CP-duration relationship, this contradicts recent evidence that such signals are predominantly orthogonal or even anti-aligned with the task (Zhao et al., 2020; Levi et al., 2023; Zahorodnii et al., 2025).

### 3.4 Task type

The tasks that elicit the largest CPs appear to be those involving bistable stimuli, a conclusion based on experiments using the rotating cylinder stimulus (Dodd et al., 2001; Krug et al., 2004, 2016) and binocular flash suppression (Maier et al., 2007). Despite more than two decades of research, a comprehensive framework explaining the neural mechanisms underlying bistable perception remains elusive. Early explanations focused on winner-take-all feedforward models, in which stochastic fluctuations in sensory neurons within two competing pools flip the percept (Taylor and Aldridge, 1974; Wilson, 2003). While influential, such models struggle to explain why only some ambiguous stimuli evoke bistable perception. Hierarchical inference models link CP directly to decision confidence (Haefner et al., 2016; Lange and Haefner, 2022), and because bistable stimuli evoke highly confident percepts (Bainbridge et al., 2015), those models naturally predict elevated CPs – a hypothesis testable using recent paradigms that simultaneously measure choice and confidence in monkeys (So and Stuphorn, 2016; Cai et al., 2022; Boundy-Singer et al., 2025). Alternatively, Wasmuht et al. (2019) suggested that bistable stimuli evoke stronger feedback to stabilize the dominant percept rather than to communicate a stronger prior. Puzzlingly, Williams et al. (2003) tested a stimulus conceptually related to the rotating cylinder (displaced columns of dots) but found no significant CP at all. Therefore, replicating the large CPs evoked by the rotating cylinder using different bistable stimuli – ideally targeting regions beyond area MT – is important for excluding stimulus-specific confounds.

After controlling for other experimental variables, we found no significant differences in mean CP between coarse- and fine-discrimination or detection tasks, suggesting they rely on the same underlying decision-making process – consistent with theoretical models that make no distinction between task types (Haefner et al., 2013; Chicharro et al., 2021). Our findings did confirm that neuronal sensitivity is lower in fine- than in coarse-discrimination tasks, a pattern observed across multiple areas including V2 (Clery et al., 2017), MT (Purushothaman and Bradley, 2005; Uka and DeAngelis, 2006; Price and Born, 2010; Yu and Gu, 2018), and MST (Yu and Gu, 2018), supporting the view that this is a universal rather than area-specific phenomenon. Critically, once neuronal sensitivity was matched, mean CP was indistinguishable between task types – arguing against the hypothesis that fine-discrimination tasks engage a different decision-making process by drawing on larger neuronal pools (Xu et al., 2014; Kim et al., 2015). This raises the question of what drives the sensitivity difference itself. In a feedforward framework, it has been suggested that task-relevant information may simply be more diffusely represented in fine-discrimination tasks (Purushothaman and Bradley, 2005; Uka and DeAngelis, 2006). Alternatively, in hierarchical inference models both lower sensitivity and lower CP may be due to weaker feedback-driven redistribution of task information by feedback signals (Liu et al., 2026) – a signal that may be insufficiently precise to be effective in fine-discrimination settings.

### 3.5 Other conclusions

Only three data points in our dataset show a mean CP significantly below 0.5. Such “negative” CPs contradict feedforward models, where increased neuronal activity should support the preferred choice. Interestingly, the negative CPs in V1 and V2 reported by Goris et al. (2017) and in MT by Levi et al. (2023) both emerged late in the trial. Furthermore, Levi et al. (2023) observed this negative CP only in the task condition that promoted early decision commitment. This timing suggests negative CP may reflect post-decision feedback that is oppositely aligned to neuronal preferences—a mechanism hypothesized to protect decision variables maintained in the working memory from ongoing sensory interference (Zahorodnii et al., 2025). Nevertheless, the low incidence of such findings leaves open the possibility that they merely reflect low-probability sampling noise or idiosyncratic behavioral strategies. It is highly plausible that such misalignment is present in many other datasets, but simply lacks the magnitude required to overcome the primary drivers of “positive” mean CP.

We found a clear decline effect in reported mean CP over time (Fig. 3a), similar to trends observed in other fields (Schooler, 2011; Pietschnig et al., 2019). However, unlike the classic explanation of publication bias, our analysis reveals that this decline is primarily driven by methodological shifts. Over three decades, studies gradually moved away from precisely tailoring stimuli to individual neurons leading to a drop in neural sensitivity (Fig. 3c) – a compromise necessary for collecting larger neuronal samples (Fig. S27). A secondary contributor to the decline effect is the expansion of CP studies beyond area MT into brain regions with inherently lower CP, most notably V1 (Fig. S28). Publication bias may also have played a role: during the early years of the field, null results from V1 may have been less likely to be published.

Our findings suggest that the method used to calculate CP does not systematically bias mean CP estimates across studies. Concerns surrounding CP computation have long been a major focus of the field, with numerous studies examining potential artifacts (Britten et al., 1996; Kang and Maunsell, 2012; Nienborg and Cumming, 2009; Zaidel et al., 2017; Chicharro et al., 2021). Our results show that such methodological variation is unlikely to be a major concern. However, given the relatively small sample size, we caution against over-interpreting these null results—particularly the finding that preference estimation method (passive-viewing versus task trials) had no measurable effect on mean CP. This may hold true for most studies in our dataset, where recorded neurons likely had strong feedforward tuning and were therefore less susceptible to preference flips driven by feedback (Zaidel et al., 2017). However, in chronic multi-electrode recordings and new Neuropixel technology, where low-sensitivity neurons are more common, this issue may still introduce substantial bias.

Small eye movements during fixation—such as oculomotor drift and microsaccades—can also confound CP measurements by simultaneously modulating neural responses and correlating with behavioral choices (Britten et al., 1996; Dodd et al., 2001; Herrington et al., 2009; Nienborg and Cumming, 2014; Verhoef and Maunsell, 2017). This may also impact the observed relationships between mean CP and brain area (as eye movement effects may be more pronounced for smaller receptive fields) or stimulus duration (as the probability of microsaccades increases over time).

Recent work suggests that CP may also be influenced by saccade planning signals (Laamerad et al., 2024; Zhang and Gu, 2026). However, our data were insufficient to rigorously test the hypothesis (see Supplement 8.3).

Studies also inevitably differ in the laminar composition of their recorded neurons due to specific recording techniques and experimenter choices, which may systematically affect CP (Steinfeld et al., 2024); unfortunately, explicit data on laminar depth is largely unavailable. Likewise, differing reward schemes can impact the responses of sensory neurons (Takagaki and Krug, 2020) and potentially influence the strength of choice signals, but a standardized methodology for across-study comparison is currently lacking.

### 3.6 Link between CP and population metrics

Recent advances in simultaneous population recordings have prompted the development of multivariate metrics to generalize the concept of CP (Zhao et al., 2020; Levi et al., 2023; Bondy et al., 2025). Unlike traditional CP, which evaluates neurons in isolation, these modern approaches typically employ a choice decoder—a model trained to predict the animal’s behavioral choice from the entire population’s activity. This choice decoder is primarily characterized by two quantities: its accuracy and its weight profile. Notably, some authors have even referred to this decoding accuracy directly as “choice probability” (Zhao et al., 2020).

However, a fundamental distinction exists between choice decoder accuracy and traditional CP. While a choice decoder captures all available choice-related information in a population regardless of sensory tuning, standard CP strictly quantifies choice signals that align with a neuron’s stimulus preference. Therefore, the most accurate population-level equivalent to CP is the accuracy of an optimal stimulus decoder in predicting behavioral choices (for trials with ambiguous or constant stimulus). Conceptually, an individual neuron’s CP represents its fractional contribution to this stimulus decoder’s accuracy but only in the population of independent neurons. Because inter-neuronal correlations introduce redundancy, the overall population accuracy is substantially lower than the sum of the individual contributions.

Equation (1) bridges the single-neuron CP metric and population-level concepts. Specifically, the readout weights (**w**) in Equation (1) represent the choice decoder weights, while the neuronal preferences (sign(*f_i_*)) correspond to the signs of the stimulus decoder weights. Under theoretical optimal readout, the weights of the choice and stimulus decoders should be perfectly aligned. Yet, empirical studies consistently reveal a substantial misalignment between them (Katz et al., 2016; Clery et al., 2017; Zhao et al., 2020; Levi et al., 2023). Consequently, traditional mean CP captures only a fraction of the total choice-related activity in a population (i.e. choice decoder accuracy) – not merely because it is a single-neuron metric, but fundamentally because it is blind to any choice signals not aligned with the population’s sensory tuning. Whether this is a feature or a bug depends on the scientific question.

### 3.7 Suggestions for Future Studies

Our meta-analysis demonstrates the continued utility of the CP metric, and we strongly encourage future studies to keep reporting it. The field’s shift toward large-scale population recordings raises the question of how best to aggregate individual CPs into a population-level metric. Since current population metrics (Bondy et al., 2018; Zhao et al., 2020; Levi et al., 2023) – particularly the accuracy of decoders – are difficult to compare across studies, reporting CP remains highly beneficial. However, we propose several methodological refinements:

#### Report CP-normalized sensitivity slopes alongside mean CP

Comparing mean CP across studies yields limited insights without accounting for the neuronal sensitivity of the sampled populations (section 2.4). The CP-sensitivity slope provides a directly comparable and theoretically motivated metric that incorporates both neuronal preferences and task sensitivity, thereby better constraining decision-making models. Researchers should transition from the common practice of reporting correlation coefficients between CP and neurometric thresholds, and instead report the slope of CP versus normalized sensitivity—for example, the psychometric-to-neurometric threshold ratio (P/N ratio) or the equivalent ratio of neuronal to behavioral sensitivity (*d^′^_neuron_*/*d^′^_behavior_*). Theoretically, the CP-threshold relationship is highly nonlinear and correlation coefficients confirm a monotonic trend but fail to quantify its magnitude. Furthermore, the intercept in a CP-threshold regression lacks a meaningful interpretation, while the CP-sensitivity offset does. Conversely, the CP-normalized sensitivity slope maps directly onto theoretical predictions and enables cross-study comparisons (Pitkow et al., 2015). Because P/N ratios can vary significantly depending on the choice of the underlying metrics (Prince et al., 2000), we propose additionally reporting the standard neuron-anti-neuron metric (Section 7).

#### Estimate sensitivity independently of choice signals

Interpreting CP–sensitivity relationships is complicated by the standard practice of estimating neuronal sensitivity using trials from the task itself—data that may be “contaminated” by choice-related top-down signals (Zaidel et al., 2017). Sensitivity measured this way inevitably reflects a mixture of feedforward and feedback components, whereas most theoretical models define sensitivity strictly in feedforward terms. Partial correlation methods can be used to separate stimulus-driven and choice-related components of neural activity (Zaidel et al., 2017), or sensitivity can be computed conditionally on the choice. However, a significant limitation of these computational approaches is their inability to isolate feedback that varies in strength within a single choice condition. For instance, internal variables like decision confidence may vary even when the choice remains the same, leading to feedback-driven modulation of apparent sensitivity that conditional metrics cannot remove. Consequently, while limited by reduced statistical power, sensitivity estimates derived from passive-viewing sessions may offer the cleanest separation from choice signals and the closest alignment with theoretical frameworks, albeit with the caveat that over-trained animals may still covertly perform the task.

#### Report asymptotic extrapolations of population metrics

The accuracy of both choice decoders and optimal stimulus decoders in predicting behavior depends on the number of neurons in the recorded sample. One way to facilitate comparisons between studies is by reporting the dependence of accuracy on number of neurons within the population, and in particular their asymptotic limit for infinitely large populations.

#### New approaches to tailoring stimuli

Recent nonparametric advances in closed-loop active stimulus optimization (Yamane et al., 2008; Wang and Ponce, 2022, 2026; Willeke et al., 2025) promise to significantly advance stimulus tailoring. This approach should yield higher neural sensitivities and CPs. Furthermore, by mitigating the poor responsiveness of higher level areas to traditional hand-crafted stimuli, these methods will facilitate CP comparisons across different brain areas.

#### Non-sensory manipulations of the percept

Historically, neural stimulation and inactivation have been the primary tools used to establish a causal relationship between specific neuronal populations and perception (Liu and Newsome, 2005; Uka and DeAngelis, 2006; Shiozaki et al., 2012; Verhoef et al., 2012; Yu and Gu, 2018). Advances in targeted neural stimulation provide novel methods to probe individual neuronal contributions to perception (Shahbazi et al., 2024). Linking these contributions with CP measurements from the same neurons represents a potentially fruitful direction for future studies.

#### Standardize reporting and maximize data sharing

While our meta-analysis benefited from the field’s convention of reporting mean CP and basic experimental parameters, missing data remained a significant hurdle – especially when standard metrics (like N/P ratios) were omitted or when novel metrics (like the CP–P/N ratio slope) were required. Beyond basic task sensitivity, detailed behavioral metrics (e.g., lapse rates, psychometric kernels), spatial configurations (stimulus size, stimulus and target locations), and physiological characteristics (e.g., baseline firing rates, receptive fields, cortical layer, cell type, and latency) should be reported. Although main texts typically focus on population summaries, we urge researchers to share comprehensive per-neuron and per-session data in supplementary materials or public repositories to maximize data reuse and power future large-scale meta-analyses (Fig. 1b).

### 3.8 Conclusion

Meta-analytic approaches offer a powerful complement to individual studies by pooling information across tasks, animals, and laboratories, thereby increasing the robustness of conclusions and isolating the underlying drivers of neural measurements – an approach that will remain valuable even as the number of simultaneously recorded neurons continues to grow. This meta-study underscores the value of field-wide standardization: it was only possible because a large number of primate neurophysiology labs consistently reported identical, well-established quantitative measures. For choice probability specifically, the present study has (i) provided evidence for a remarkably consistent and general relationship with neural sensitivity, (ii) identified both expected and unexpected empirical drivers of its magnitude, (iii) proposed extensions to population-level measures better suited to modern large-scale recordings, and (iv) offered concrete recommendations for future studies to enhance its utility in constraining models of sensory processing.

## 4 Acknowledgments

We thank Christopher R. Fetsch for valuable feedback on an early draft of the manuscript, and Yong Gu for assistance with data reanalysis. This work was supported by the National Institutes of Health (NIH) grants EY028811 to R.M.H., EY029438 and EY035005 to A.R., EY032999 to R.L.T.G., EY105260 to J.I.G.; the Retina Research Foundation Edwin and Dorothy Gamewell Professorship at the McPherson Eye Research Institute to A.R.; a Canadian Institutes of Health Research (CIHR) grant PJT178071 to C.C.P.; an Israel Science Foundation (ISF) grant (No. 1291/20) to A.Z.; National Science Foundation (NSF) CAREER awards to R.M.H. (IIS-2143440) and R.L.T.G. (#2146369); the German Research Foundation (425899996, SFB 1436) to K.K.

## 5 Contributions

Conceptualization: R.M.H., A.P., G.C.D., C.C.P., R.T.B.; Data analysis: A.P.; Writing –original draft: A.P., R.M.H.; Data curation: A.P.; Data preprocessing: A.P., G.C.D., C.C.P., R.L.T.G., C.M.Z., A.Z., A.J.L., A.R., T.U., R.D., P.L., R.D.L., X.Y., I.F.; Writing – review & editing: R.M.H., A.P., G.C.D., C.C.P., R.L.T.G., C.M.Z., A.Z., A.J.L., A.R., H.N., M.S., A.C.H., A.J.P., K.K., R.T.B., I.F., J.I.G.

## 6 Data Availability

The complete curated dataset and associated experimental variables compiled for this meta-analysis are publicly available on Google Sheets at: https://docs.google.com/spreadsheets/ d/1NZ6SM1AhTI6UqxGe0b6JOxsWTUqzsDsHZIXqa9xZ5nc/edit?usp=sharing.

## 7 Method

### 7.1 Data Collection

We included studies in our dataset based on the following criteria: (1) the study reported data from macaque monkeys of any species (most commonly Macaca mulatta, but also M. fuscata); (2) the data came from sensory areas of the visual cortex; and (3) the study reported a population mean choice probability (CP). We excluded studies involving sensorimotor regions – in particular, VIP and LIP – due to the prevalence of motor-related signals in those areas. However, we retained data from the caudal intraparietal area (CIP), as prior work has shown that motor-related signals in CIP do not exceed those observed in V3A (Doudlah et al., 2022; Zhu et al., 2024). Excluding CIP from our dataset did not qualitatively affect the results.

Our literature search followed a three-stage process. First, we compiled an initial list of CP studies known to us. We then identified additional studies by recursively examining the references cited in these papers. Second, we used AI-based tools – including ChatGPT, NotebookLM, and Undermind AI(OpenAI, 2023; Research, 2023; AI, 2024) – to identify further relevant publications. Third, we contacted authors of known CP papers and asked whether they were aware of additional studies not yet included.

From each study, we extracted a predefined set of experimental variables (see dataset). This list was developed based on: (1) prior empirical and theoretical findings; (2) open hypotheses proposed in the CP literature; and (3) our own theoretical considerations grounded in existing CP models. The list was further refined with input from CP study authors, who were invited to suggest additional variables. All variables were manually extracted by the first author. Some variables added later in the process – notably the CP–sensitivity correlation coefficient – were extracted using Gemini (DeepMind, 2023). When multiple papers were based on the same or overlapping neural datasets, we retained only one, preferring the study with the larger or more complete dataset. After each paper was processed, we contacted the corresponding authors to verify the extracted data and to fill in any missing information.

We aimed to collect data on a per-monkey basis; however, 22 out of the 59 analyzed papers included data that were either partially or fully pooled across multiple monkeys. Many CP studies also reported results from multiple brain areas and/or tasks. As a result, the number of data points contributed by each study varied, depending on how many monkeys, brain areas, and task conditions it included. While the median number of data points per study was two – as expected for a typical CP study with two monkeys – the interquartile range (IQR) was 1.5 to 4, with a full range from 1 to 7. In some studies, although mean CP was available on a per-monkey basis, certain experimental variables – specifically the mean N/P ratio, significance of the mean CP, within-study CP–sensitivity slope, and alternative CP metrics – were reported only for data combined across monkeys. In these cases, for analyses involving such variables, the per-monkey data points were replaced with a single data point representing the combined data.

### 7.2 CP metric

In our meta-analysis, we aimed to use a *CP metric* that was as consistent as possible across studies. Whenever available, we used the grand CP computed from random-seed stimuli, as this was the most commonly reported method and was available for 62.5% of data points. If this metric was unavailable, we selected the closest available alternative based on the following hierarchy: (1) grand CP with frozen seeds (5%), (2) CP calculated from ambiguous stimulus trials with random seeds (20.5%), and (3) CP from ambiguous stimulus trials with frozen seeds (12%).

When a study reported *mean CP for different subsets of neurons*, we selected the estimate most comparable across studies. A common practice in CP studies is to exclude neurons that are not sufficiently tuned to the stimulus – either during neuron selection (in non-chronic recordings) or during data analysis. This choice is justified, as the mean estimate (in our case, mean CP) is more meaningful when calculated over a relatively homogeneous population with few outliers (Huber and Ronchetti, 2009). However, not all studies applied such filtering and instead reported population means that included low-sensitive neurons; some of them additionally reported mean CP values for a subset of task-sensitive neurons. When such filtered estimates were available, we included them in the dataset in place of the unfiltered population mean, in order to improve comparability across studies.

In the study by Purushothaman and Bradley (2005), neurons were filtered not based on task sensitivity, but on whether their preferred direction included an upward component – a criterion not strictly tied to sensitivity. This decision was motivated by the authors’ observation that only neurons with an upward component exhibited CP values significantly above 0.5. While this methodological choice may be justifiable given that the reference direction in their fine-discrimination task was always upward, it introduces a bias that complicates comparison with other studies, which did not filter neurons based on CP. To improve comparability, we estimated the full population mean CP by calculating a simple, unweighted average of the two reported subpopulation means (upward- and downward-preferring). We intentionally avoided a weighted average, as the original study’s screening process artificially underrepresented the downward-preferring neurons.

Several studies included in our dataset did not originally publish mean CP values. For these studies, the original authors calculated and provided these metrics specifically for this meta-analysis. Below, we briefly outline the methods used to compute these newly provided mean CPs and, where applicable, their corresponding Neurometric-to-Psychometric (N/P) ratios.

#### Chang et al. (2020) and Doudlah et al. (2022)

Choice probabilities were measured in V3A and CIP while monkeys performed an eight-alternative, forced-choice tilt discrimination task (Chang et al., 2020; Doudlah et al., 2022). During the task, three-dimensional planar surface orientation tuning was measured at four stimulus distances. On each trial, the planar tilt (which side of the plane was closest to the monkey) was reported via a saccadic eye movement to one of eight choice targets corresponding to the eight presented tilt angles. Choice probabilities were computed using the task ambiguous, frontoparallel plane (slant = 0°, tilt undefined) trials only. Responses at each distance were z-scored and then pooled. Behavioral responses to the two choice targets corresponding to the preferred and anti-preferred tilts determined from the surface orientation tuning curves were then selected for the CP calculation. At least three trials in each of the two choice directions were required for inclusion. In CIP, the average CPs were 0.56 (95% CI: 0.55, 0.58; N = 197) and 0.65 (95% CI: 0.62, 0.67; N = 212) for monkeys L and F, respectively. In V3A, the average CPs were 0.52 (95% CI: 0.51, 0.54; N = 260), 0.60 (95% CI: 0.58, 0.63; N = 243), and 0.56 (95% CI: 0.54, 0.58; N = 113), for monkeys L, F, and W, respectively.

#### Lange et al. (2023)

Because the original study reported CPs without aligning them to neuronal preferences, we realigned these values using the sign of the reported *d^′^*. Mean CPs were then calculated by pooling data across tasks (solo and interleaved epochs) for each monkey. For Monkey A, data from two separate recording epochs (2017 and 2019) were also combined. Finally, 95% confidence intervals were derived using hierarchical bootstrapping across electrodes and days. The N/P ratio was calculated as *d^′^_monkey_*/*d^*′*^_*neuron*_*, utilizing the originally reported values for *d^′^_neuron_*. Behavioral sensitivity (*d^′^_monkey_*) was derived from the psychometric curve slope (es_√_timated via psignifit) by assuming a standard cumulative Gaussian: 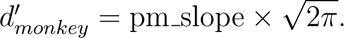

### 7.3 Other variables

Most of the experimental variables used in our analysis are described in sufficient detail in the main text. In this section, we provide additional information for a subset of variables where further clarification or methodological detail may be useful.

Because the standard error of the mean CP (SEM) was explicitly reported for only 29 data points, we estimated SEM for additional cases using other information provided in the papers. For 9 data points, SEM was derived from reported confidence intervals, and for 13 data points it was estimated from reported p-values under the assumption of a Gaussian distribution of CP values within a study.

We systematically classified *tailoring* along three dimensions: (1) stimulus size, (2) task-relevant parameter (e.g., orientation in an orientation discrimination task or direction of motion in direction judgments), and (3) task-irrelevant parameters that influence responsiveness but are not related to the discrimination (e.g., spatial frequency in an orientation discrimination task or speed in direction judgments). Each parameter was classified into one of four tailoring levels: (1) not fit (a stimulus parameter was predetermined and fixed), (2) fit to the recorded population (typically in multi-electrode recordings), (3) fit to individual neurons (only available in single-electrode recordings) and (4) additional level only for stimulus size in case the stimulus is smaller than the receptive field of the recorded neuron. This tailoring classification was applied independently to each of the three stimulus dimensions.

To standardize reported mean N/P ratios across our dataset, we addressed two primary methodological discrepancies. First, because CP studies typically employ one-interval behavioral tasks while neurometric functions often rely on two opposing d_√_is<u>tr</u>ibutions (Hillis et al., 2004; Elmore et al., 2019), we multiplied the neuronal thresholds by 2 for any studies where this adjustment had not already been made. Second, studies differ in whether they compare responses to two opposite stimuli (the neuron-antineuron method) (Britten et al., 1992) or to one task stimulus versus an ambiguous stimulus (Prince et al., 2000; Goris et al., 2017; Elmore et al., 2019; Ziemba et al., 2024). Because these methods yield significantly different N/P ratios without a simple analytical conversion (Prince et al., 2000), we asked authors of the latter group to re-estimate their values using the more prevalent neuron-antineuron method. Original values were retained only when re-estimation was unavailable (Elmore et al., 2019). Apart from the adjustments described above, we could not account for all minor methodological variations in neural and behavioral threshold/sensitivity computation; however, we do not expect these remaining differences to introduce systematic confounds beyond standard estimation noise.

To summarize the *reported* relationships between Choice Probability (CP) and neuronal sensitivity, we extracted reported Spearman or Pearson correlation coefficients from 29 studies, yielding 51 distinct data points (Fig. S6). The compiled studies quantified this relationship using two primary metrics: 28 data points (from 18 papers) evaluated CP against neurometric thresholds, while the remaining data points utilized measures directly reflecting sensitivity. Given that the neurometric threshold is the mathematical inverse of sensitivity, we standardized these statistics prior to synthesis by reversing the sign of the correlation coefficients for all threshold-based studies. While this sign-reversal serves as a practical approximation rather than a strictly formal mathematical identity, it facilitated a unified analysis of the aggregated dataset.

We requested that authors of the included studies provide within-study slopes and intercepts for the relationship between CP and normalized sensitivity (P/N ratio), or share raw neuron-level data for us to estimate these parameters. To minimize the impact of behavioral noise on sensitivity estimates, the median psychometric threshold across sessions was used; consequently, a single, constant psychometric threshold was applied to all neurons for a given monkey-and-task combination. To ensure methodological consistency across all datasets neurometric thresholds were derived using the neuron-antineuron method and scaled by 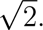 Finally, for two specific datasets (Nienborg and Cumming, 2006; Clery et al., 2017), rather than performing de novo estimations from neuron-level data, we utilized the relationships originally reported as choice correlation versus P/N ratio (Clery et al., 2017).

For the majority of data points in our meta-analysis (85%), the *stimulus duration* matched the length of the spike count window used for CP calculations. For most of the remaining data points (13%), the spike count window was shorter than the stimulus duration, as the authors selected a time window—typically during the middle or late phase of the stimulus presentation—that maximized the CP estimate. In these cases, we still used the full stimulus duration, rather than the spike count window, as the value for stimulus duration. However, for two studies involving extremely brief stimulus pulses in detection tasks, the spike count window substantially exceeded the actual stimulus duration (Herrington et al., 2009; Smith et al., 2011). In these cases, we used the spike count window instead.

We estimated task exposure as the sum of the pre-recording training period and half of the recording duration, reflecting the monkey’s cumulative experience with the task.

### 7.4 Statistical analysis

All statistical analyses were performed in R (R Core Team, 2025), using base functions and contributed packages as appropriate.

We performed linear regression analyses under the assumption that the dependent variable, mean CP, is approximately normally distributed. While CP values are theoretically bounded between 0 and 1, the observed means in our dataset ranged from 0.46 to 0.69 and did not approach the boundaries, supporting the use of Gaussian residuals. To ensure robustness, we additionally conducted beta regression using the betareg package with a logit link function, which is specifically designed for modeling continuous proportions (Jackman, 2001; Zeileis et al., 2023). Both approaches yielded qualitatively similar results, identifying the same set of predictors with statistically significant coefficients.

We did not include random effects in our regression models to account for the hierarchical structure of the data. This was primarily due to sample size limitations: our dataset contained repeated measurements from the same studies and monkeys, but modeling random intercepts for all 59 studies (or all individual monkeys) would have required substantially more data than was available. Although the data are not fully independent, we note that the number of observations contributed by each study was relatively modest and fairly evenly distributed, reducing the risk that a small number of studies dominated the results (see Section 7.1). Modeling random effects for monkeys posed an additional challenge, as individual animals could appear in multiple studies, and we did not systematically track such cross-study identities. We tested the relationship between mean CP and various experimental variables using a nested regression approach, beginning with measures of neuronal sensitivity. Two alternative sensitivity metrics were used: (1) the inverse of the mean N/P ratio, which was available for 59% of data points; and (2) a set of three variables describing the tailoring technique, which serves as a cruder proxy for neuronal sensitivity but is available for the full dataset. These two sensitivity measures were used in two separate regression models. Notably, the theoretically motivated experimental variables (listed in Fig. 2) that showed a significant effect were consistent across both models. Only additive effects of experimental dimensions were investigated, as the small sample size did not allow for the inclusion of multiplicative interactions. The two final regression equations are shown below (see Table 4 and 5 for model fitting results):

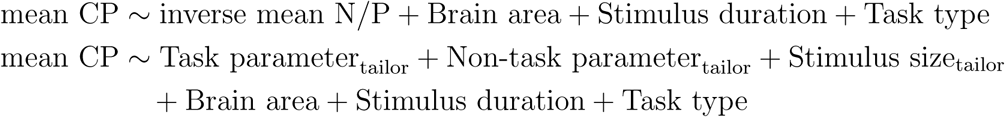

Note that most of the experimental variables tested were categorical rather than continuous, and were therefore modeled in the regression as sets of independent binary (dummy) variables. We considered an experimental variable to have a significant effect on mean CP if any of its categories showed a statistically significant (*p <* 0.05) regression coefficient. All variables included in the equations above met this criterion, with the exception of Non-task parameter_tailor_ and Stimulus size_tailor_, which did not contain any categories with significant effects. These two variables were retained in the model for consistency, as we consider them components of a unified tailoring construct that better explains variation in the inverse N/P ratio than Task parameter_tailor_ alone. Excluding these two variables had no appreciable impact on the regression results.

Additionally, to assess whether each experimental variable contributed significantly to the explanatory power of the regression models, we conducted Type II ANOVA tests using the Anova() function from the car package (Fox and Weisberg, 2023). This approach evaluates the unique effect of each variable after accounting for all others in the model. In the regression model that used the tailoring-based sensitivity proxy, the ANOVA results were consistent with our main significance criterion: all variables except Non-task parameter_tailor_ and Stimulus size_tailor_ showed a statistically significant contribution to mean CP (Task parameter_tailor_: *p* = 0.02; Brain area: *p* = 0.007; Stimulus duration: *p* = 0.0016; Task type: *p* = 0.0001). In contrast, in the regression model using the inverse mean N/P ratio as the sensitivity measure, the inverse mean N/P ratio itself (*p* = 0.02), Stimulus duration (*p* = 0.001) and Task type (*p* = 0.03) reached significance, whereas Brain area (*p* = 0.12) did not. However, collapsing the detailed brain area variable into a binary factor (V1 versus non-V1) yielded a significant main effect (p = 0.03) without altering the significance of the remaining predictors. This pattern indicates that the initial lack of significance was an artifact of low statistical power—driven by the small sample sizes in areas other than V1 and MT—and corroborates our primary conclusion that V1 plays a distinct functional role. Taken together, and despite some limitations, the ANOVA results support the conclusion that neuronal sensitivity, brain area, stimulus duration, and task type each contributed significantly to explaining variability in mean CP across studies.

## 8 Supplement

### 8.1 Tables

**Table 1:**
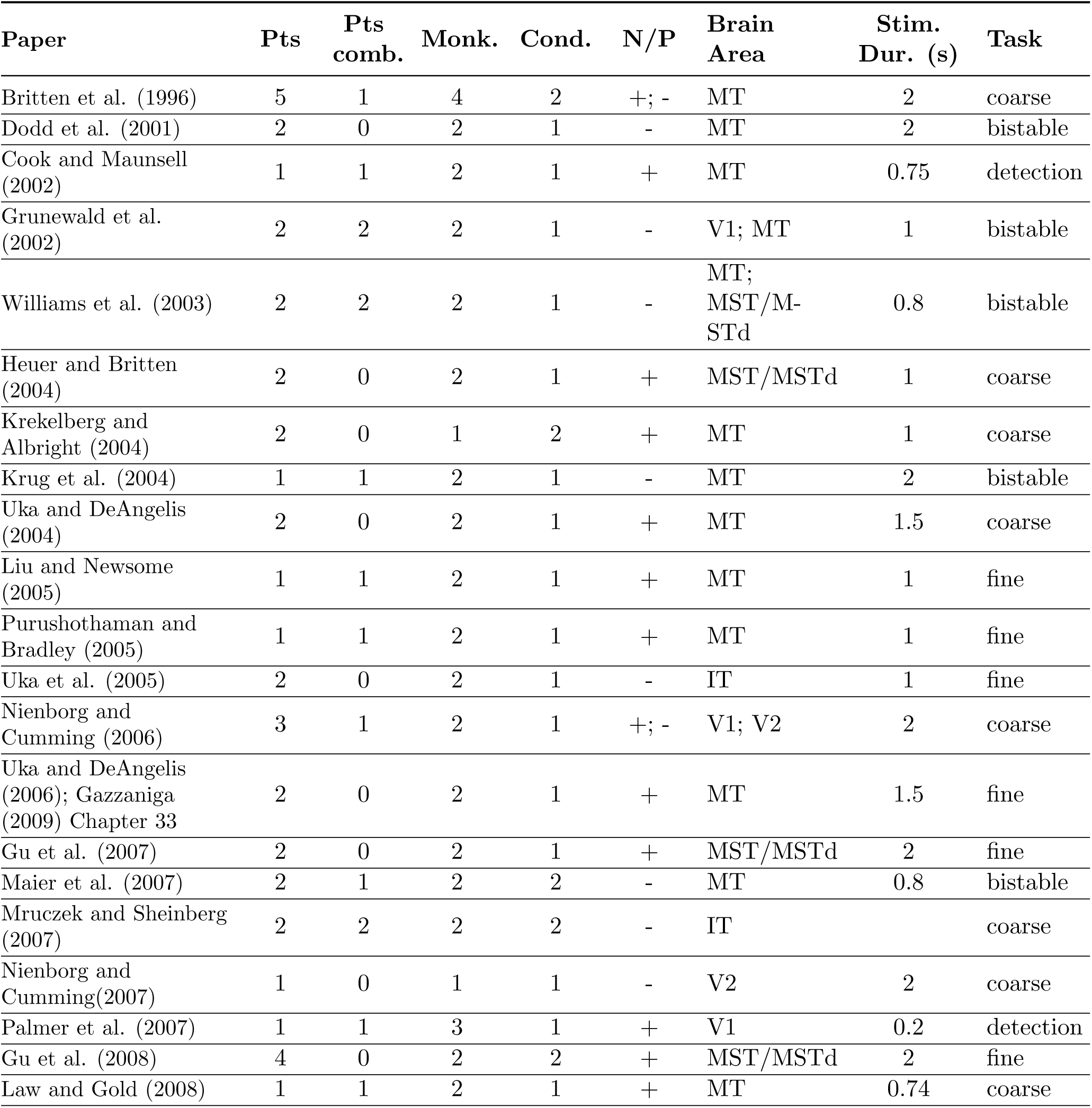
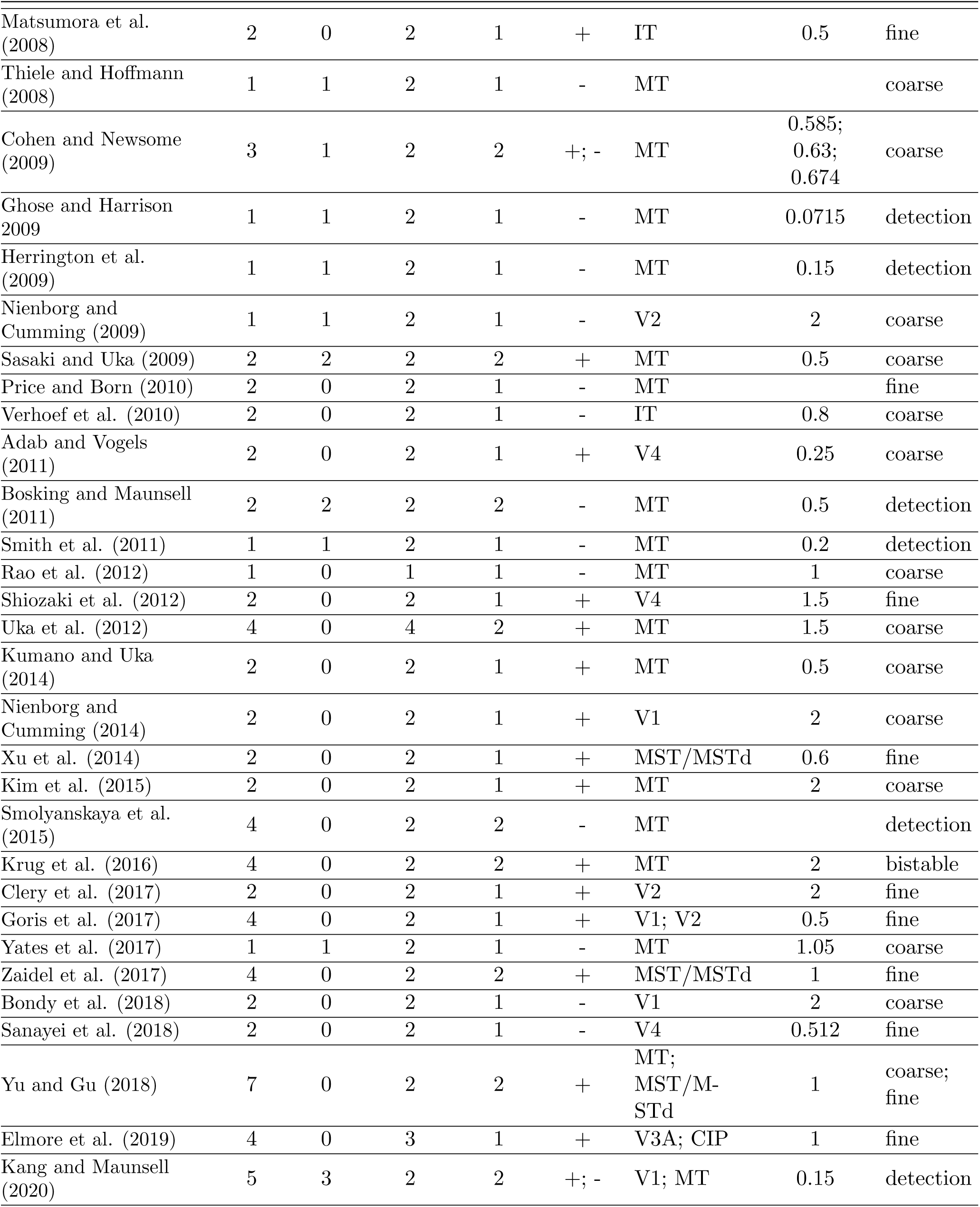
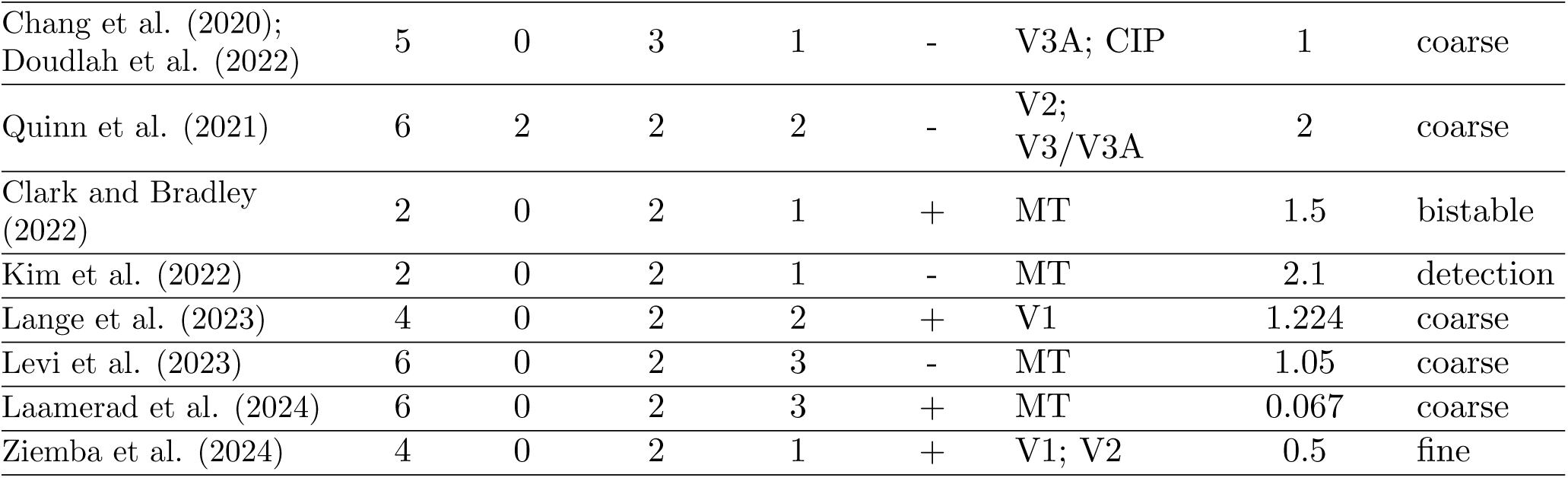
Summary of papers used in the meta-analysis (see also full dataset). **Pts**: number of data points; **Pts comb.**: number of data points for which the mean CP was estimated from pooled data across monkeys; **Monk.**: number of monkeys; **Cond.**: number of task conditions; **N/P**: whether N/P ratio was reported (if mixed, shown as “+; -”); **Stim. Dur. (s)**: stimulus duration; **Task**: task type.

**Table 2:** Explanation of terms used in Fig. 2 and Fig. S1.

| Terms | Description |
| --- | --- |
| Decoder type | Characteristics of the assumed readout mechanism used to estimate the stimulus category from neural responses – including its linearity, optimality, and the composition of the readout pool (Shadlen et al., 1996; Haefner et al., 2013; Pitkow et al., 2015) |
| Read out | Weight profile of the decoder – the contribution of each neuron in the readout pool to the decoder’s output |
| Covariability | The trial-to-trial covariability in the responses of sensory neurons when the stimulus is fixed; commonly referred to as noise correlations (Kohn et al., 2016) |
| Sensitivity | A measure of how accurately a neuron discriminates between two stimulus categories in a task (Britten et al., 1992) |
| Brain area | Functional and anatomical brain region from which neural activity was recorded and CP was calculated (Kravitz et al., 2011) |
| Decision confidence | Subjective certainty about the choice (Pouget et al., 2016) |
| Engagement <sup>3</sup> | Level of subject’s involvement and motivation during the task Aston-Jones and Cohen (2005) |
| Task learning <sup>3</sup> | How well the subject has learned the task-relevant stimulus-choice mapping (Gold and Shadlen, 2007) |
| Perceptual learning <sup>3</sup> | Improvement in the ability to discriminate or detect sensory stimuli through practice or experience (Gilbert et al., 2001; Sagi, 2011) |

| **Terms** | **Description** |
| --- | --- |
| Tailoring stimulus | The degree to which different stimulus properties were adjusted to match the tuning characteristics of recorded neurons (Britten et al., 1992) |
| Task type | Classification of perceptual decision tasks: 1. Coarse-discrimination 2. Fine-discrimination 3. Detection 4. Stimulus with bistable perception |
| Stimulus duration | Duration for which the stimulus is presented during a trial |
| Lapse rate | Proportion of errors on very perceptually easy trials (Prins, 2012; McCarley and Yamani, 2021) |
| Task exposure | Amount of time the subject spent on the task before and during data collection |
| Neuronal preferences | How neuron’s preferences used in CP calculation were derived: 1. From task trials 2. From separate passive viewing sessions |
| Psych. (psychometric) kernel <sup>4</sup> | Weights that quantify the contribution of different stimulus features to the subject’s choice (Nienborg and Cumming, 2009) |
| Reward scheme | The specific structure of reinforcement provided during the task, including the magnitude, probability, and symmetry of payoffs across different choices (Rorie et al., 2010) |
| Confounds | Factors not directly related to stimulus processing or decision-making that can nonetheless alter CP estimates |
| Saccade planning | Motor preparation signals related to eye movements to a choice target (Sommer and Wurtz, 2008) |
| Frozen vs random seed | CP is calculated using only trials with truly identical stimuli vs. stimuli with varying noise (Britten et al., 1996) |
| Zero signal vs grand CP | CP is calculated using only trials with truly ambiguous stimuli vs. combining z-scored responses from several signal strengths (Britten et al., 1996) |
| Control of eye movements | Methods that control for small movements of the eyes during fixation that cause variability in the retinal image (Gur et al., 1997; Bair and O’Keefe, 1998) |
| Stimulus and targets location | Position of the stimulus and choice targets relative to fixation point |
| Predictability of saccade targets | Quantified as the consistency or variability in target locations across trial and days (Basso and Wurtz, 1997) |

**Table 3:**
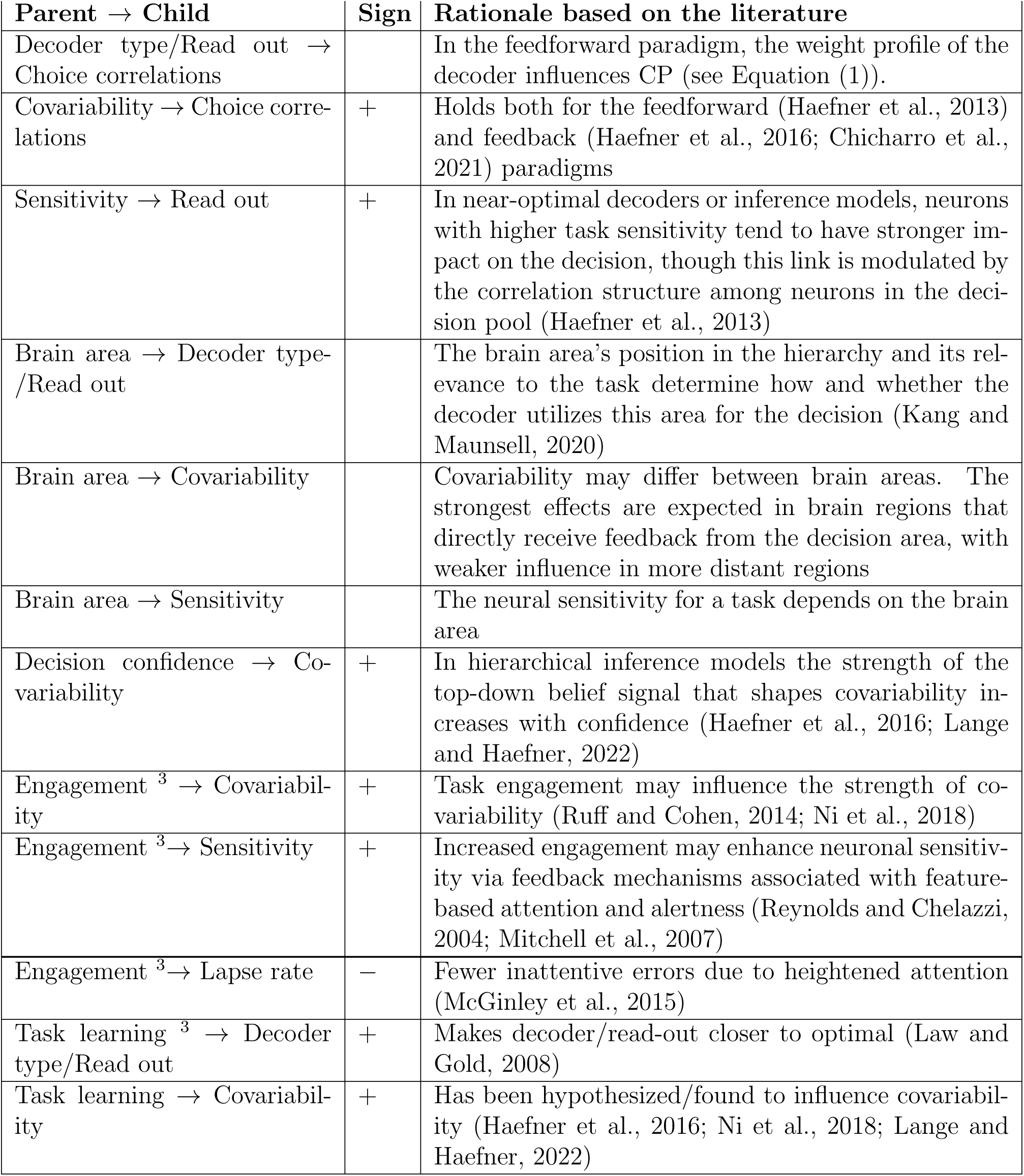
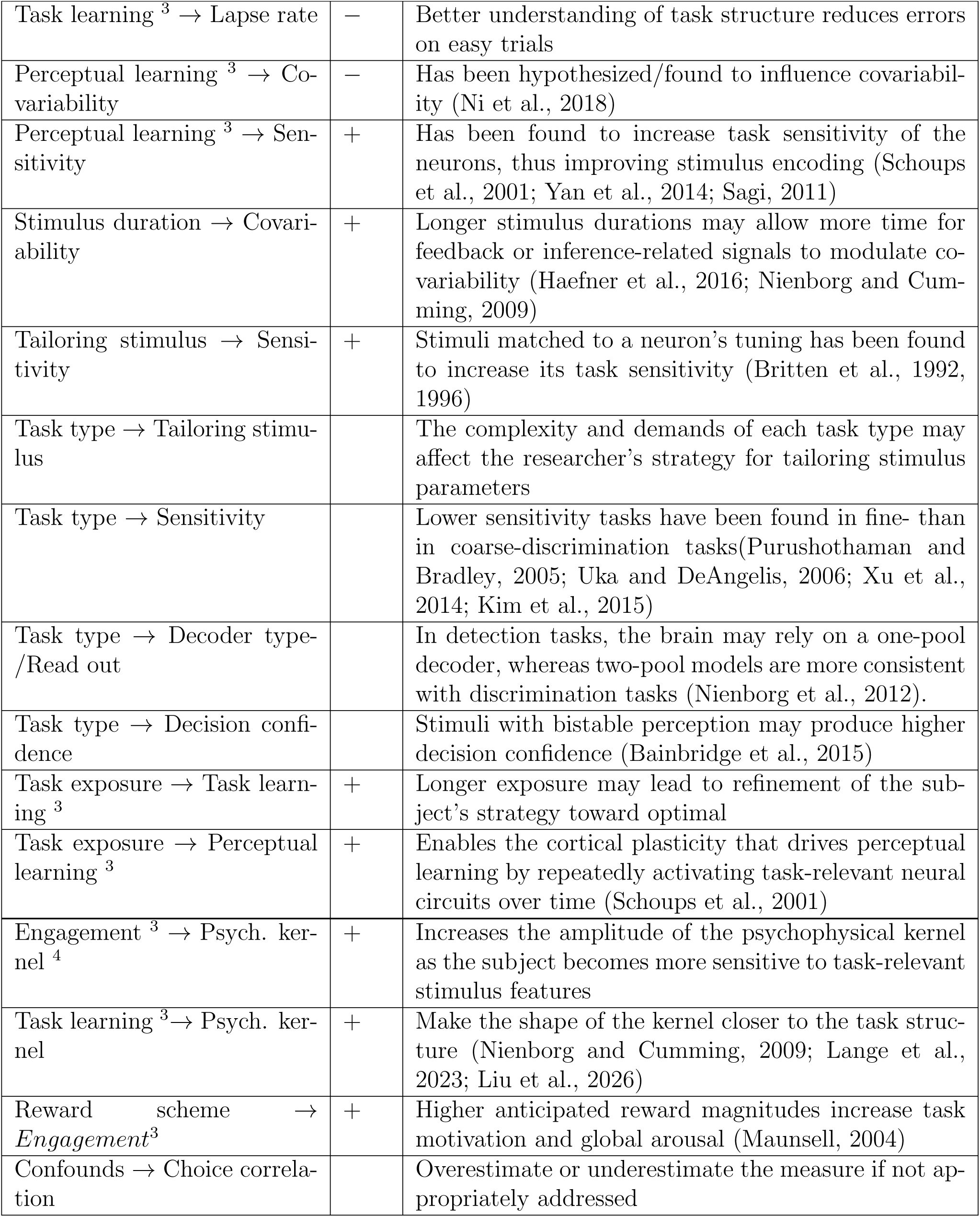
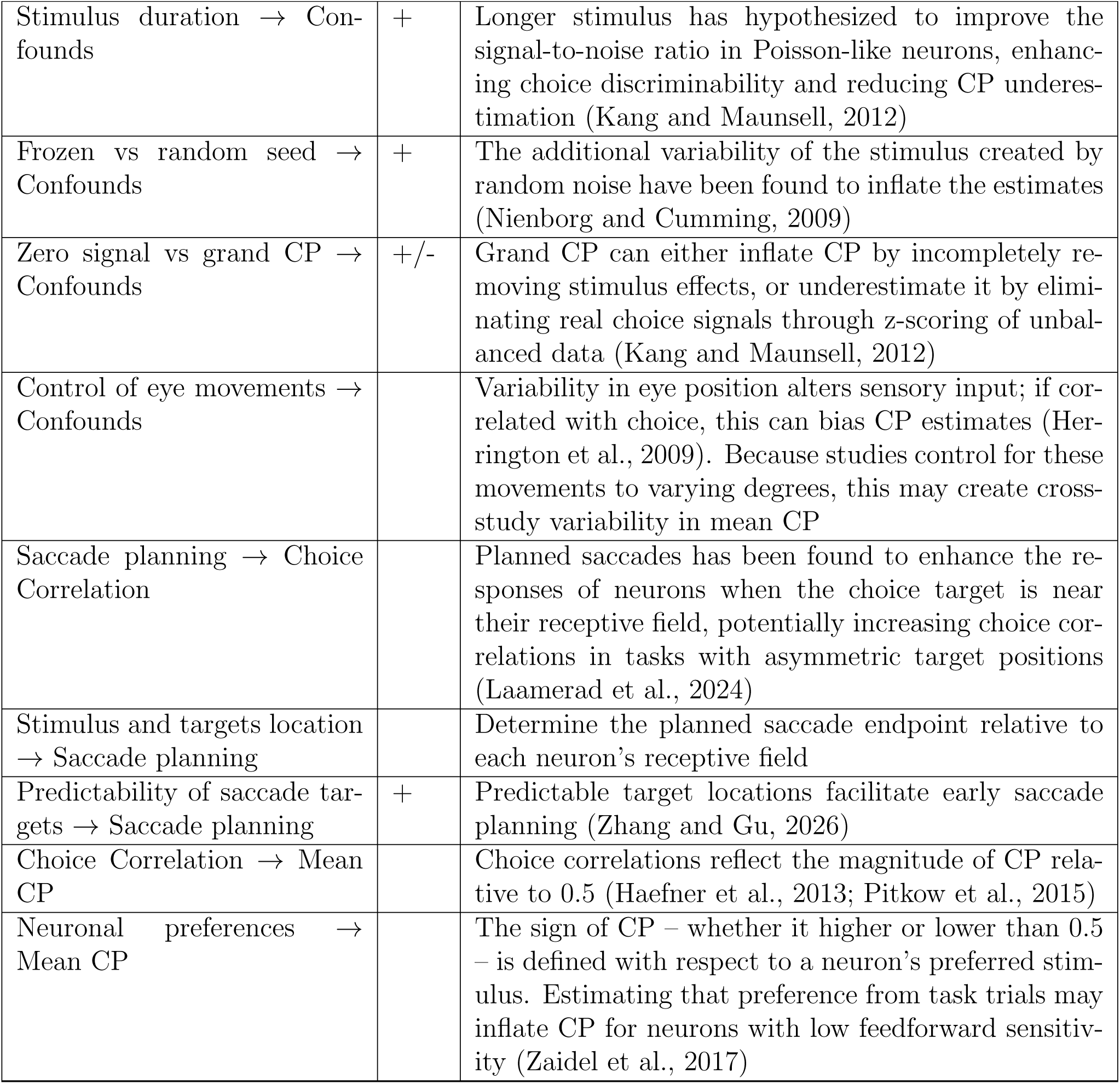
Annotation of the arrows in Fig. 2 and Fig. S1. The “Sign” column indicates the expected direction of influence. A missing sign means that the direction cannot be specified, either because the relationship is non-monotonic or because one of the variables lacks a hierarchical ordering.

| Parent → Child | Sign | Rationale based on the literature |
| --- | --- | --- |
| Decoder type/Read out → Choice correlations |  | In the feedforward paradigm, the weight profile of the decoder influences CP (see Equation (1)). |
| Covariability → Choice correlations | + | Holds both for the feedforward (Haefner et al., 2013) and feedback (Haefner et al., 2016; Chicharro et al., 2021) paradigms |
| Sensitivity → Read out | + | In near-optimal decoders or inference models, neurons with higher task sensitivity tend to have stronger impact on the decision, though this link is modulated by the correlation structure among neurons in the decision pool (Haefner et al., 2013) |
| Brain area → Decoder type-/Read out |  | The brain area’s position in the hierarchy and its relevance to the task determine how and whether the decoder utilizes this area for the decision (Kang and Maunsell, 2020) |
| Brain area → Covariability |  | Covariability may differ between brain areas. The strongest effects are expected in brain regions that directly receive feedback from the decision area, with weaker influence in more distant regions |
| Brain area → Sensitivity |  | The neural sensitivity for a task depends on the brain area |
| Decision confidence → Covariability | + | In hierarchical inference models the strength of the top-down belief signal that shapes covariability increases with confidence (Haefner et al., 2016; Lange and Haefner, 2022) |
| Engagement <sup>3</sup> → Covariability | + | Task engagement may influence the strength of covariability (Ruff and Cohen, 2014; Ni et al., 2018) |
| Engagement <sup>3</sup> → Sensitivity | + | Increased engagement may enhance neuronal sensitivity via feedback mechanisms associated with feature-based attention and alertness (Reynolds and Chelazzi, 2004; Mitchell et al., 2007) |
| Engagement <sup>3</sup> → Lapse rate | − | Fewer inattentive errors due to heightened attention (McGinley et al., 2015) |
| Task learning <sup>3</sup> → Decoder type/Read out | + | Makes decoder/read-out closer to optimal (Law and Gold, 2008) |
| Task learning → Covariability | + | Has been hypothesized/found to influence covariability (Haefner et al., 2016; Ni et al., 2018; Lange and Haefner, 2022) |

| Parent → Child | Sign | Rationale based on the literature |
| --- | --- | --- |
| Task learning <sup>3</sup> → Lapse rate | – | Better understanding of task structure reduces errors on easy trials |
| Perceptual learning <sup>3</sup> → Co-variability | – | Has been hypothesized/found to influence covariability (Ni et al., 2018) |
| Perceptual learning <sup>3</sup> → Sensitivity | + | Has been found to increase task sensitivity of the neurons, thus improving stimulus encoding (Schoups et al., 2001; Yan et al., 2014; Sagi, 2011) |
| Stimulus duration → Covariability | + | Longer stimulus durations may allow more time for feedback or inference-related signals to modulate covariability (Haefner et al., 2016; Nienborg and Cumming, 2009) |
| Tailoring stimulus → Sensitivity | + | Stimuli matched to a neuron’s tuning has been found to increase its task sensitivity (Britten et al., 1992, 1996) |
| Task type → Tailoring stimulus | | The complexity and demands of each task type may affect the researcher’s strategy for tailoring stimulus parameters |
| Task type → Sensitivity | | Lower sensitivity tasks have been found in fine- than in coarse-discrimination tasks(Purushothaman and Bradley, 2005; Uka and DeAngelis, 2006; Xu et al., 2014; Kim et al., 2015) |
| Task type → Decoder type-/Read out | | In detection tasks, the brain may rely on a one-pool decoder, whereas two-pool models are more consistent with discrimination tasks (Nienborg et al., 2012). |
| Task type → Decision confidence | | Stimuli with bistable perception may produce higher decision confidence (Bainbridge et al., 2015) |
| Task exposure → Task learning <sup>3</sup> | + | Longer exposure may lead to refinement of the subject’s strategy toward optimal |
| Task exposure → Perceptual learning <sup>3</sup> | + | Enables the cortical plasticity that drives perceptual learning by repeatedly activating task-relevant neural circuits over time (Schoups et al., 2001) |
| Engagement <sup>3</sup> → Psych. kernel <sup>4</sup> | + | Increases the amplitude of the psychophysical kernel as the subject becomes more sensitive to task-relevant stimulus features |
| Task learning <sup>3</sup> → Psych. kernel | + | Make the shape of the kernel closer to the task structure (Nienborg and Cumming, 2009; Lange et al., 2023; Liu et al., 2026) |
| Reward scheme → Engagement<sup>3</sup> | + | Higher anticipated reward magnitudes increase task motivation and global arousal (Maunsell, 2004) |
| Confounds → Choice correlation | | Overestimate or underestimate the measure if not appropriately addressed |
| Stimulus duration → Confounds | + | Longer stimulus has hypothesized to improve the signal-to-noise ratio in Poisson-like neurons, enhancing choice discriminability and reducing CP underestimation (Kang and Maunsell, 2012) |
| Frozen vs random seed → Confounds | + | The additional variability of the stimulus created by random noise have been found to inflate the estimates (Nienborg and Cumming, 2009) |
| Zero signal vs grand CP → Confounds | +/- | Grand CP can either inflate CP by incompletely removing stimulus effects, or underestimate it by eliminating real choice signals through z-scoring of unbalanced data (Kang and Maunsell, 2012) |
| Control of eye movements → Confounds |  | Variability in eye position alters sensory input; if correlated with choice, this can bias CP estimates (Herrington et al., 2009). Because studies control for these movements to varying degrees, this may create cross-study variability in mean CP |
| Saccade planning → Choice Correlation |  | Planned saccades has been found to enhance the responses of neurons when the choice target is near their receptive field, potentially increasing choice correlations in tasks with asymmetric target positions (Laamerad et al., 2024) |
| Stimulus and targets location → Saccade planning |  | Determine the planned saccade endpoint relative to each neuron’s receptive field |
| Predictability of saccade targets → Saccade planning | + | Predictable target locations facilitate early saccade planning (Zhang and Gu, 2026) |
| Choice Correlation → Mean CP |  | Choice correlations reflect the magnitude of CP relative to 0.5 (Haefner et al., 2013; Pitkow et al., 2015) |
| Neuronal preferences → Mean CP |  | The sign of CP – whether it higher or lower than 0.5 – is defined with respect to a neuron’s preferred stimulus. Estimating that preference from task trials may inflate CP for neurons with low feedforward sensitivity (Zaidel et al., 2017) |

**Table 4:**
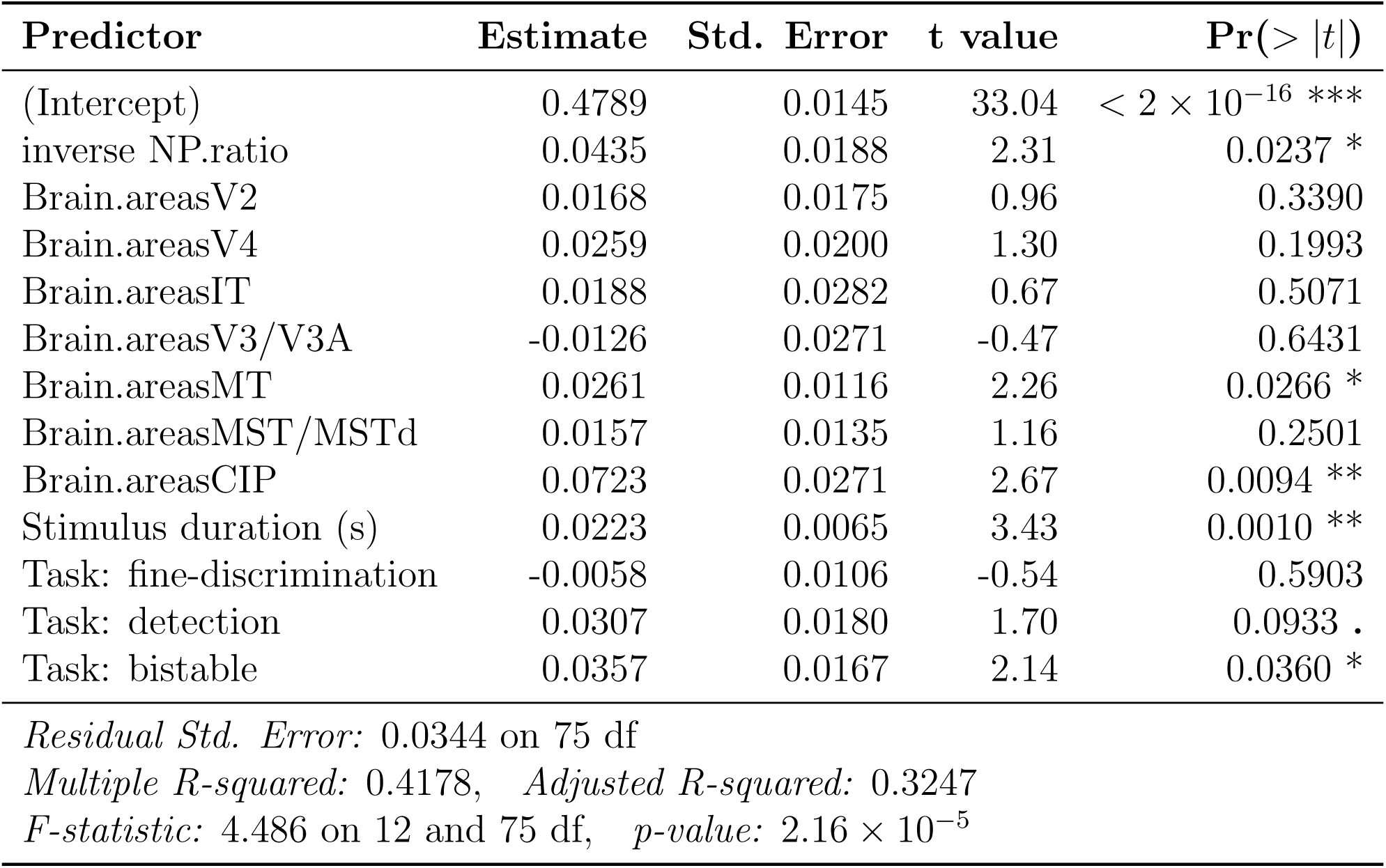
Linear regression results predicting mean CP using inverse mean N/P ratio, brain area, stimulus duration, and task type.

| Predictor | Estimate | Std. Error | t value | Pr(> \|t\|) |
| --- | --- | --- | --- | --- |
| (Intercept) | 0.4789 | 0.0145 | 33.04 | $< 2 \times 10^{-16}$ *** |
| inverse NP.ratio | 0.0435 | 0.0188 | 2.31 | 0.0237 * |
| Brain.areasV2 | 0.0168 | 0.0175 | 0.96 | 0.3390 |
| Brain.areasV4 | 0.0259 | 0.0200 | 1.30 | 0.1993 |
| Brain.areasIT | 0.0188 | 0.0282 | 0.67 | 0.5071 |
| Brain.areasV3/V3A | -0.0126 | 0.0271 | -0.47 | 0.6431 |
| Brain.areasMT | 0.0261 | 0.0116 | 2.26 | 0.0266 * |
| Brain.areasMST/MSTd | 0.0157 | 0.0135 | 1.16 | 0.2501 |
| Brain.areasCIP | 0.0723 | 0.0271 | 2.67 | 0.0094 ** |
| Stimulus duration (s) | 0.0223 | 0.0065 | 3.43 | 0.0010 ** |
| Task: fine-discrimination | -0.0058 | 0.0106 | -0.54 | 0.5903 |
| Task: detection | 0.0307 | 0.0180 | 1.70 | 0.0933 . |
| Task: bistable | 0.0357 | 0.0167 | 2.14 | 0.0360 * |
| *Residual Std. Error:* 0.0344 on 75 df |  |  |  |  |
| *Multiple R-squared:* 0.4178, *Adjusted R-squared:* 0.3247 |  |  |  |  |
| *F-statistic:* 4.486 on 12 and 75 df, *p-value:* $2.16 \times 10^{-5}$ | | | | |

**Table 5:** Linear regression results predicting mean CP using tailoring-based sensitivity proxy, brain area, stimulus duration, and task type.

| Predictor | Estimate | Std. Error | t value | Pr(> \|t\|) |
| --- | --- | --- | --- | --- |
| (Intercept) | 0.4912 | 0.0192 | 25.59 | $< 2 \times 10^{-16}$ *** |
| Task_parm: population | -0.0324 | 0.0153 | -2.12 | 0.0360 * |
| Task_parm: single neuron | 0.0056 | 0.0105 | 0.53 | 0.5947 |
| Non_task_parm: population | 0.0036 | 0.0135 | 0.27 | 0.7871 |
| Non_task_parm: single neuron | 0.0105 | 0.0107 | 0.98 | 0.3284 |
| Stimulus size: fit to population | 0.0058 | 0.0194 | 0.30 | 0.7659 |
| Stimulus size: fit to RF | -0.0071 | 0.0188 | -0.38 | 0.7042 |
| Stimulus size: smaller than RF | 0.0028 | 0.0216 | 0.13 | 0.8988 |
| Brain areasV2 | 0.0388 | 0.0157 | 2.47 | 0.0149 * |
| Brain areasV4 | 0.0468 | 0.0203 | 2.31 | 0.0226 * |
| Brain areasIT | 0.0374 | 0.0247 | 1.51 | 0.1332 |
| Brain areasV3/V3A | 0.0292 | 0.0199 | 1.47 | 0.1438 |
| Brain areasMT | 0.0369 | 0.0129 | 2.87 | 0.0049 ** |
| Brain areasMST/MSTd | 0.0244 | 0.0183 | 1.34 | 0.1838 |
| Brain areasCIP | 0.0869 | 0.0255 | 3.41 | 0.0009 *** |
| Stimulus duration (s) | 0.0194 | 0.0060 | 3.22 | 0.0016 ** |
| Task: fine-discrimination | -0.0212 | 0.0092 | -2.31 | 0.0226 * |
| Task: detection | 0.0235 | 0.0129 | 1.82 | 0.0711 . |
| Task: bistable | 0.0434 | 0.0133 | 3.26 | 0.0015 ** |
| *Residual Std. Error:* 0.0374 on 121 df |  |  |  |  |
| *Multiple R-squared:* 0.4276, *Adjusted R-squared:* 0.3424 |  |  |  |  |
| *F-statistic:* 5.021 on 18 and 121 df, *p-value:* $2.33 \times 10^{-8}$ | | | | |

### 8.2 Figures

**Figure S1:**
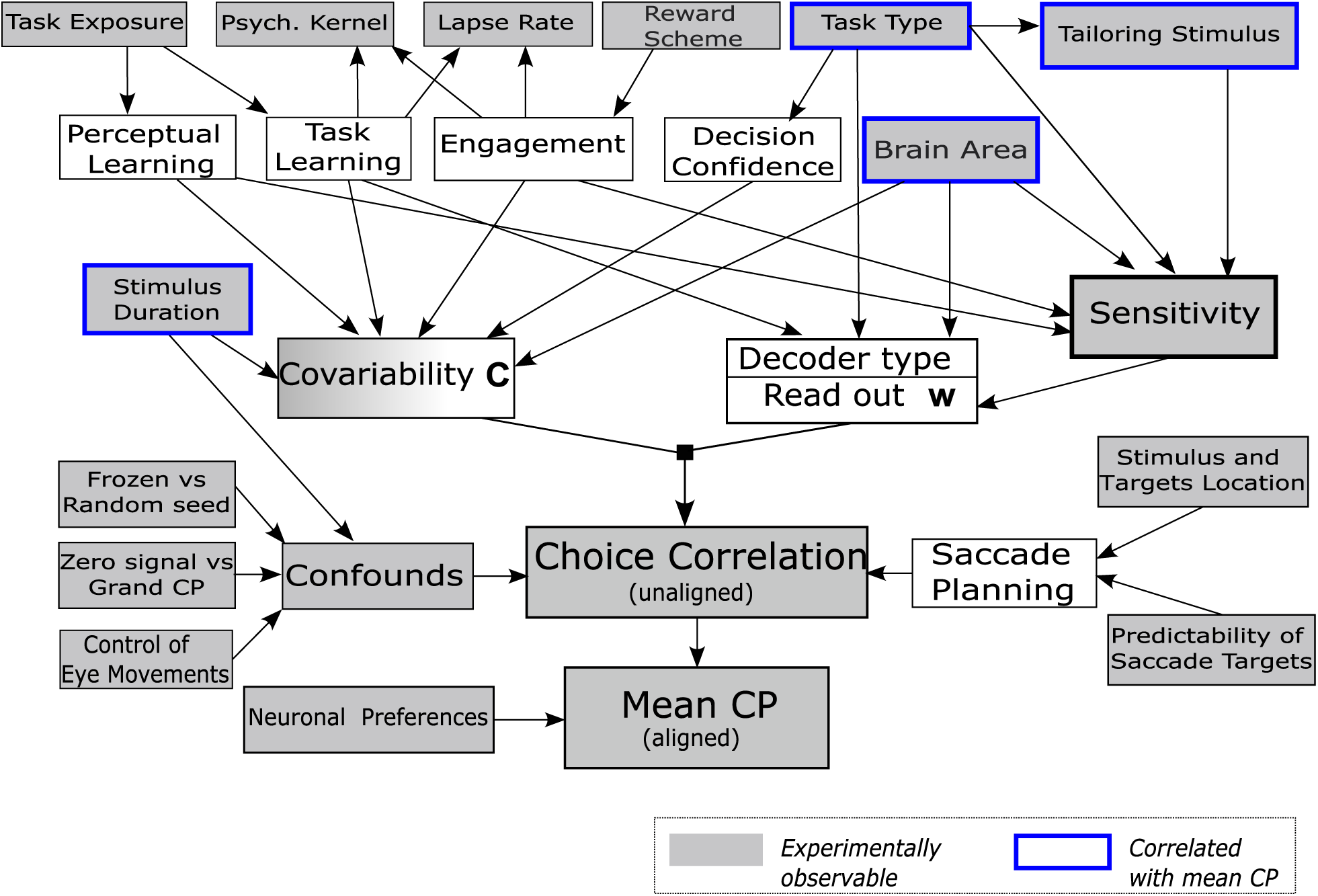
Expanded schematic of hypothesized factors influencing Choice Probability (CP). This supplementary diagram extends the framework presented in Fig. 2 by incorporating additional elements: (1) potential confounds and pre-motor signals, and (2) experimental factors that could not be evaluated in this meta-analysis due to data limitations (Psych. Kernel, Reward Scheme). It was proposed that at least part of the observed choice correlations may be explained by pre-motor signals during saccade planning (Laamerad et al., 2024; Zhang and Gu, 2026)(lower right part of the diagram). Methodological choices that date back to Britten et al. (1996) such as CP estimation procedures (“Grand CP”) and stimulus control techniques (“frozen noise”, eye movement control) may influence CP estimates (Nienborg and Cumming, 2009; Kang and Maunsell, 2012; Herrington et al., 2009) (lower left of the diagram). Additionally, compared to Fig. 2 the “Learning and Engagement” node has been subdivided into “Task learning,” “Perceptual learning,” and “Engagement.” Arrows denote putative causal relationships between experimental design choices, intermediate variables, and core factors driving CP (see Tables 2 and 3 for details). Gray boxes denote experimentally observable variables. Thick blue borders highlight variables that exhibited a statistically significant relationship with mean CP in our analysis.

**Figure S2:**
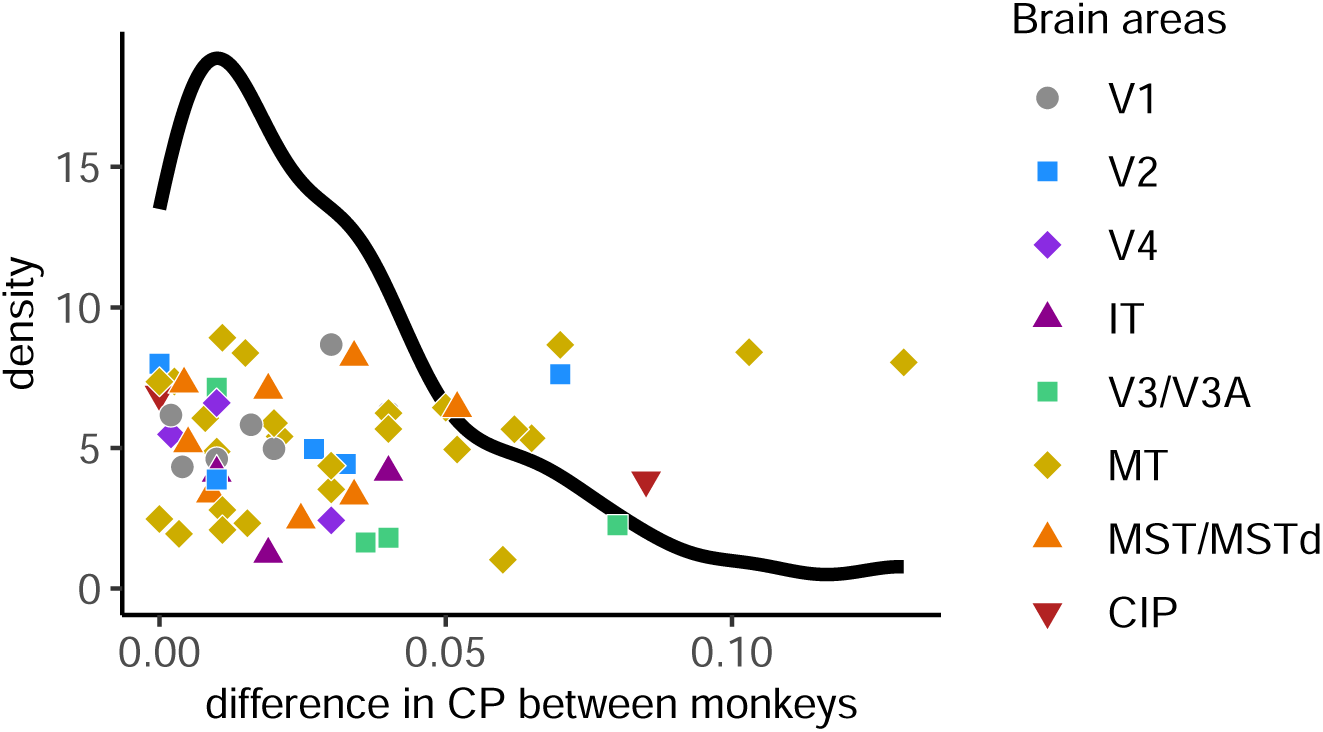
Differences in mean CP between monkeys within the same study. Each point shows the difference between two monkeys, computed by subtracting the lower CP value from the higher. Vertical jitter was added for visualization. The black curve is a density function

**Figure S3:**
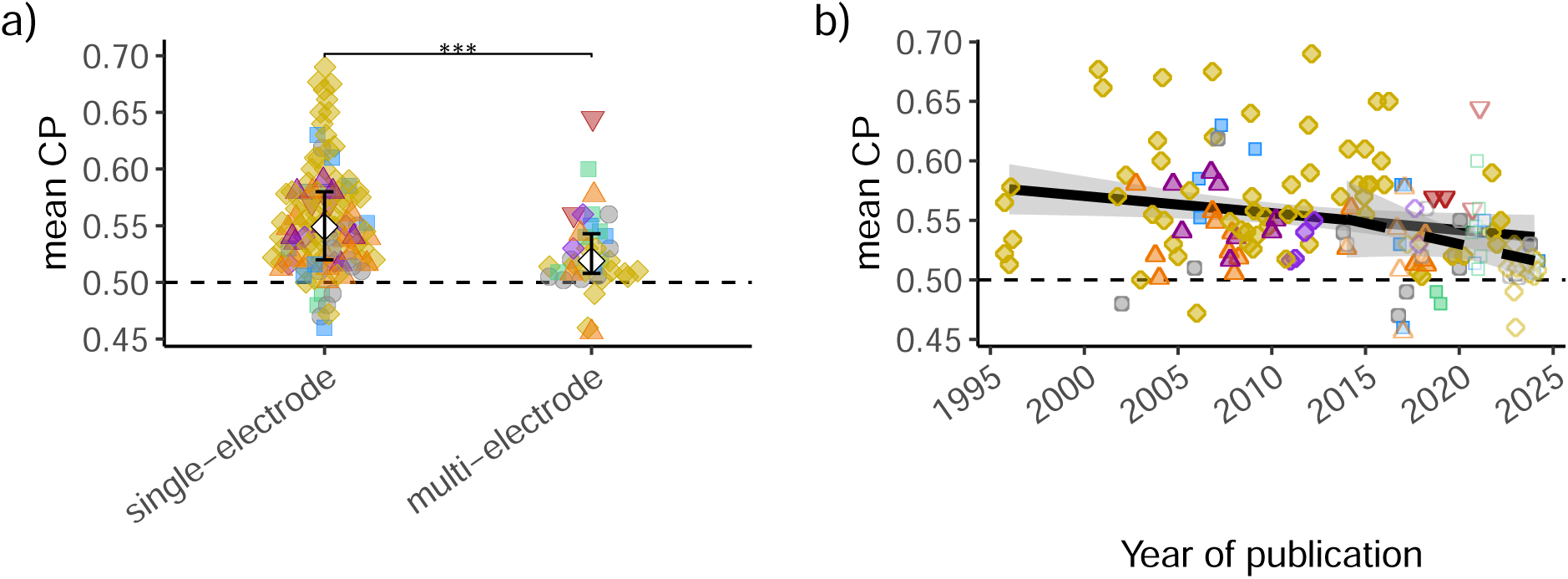
Mean CP is lower in multi-electrode studies and declines over time in single-electrode studies. **(a)** Mean CP by recording technique. Multi-electrode studies report significantly lower CP values (*** *p <* 0.001). **(b)** Relationship between mean CP and year of publication, shown separately for single-electrode (filled symbols, solid regression line) and multi-electrode studies (open symbols, dashed line). A significant decline is observed even when restricting the analysis to single-electrode studies (*β* = −0.0015, *p* = 0.02).

**Figure S4:**
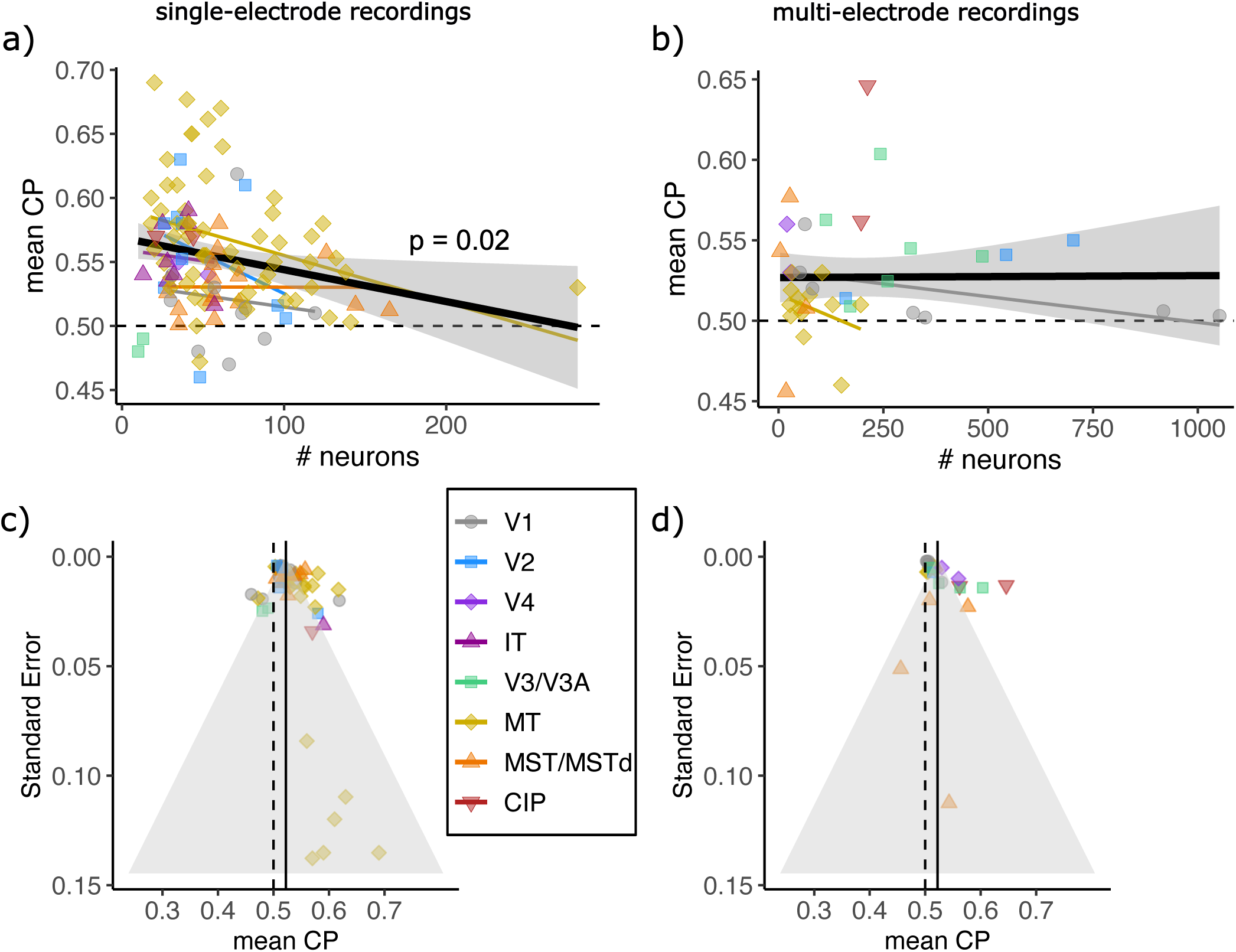
Mean CP values decrease with statistical power of the study. **(a)** Relationship between mean CP and the number of neurons contributing to the estimate, restricted to single-electrode studies to reduce confounding. A negative trend (slope = −0.0003, *p* = 0.02, *N* = 109) suggests that low-powered studies (fewer neurons) tend to report inflated CP values. **(b)** same as (a) but restricted to multi-electrode studies. No trend is observed (*p* = 0.96, *N* = 37) **(c)** Funnel plot showing standard error (y-axis) versus mean CP (x-axis). Restricted to single-electrode studies to reduce confounding. The solid vertical line marks the meta-analytic mean CP (0.52), computed as an inverse-variance-weighted average. The shaded triangle represents the 95% expected region assuming this true mean. An asymmetry is evident, with higher-error studies skewed toward elevated CP values, consistent with publication bias (Egger’s test: *p* = 0.01, *N* = 51). **(d)** same as (c) but restricted to multi-electrode studies. Similar asymmetry as for single-electrode studies is observed (Egger’s test: *p <* 0.001, *N* = 24).

**Figure S5:**
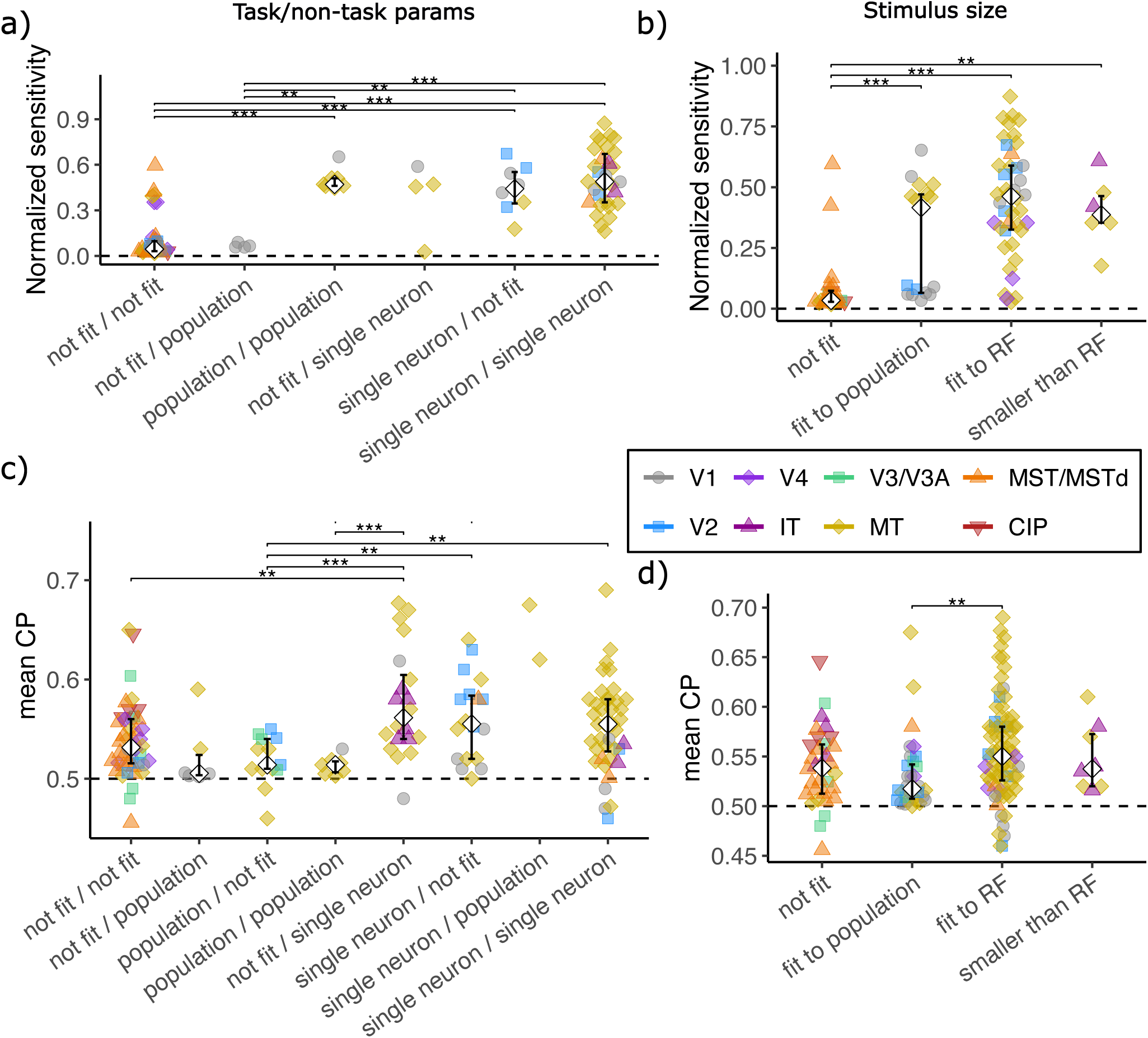
Effect of tailoring techniques on neuronal sensitivity (a, b) and mean CP (c, d). Left column (**a, c**) shows the effect of tailoring two types of stimulus parameters: task-relevant (“task”) and task-irrelevant but response-modulating (“non-task”). X-axis labels represent combinations of tailoring choices. For example, the label ‘not fit/single neuron’ indicates that the task parameter was not tailored, whereas the non-task parameter was optimized for each individual neuron. Note that in panel **a**, nearly all available sensitivity data come from single-electrode studies, so tailoring to population-level properties is mostly absent. Right column (**b, d**) shows the effect of tailoring stimulus size relative to the receptive field. Asterisks indicate significant differences between two categories based on Mann–Whitney tests (* *p <* 0.05, ** *p <* 0.01, *** *p <* 0.001). For **a, c** *p <* 0.05 results are omitted as they would not survive correction for multiple comparisons.

**Figure S6:**
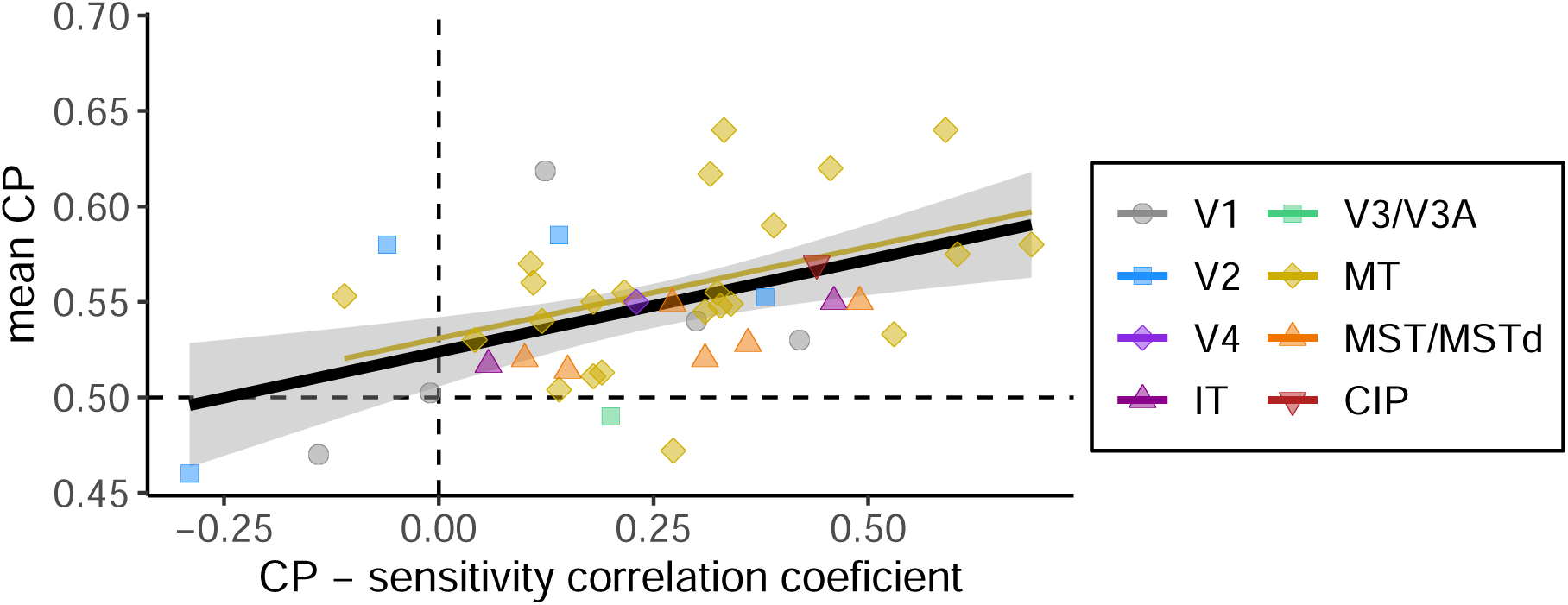
Relationship between mean CP and the within study CP–sensitivity correlation coefficient (Pearson or Spearman). For comparability, coefficients based on neuronal thresholds were sign-inverted. The black line shows the overall regression (slope = 0.10, *p* = 0.001, N = 41); the thin golden line shows the regression within MT (data from other areas were too sparse for separate analysis).

**Figure S7:**
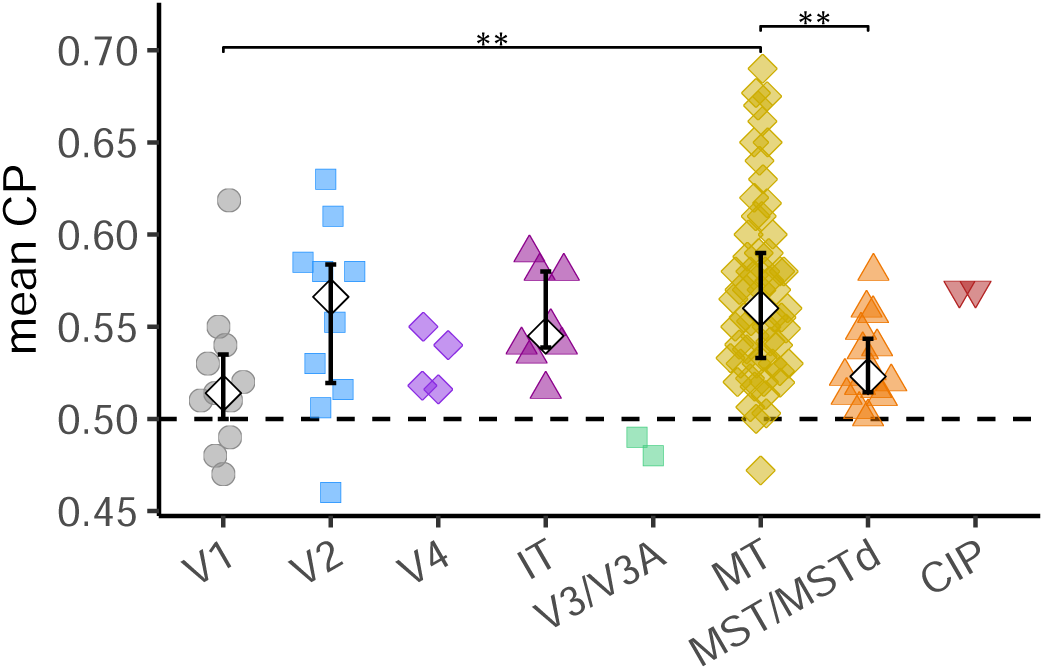
Mean CP values across brain areas for single-electrode recordings. Diamonds and error bars represent medians and interquartile ranges for areas with more than five observations. Asterisks indicate significant differences between two brain areas based on Mann–Whitney tests (** *p <* 0.01, *** *p <* 0.001); *p <* 0.05 results are omitted as they would not survive correction for multiple comparisons.

**Figure S8:**
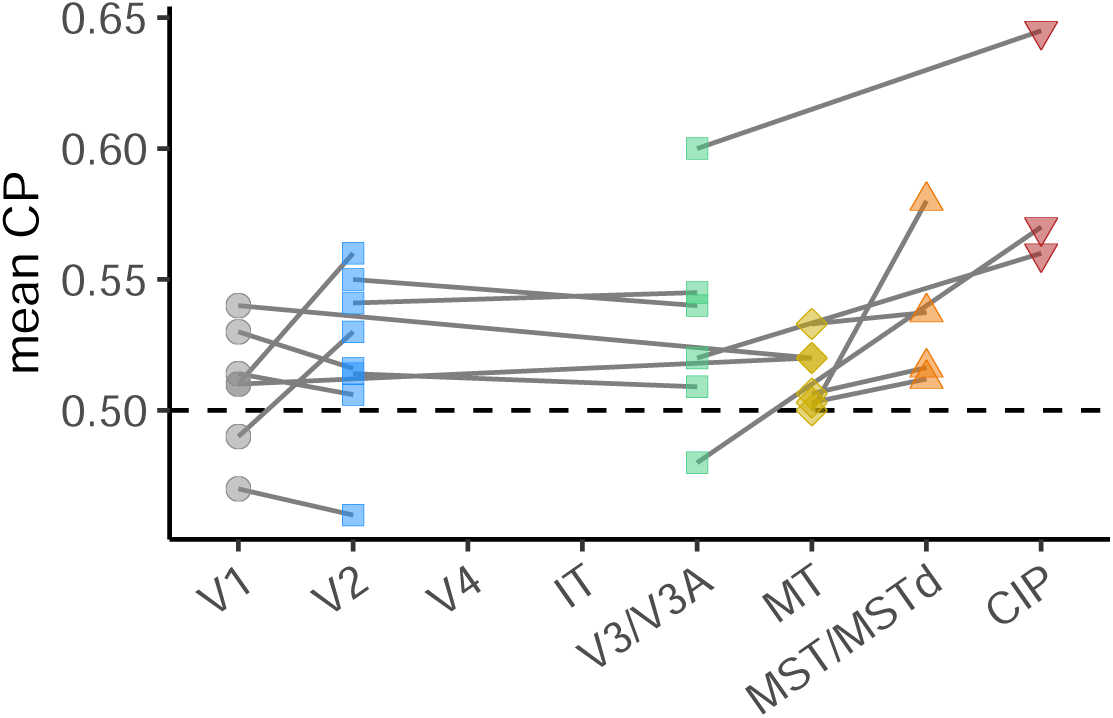
Mean CP values across brain areas within the same study and monkey. Lines connect data points from the same monkey within each study.

**Figure S9:**
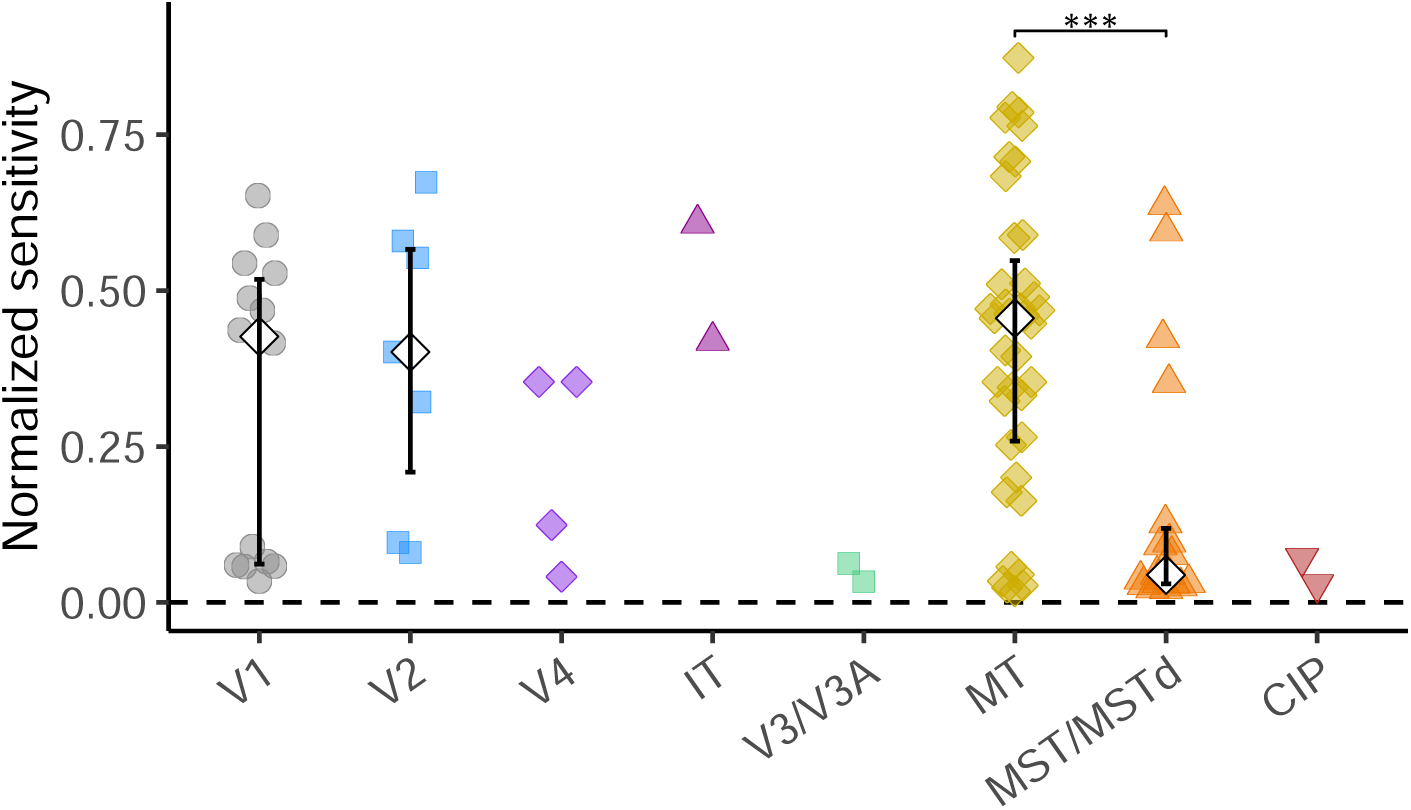
Comparison of task sensitivities between brain areas. Asterisks indicate significant differences between two brain areas based on Mann–Whitney tests (** *p <* 0.01, *** *p <* 0.001); *p <* 0.05 results are omitted as they would not survive correction for multiple comparisons.

**Figure S10:**
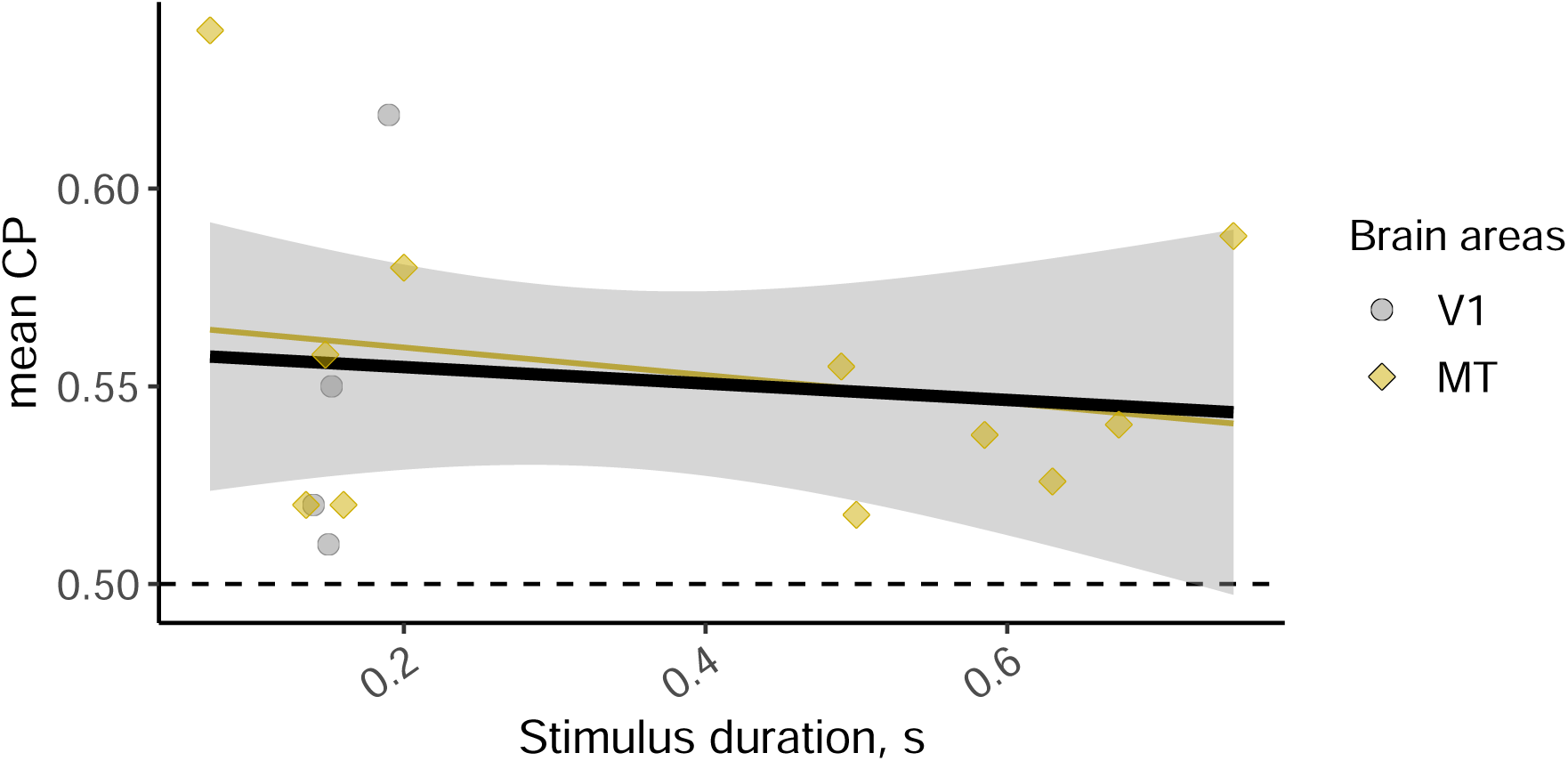
Relationship between mean CP and stimulus duration for reaction time tasks.

**Figure S11:**
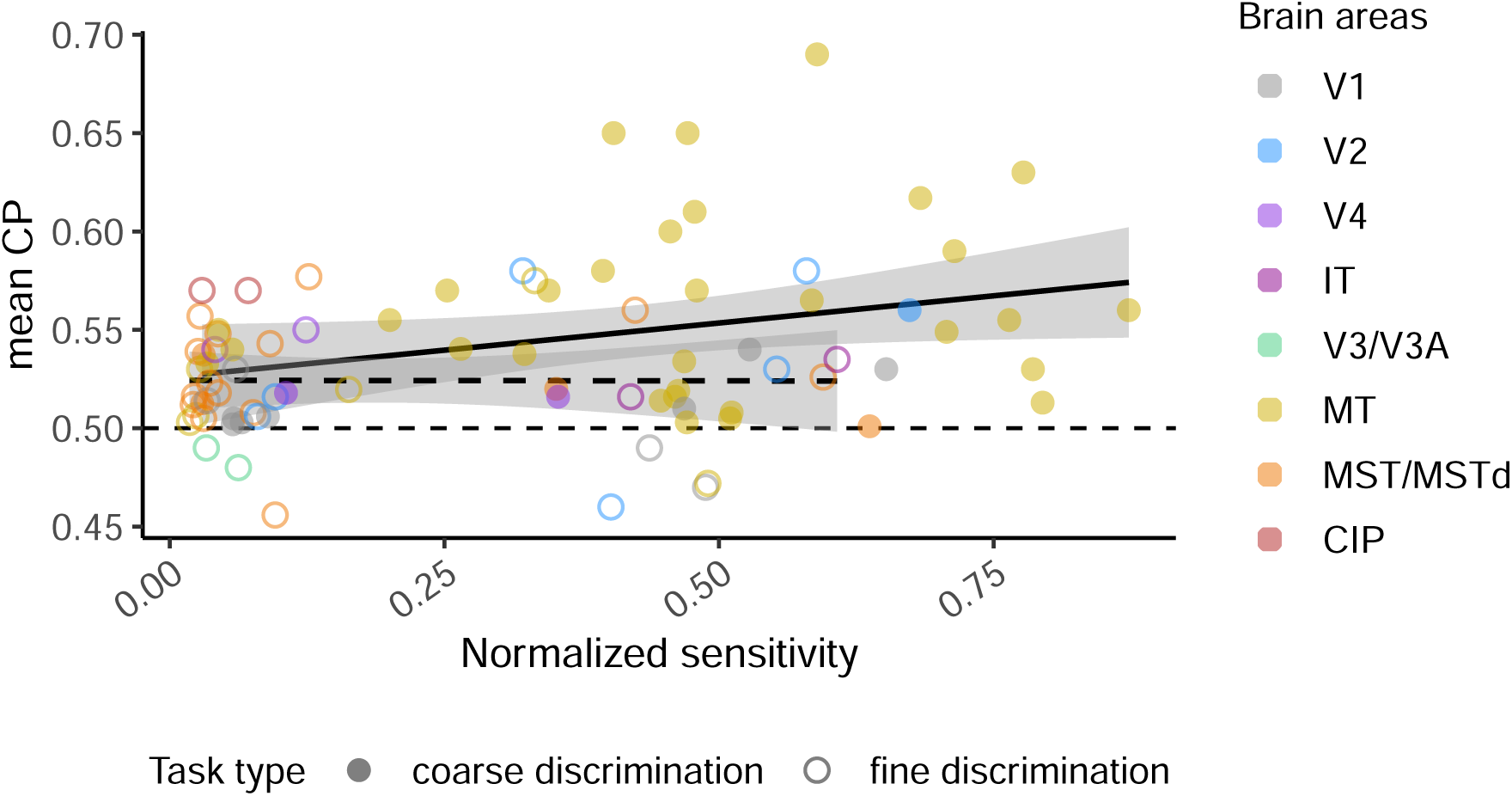
Relationship between mean CP and normalized sensitivity for fine-vs. coarse-discrimination tasks. Linear regression analyses were performed separately for the two task types: solid line – coarse- and dashed line – fine-discrimination task). A separate model with interaction terms revealed no significant difference in slope or intercept: Δ*β* = *β*_fine_ − *β*_coarse_ = −0.03, *p* = 0.3; Δ*β*_0_ = −0.007, *p* = 0.7.

**Figure S12:**
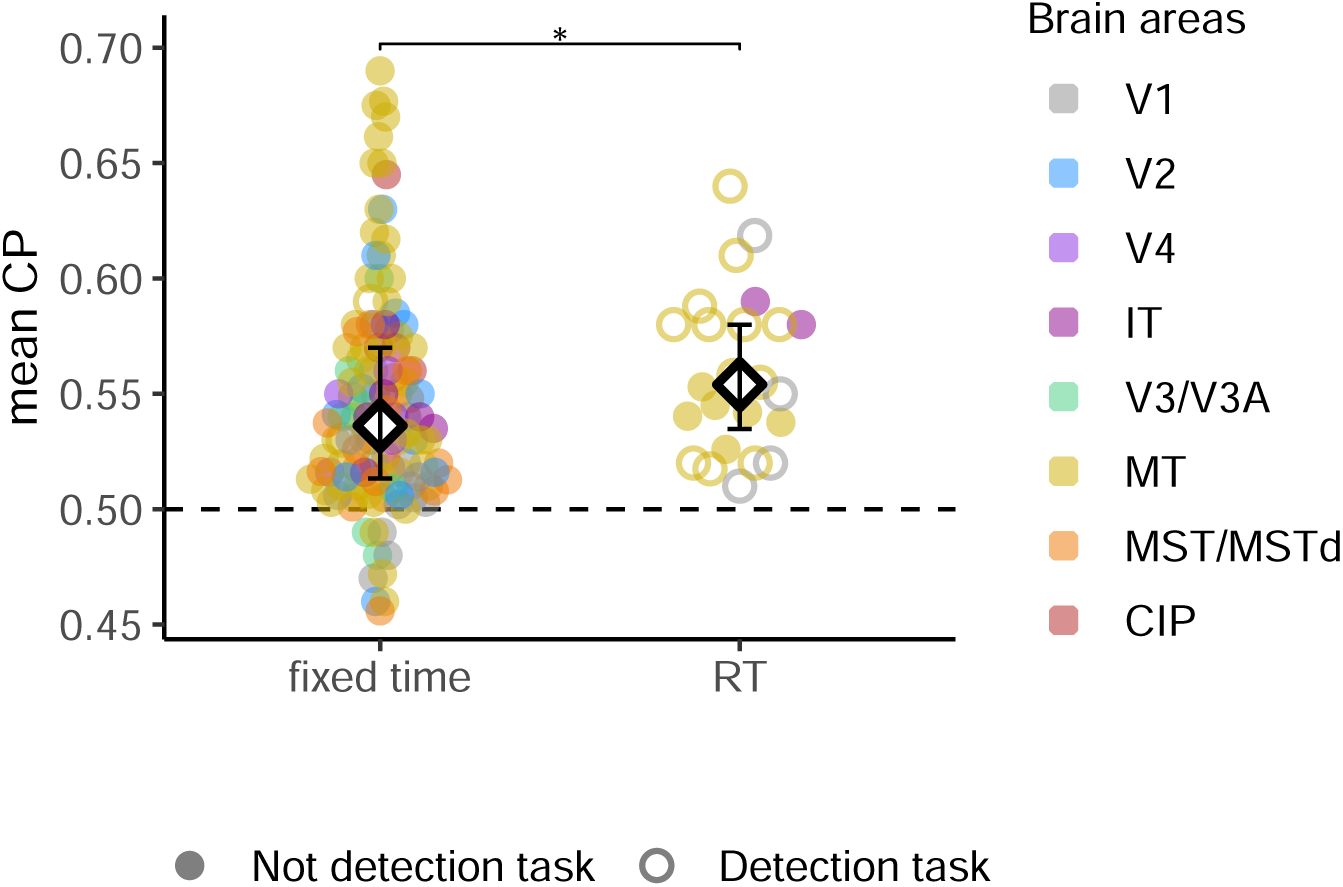
Relationship between Mean CP values with whether experiment have reaction-time design. The empty circles represent detection tasks; solid circles – all the other task types. In RT tasks (N = 24), 16 data points are from the detection tasks, 8 –from coarse-discrimination ones.

**Figure S13:**
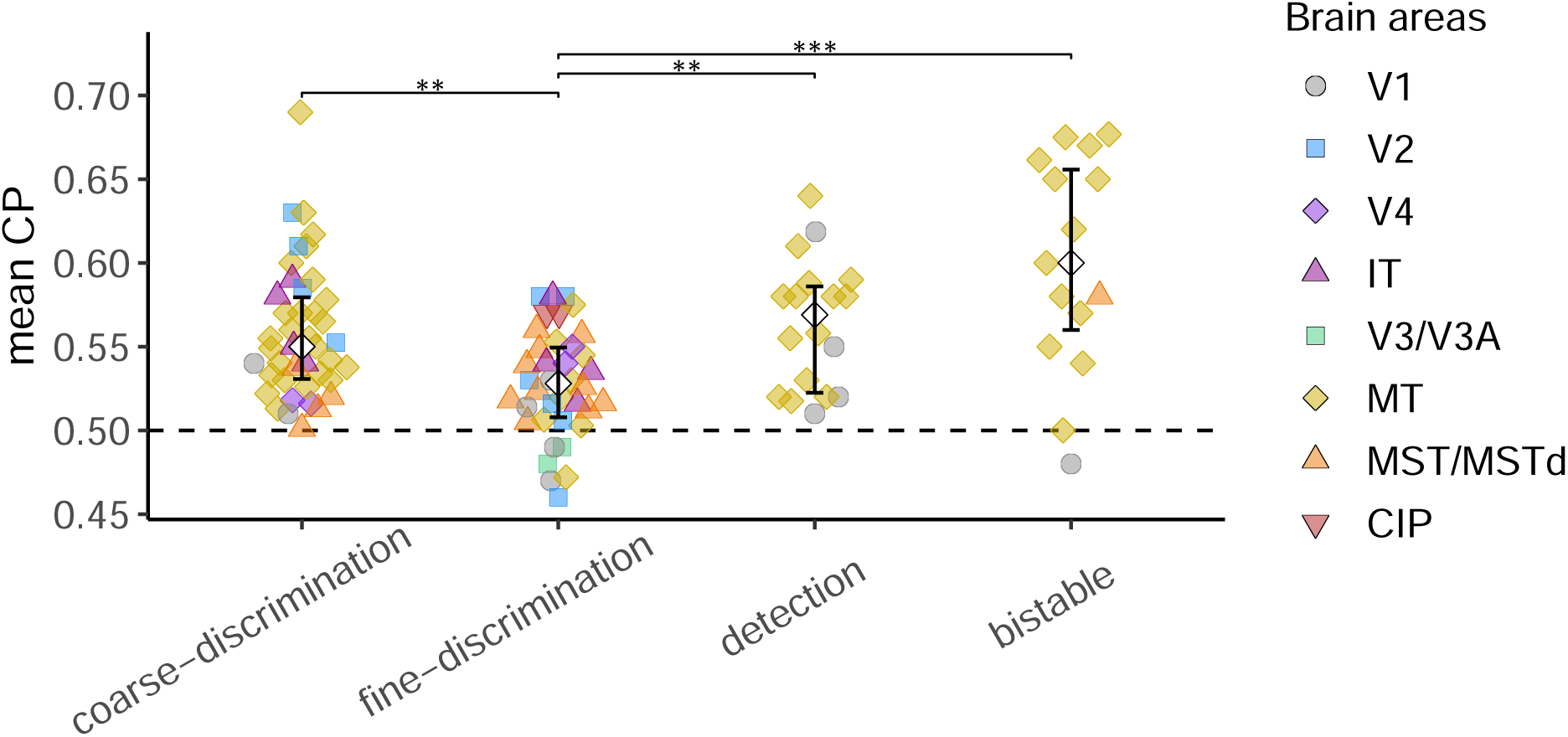
Mean CP values across task types for single-electrode recordings. Diamonds and error bars represent medians and interquartile ranges. Asterisks indicate significant differences between two task types based on Mann–Whitney tests (** *p <* 0.01, *** *p <* 0.001).

**Figure S14:**
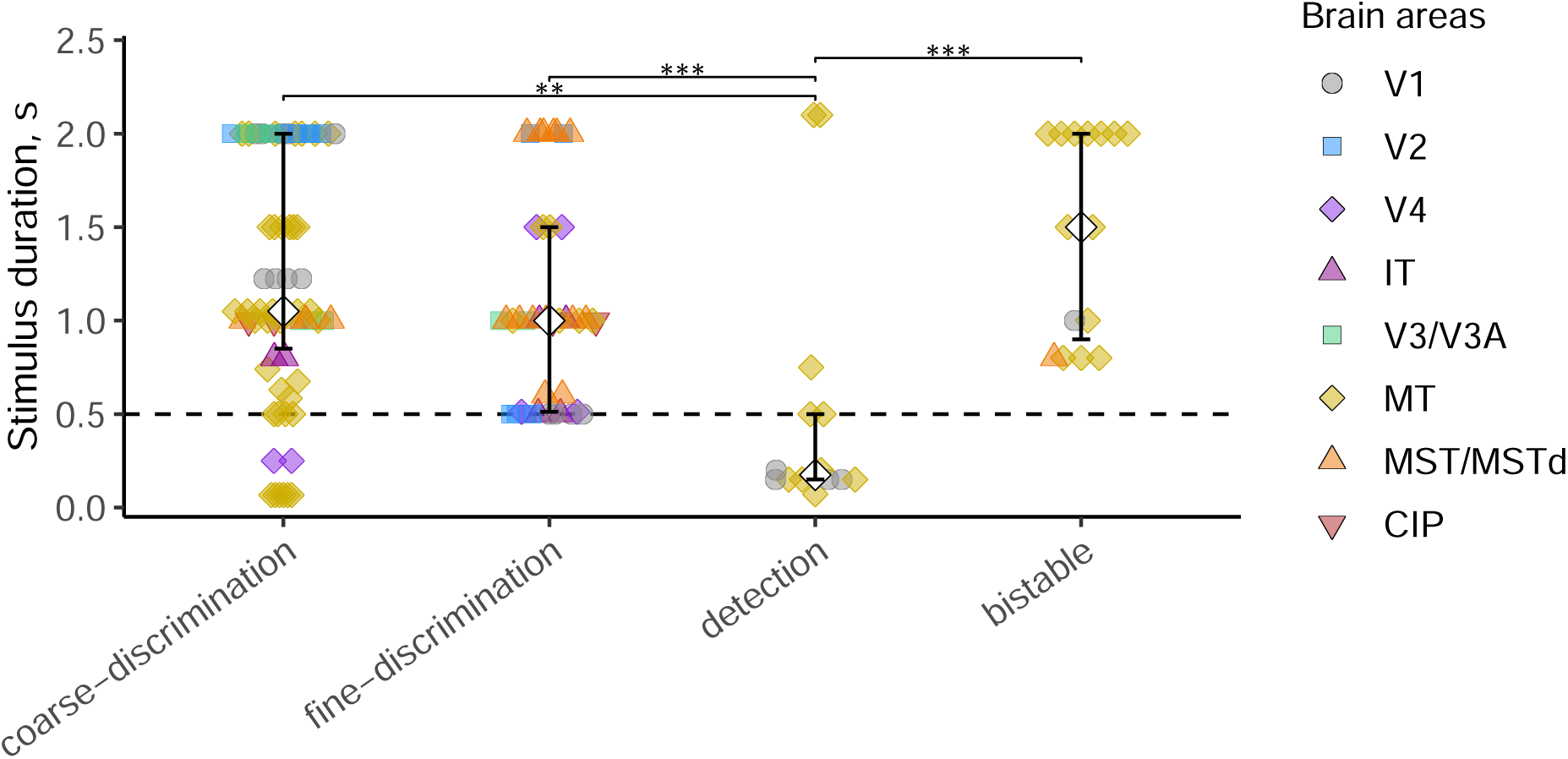
Relationship between stimulus duration and task type. Asterisks indicate significant differences between two task types based on Mann–Whitney tests (** *p <* 0.01, *** *p <* 0.001).

**Figure S15:**
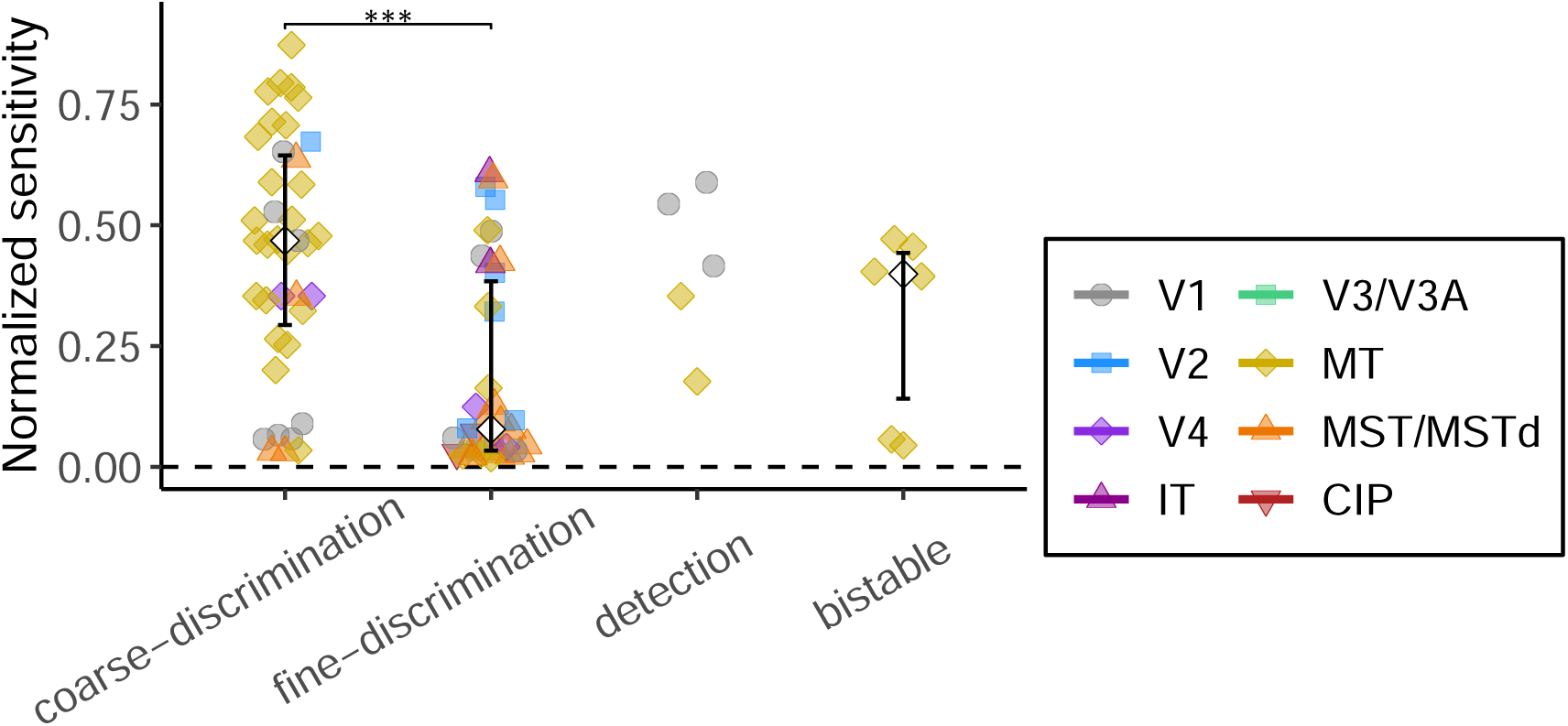
Fine-discrimination tasks show substantially lower sensitivity, which may account for their reduced mean CP. Normalized neuronal sensitivity (inverse mean N/P ratio) across task types. Diamonds and error bars represent medians and interquartile ranges with more than five observations. Asterisks indicate significant differences between two task types based on Mann–Whitney tests (** *p <* 0.01, *** *p <* 0.001).

**Figure S16:**
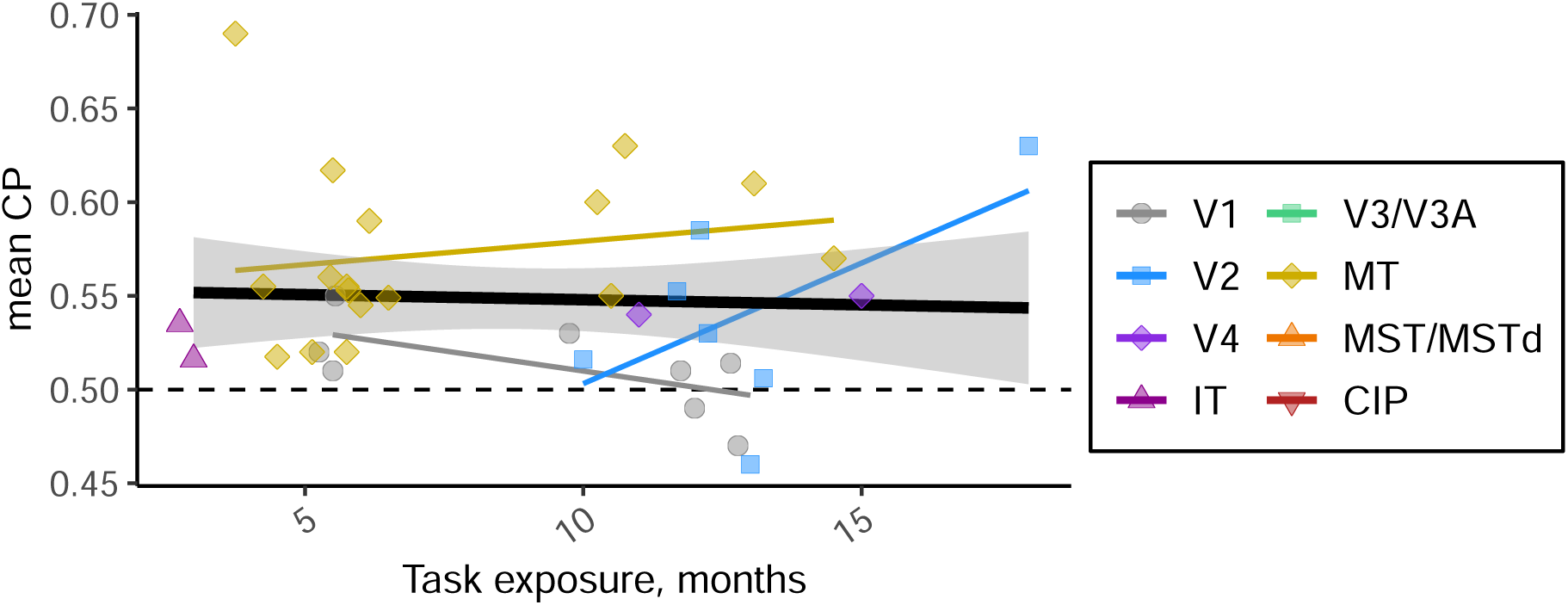
Relationship between mean CP values and task exposure.

**Figure S17:**
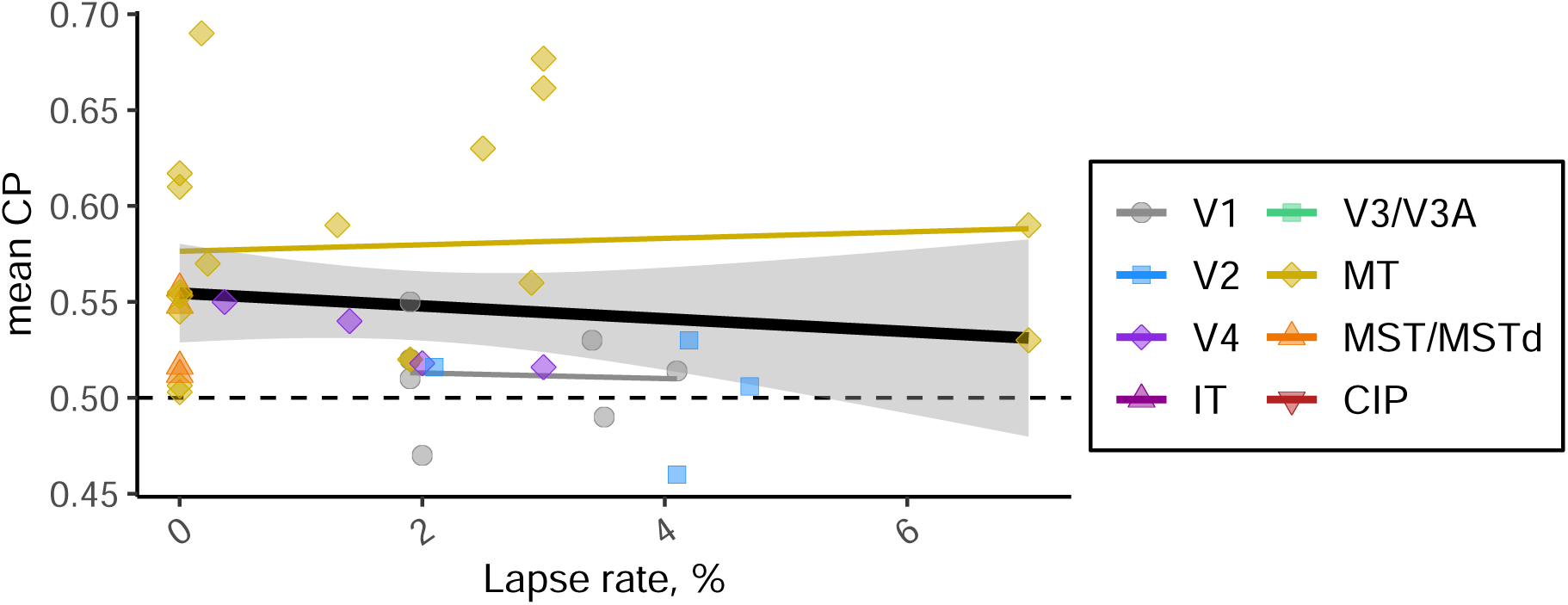
Relationship between Mean CP values and monkey’s lapse rate.

**Figure S18:**
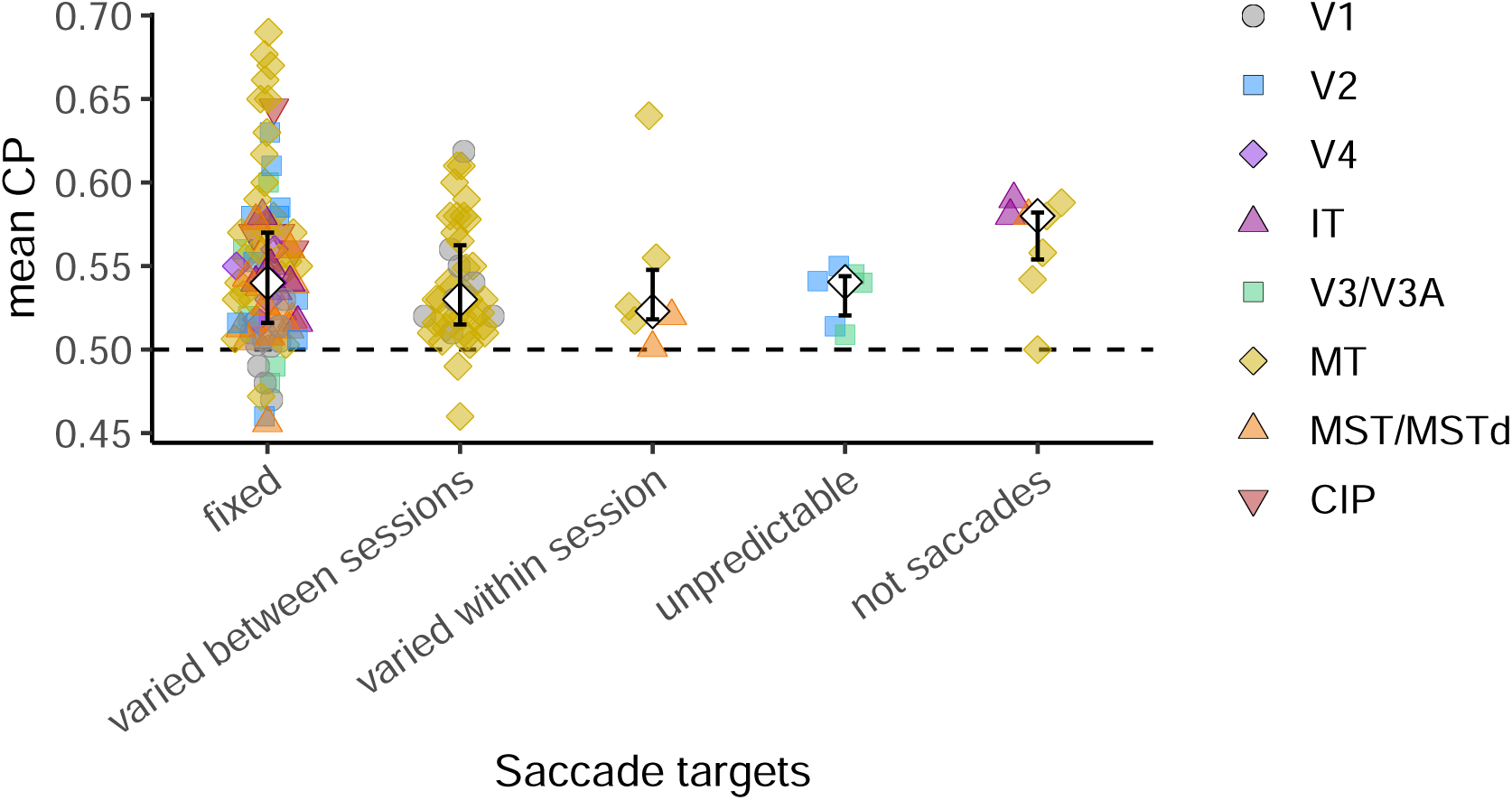
Relationship between Mean CP values and predictability of targets. No pairwise comparisons were statistically significant (Mann-Whitney test, corrected for multiple comparisons).

**Figure S19:**
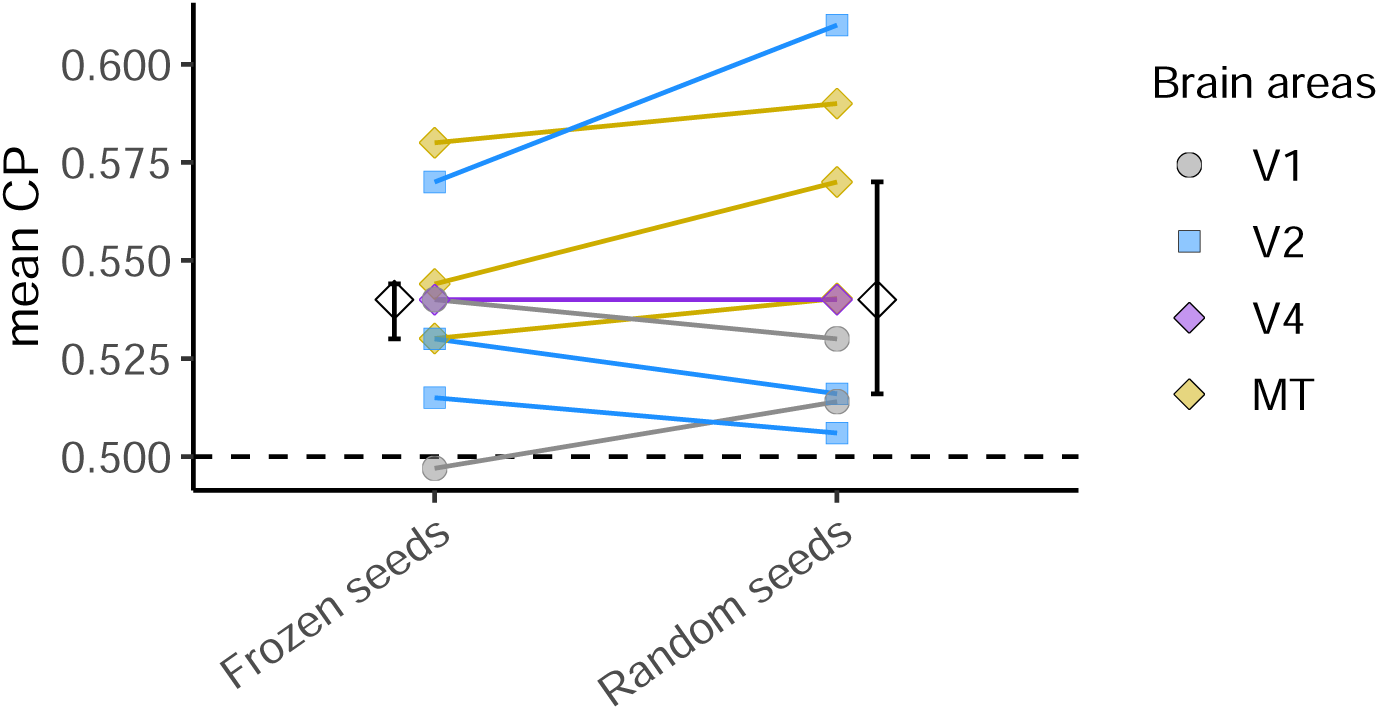
Relationship between mean CP values and stimulus noise structure. Mean CP values are shown for conditions using frozen vs. random stimulus seeds. Lines connect data points from the same study. Wilcoxon signed-rank tests have not revealed statistical difference (*p* = 0.3).

**Figure S20:**
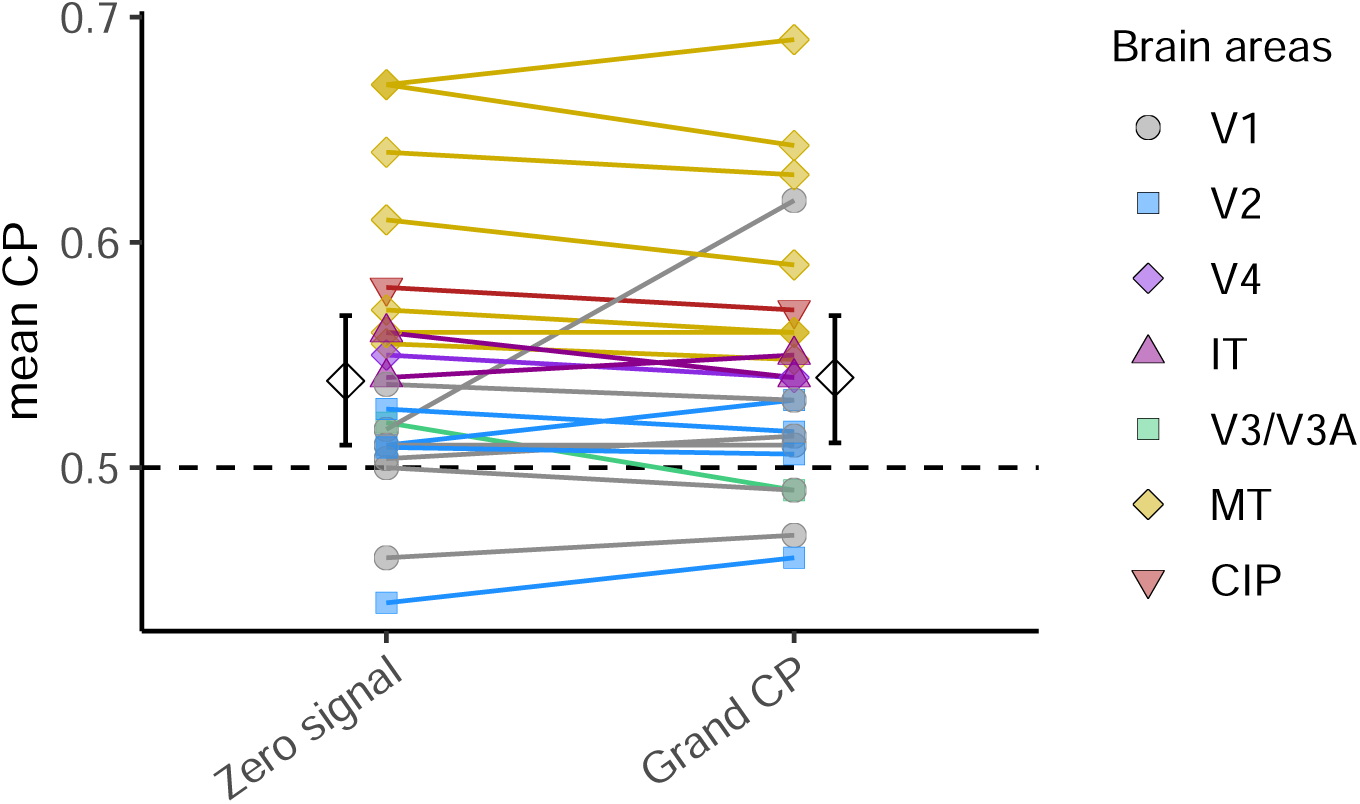
Relationship between mean CP values and CP estimation method. Lines connect data points from the same study. Wilcoxon signed-rank test has not revealed statistical difference (*p* = 0.4)

**Figure S21:**
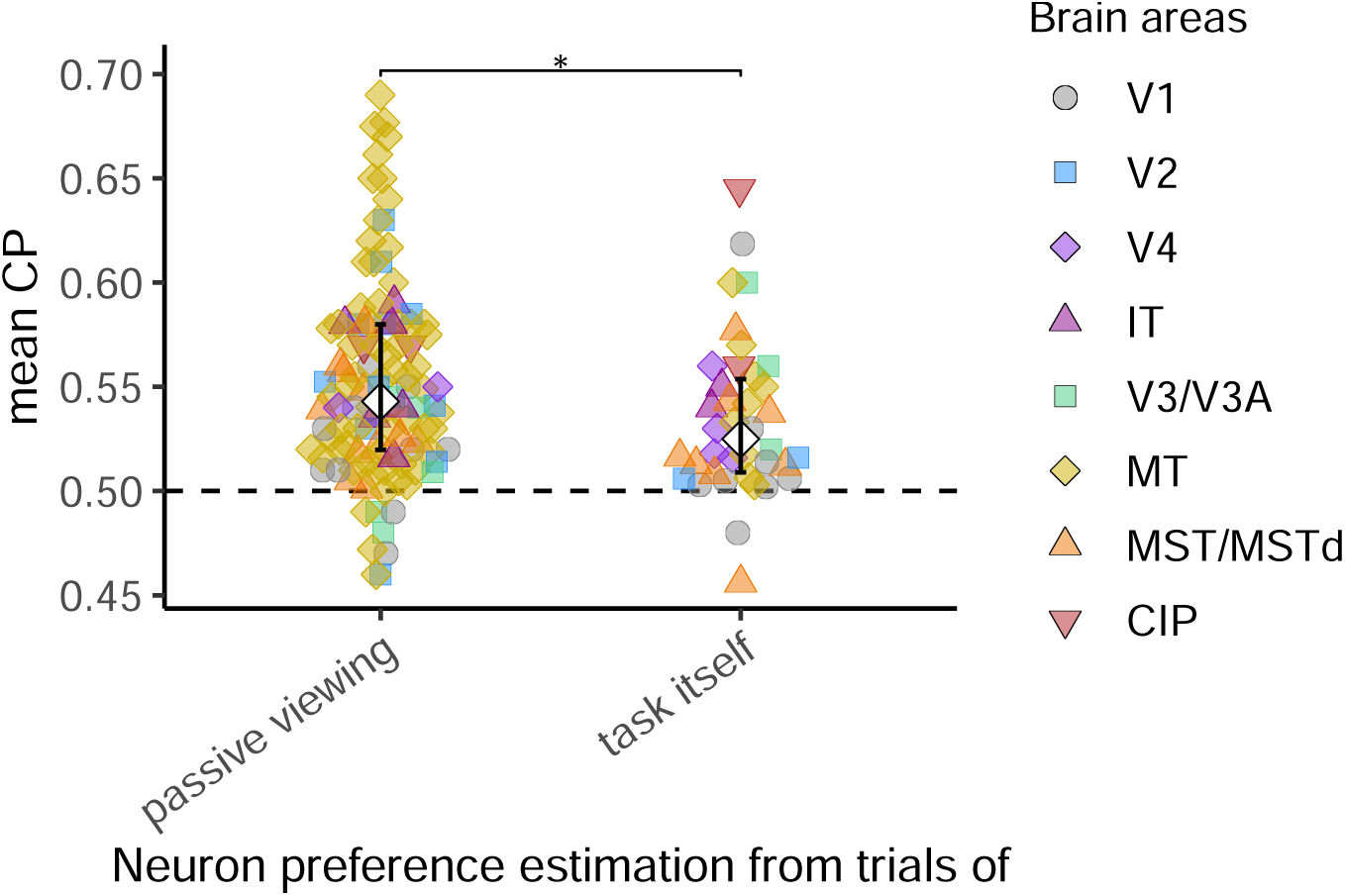
Relationship between Mean CP values and method of neuron preference estimation. Mann–Whitney test revealed that studies with preferences estimated from the task trials exhibited lower mean CP compared to those that used preferences estimated from passive viewing sessions (* asterisk in the figure, *β* = −0.018, *p* = 0.03), but this effect appears to be driven by other factors (see section 8.3).

**Figure S22:**
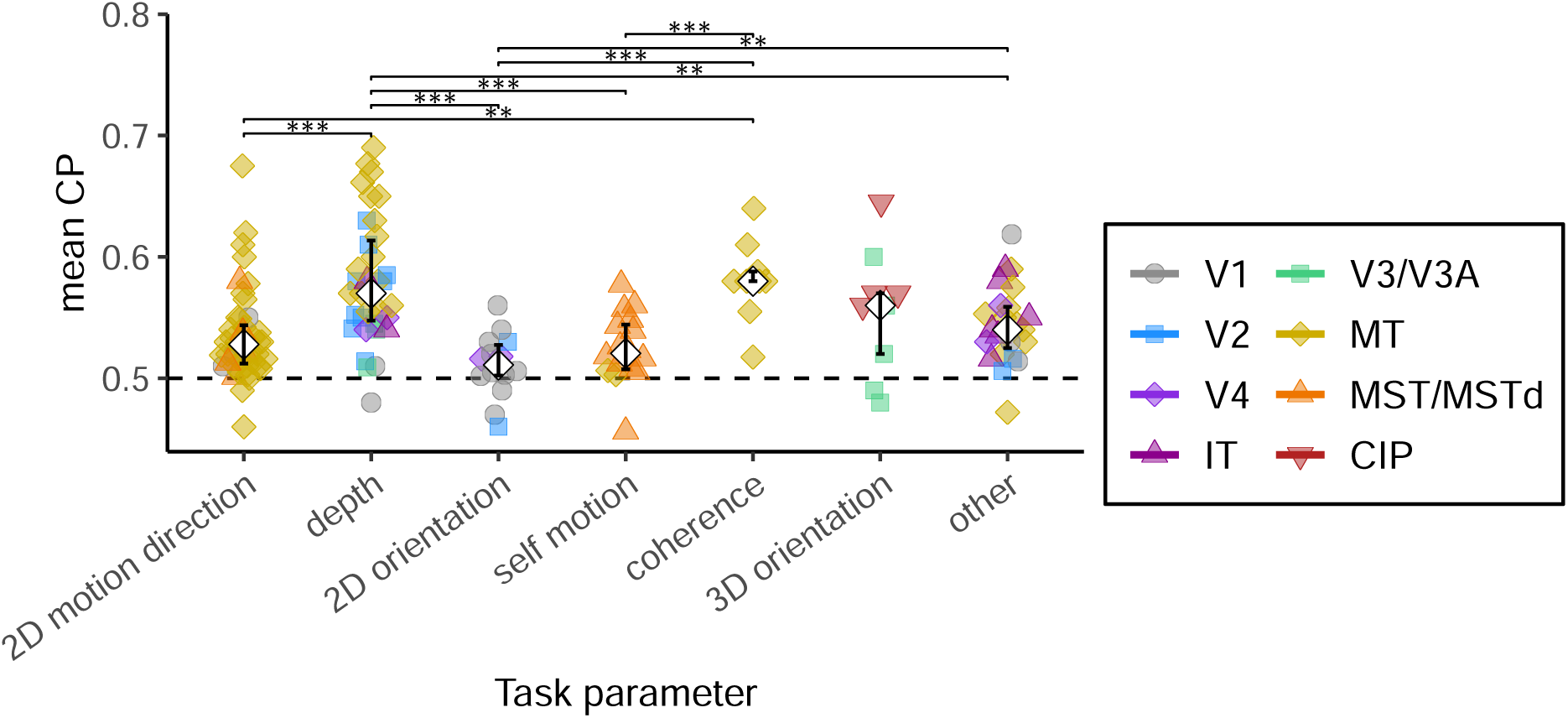
Relationship between Mean CP values and task variable. Asterisks indicate significant differences between two task variables based on Mann–Whitney tests (** *p <* 0.01, *** *p <* 0.001); *p <* 0.05 results are omitted as they would not survive correction for multiple comparisons. Note that the observed difference is largely driven by the effect of brain area, as task characteristics and brain area are strongly coupled.

**Figure S23:**
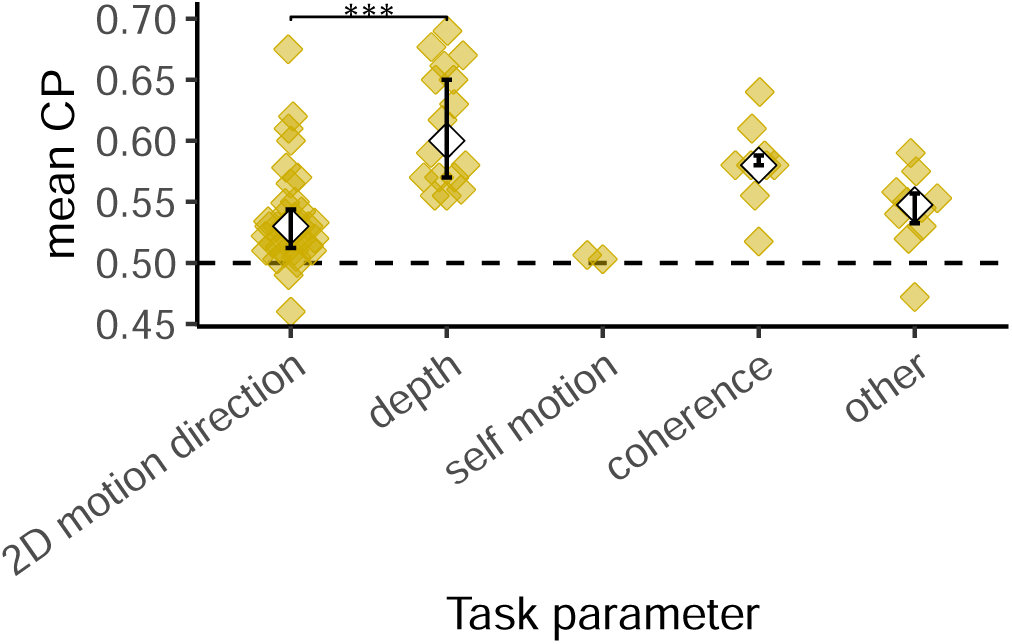
Relationship between mean CP values and task variable within area MT. Asterisks indicate significant differences between task conditions based on Mann–Whitney tests (*** *p <* 0.001); results with *p <* 0.05 are omitted as they would not survive correction for multiple comparisons. Unlike the across-area analysis, the observed difference between motion and depth discrimination tasks in MT cannot be attributed to brain area and remains significant after controlling for sensitivity, stimulus duration, and task type (see section 8.3).

**Figure S24:**
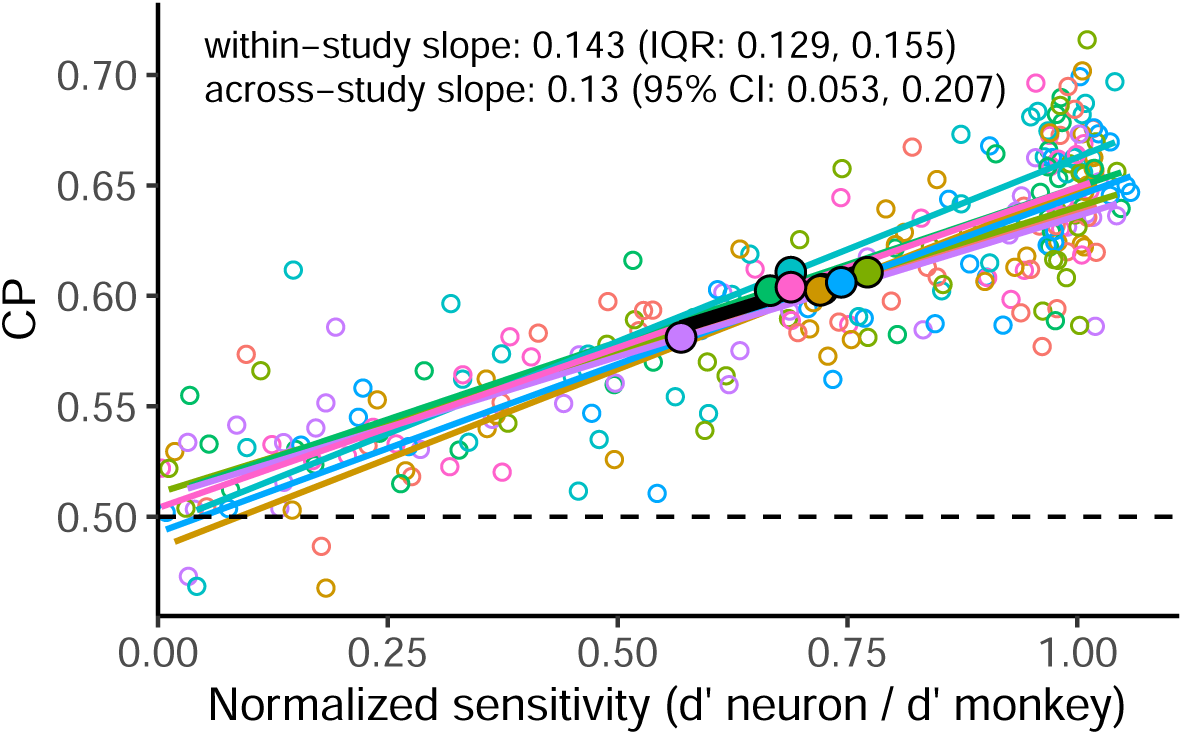
Hierarchical inference model simulations with task-aligned feedback and incomplete learning predict within- and across-study CP–normalized sensitivity slopes of approximately 0.15. Normalized sensitivity (x-axis) is defined as the *d^*′*^_neuron_*/ *d*^*′*^_*behavior*_, where *d*^′^_neuron_ is estimated during task performance and reflect both feedforward and feedback influences (see Section 2.4.1). Simulations utilized the hierarchical inference model of Haefner et al. (2016) with 256 sensory neurons randomly divided into 8 “studies” of 32 neurons each (indicated by color). Feedback strength was controlled by the parameter *δ* and set to 0.08—the maximum value used in Haefner et al. (2016)—which corresponds to proficient but incomplete task learning. Small open circles represent individual neurons, and large filled circles denote study means. Thin lines indicate within-study regressions; the thick black line represents the across-study regression.

**Figure S25:**
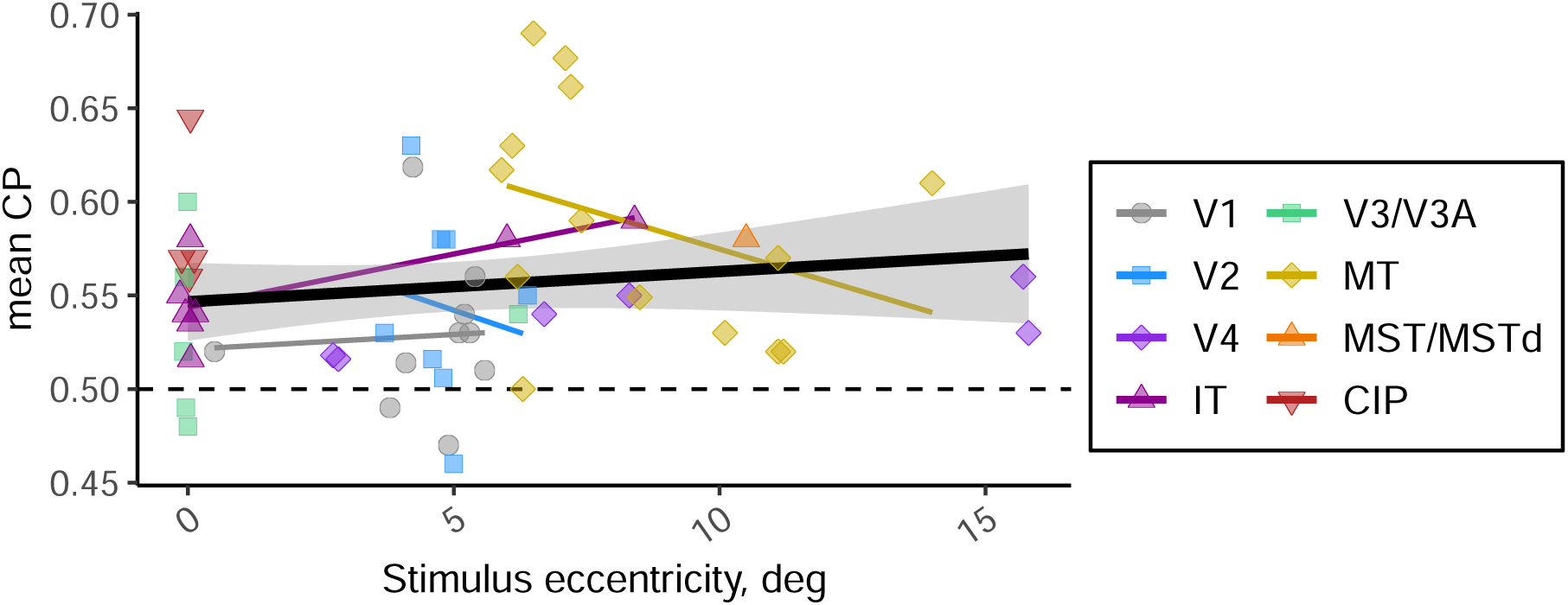
Relationship between mean CP values and stimulus eccentricity.

**Figure S26:**
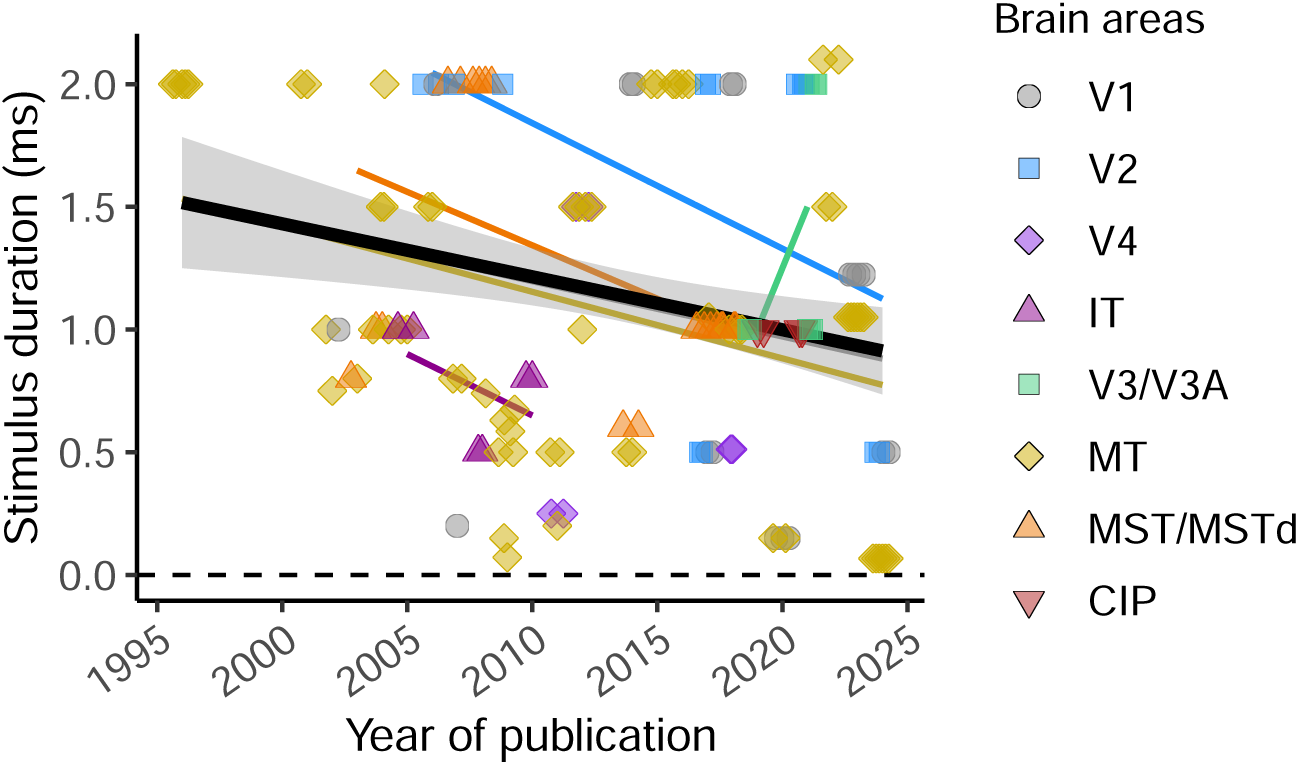
Relationship between stimulus duration and year of publication.

**Figure S27:**
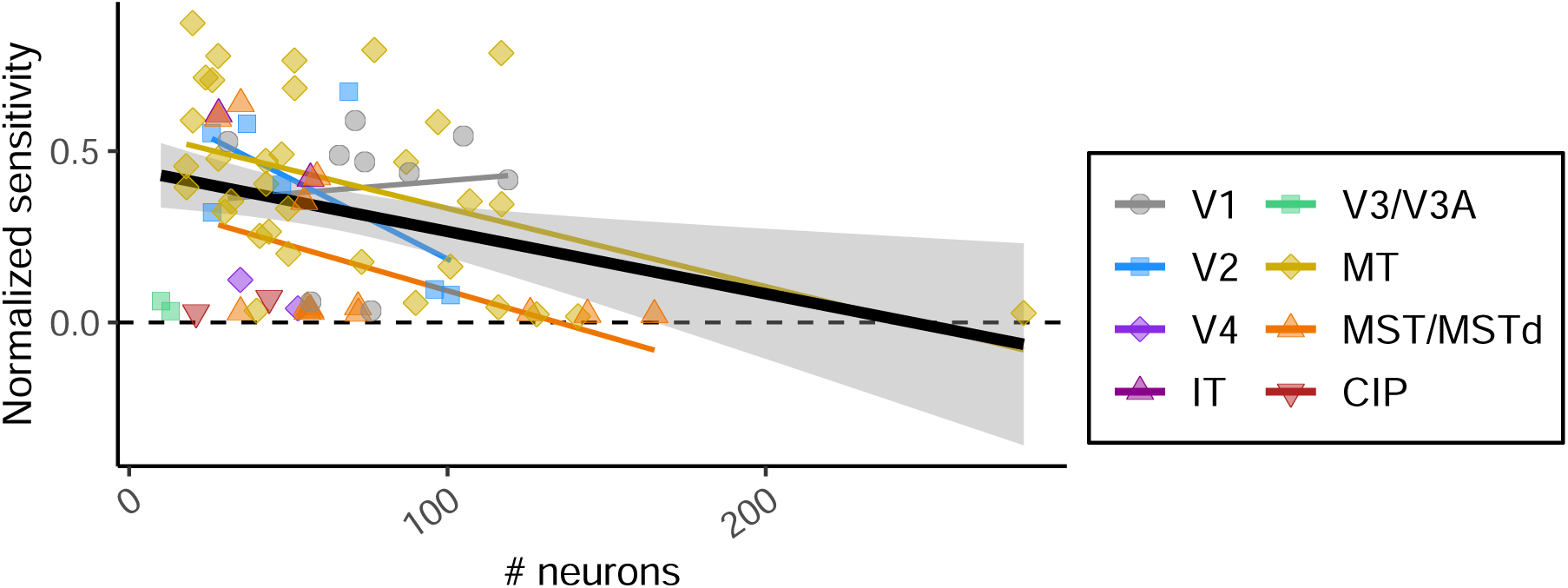
Relationship between neural sensitivity and number of neurons recorded in the study (only single electrode recordings).

**Figure S28:**
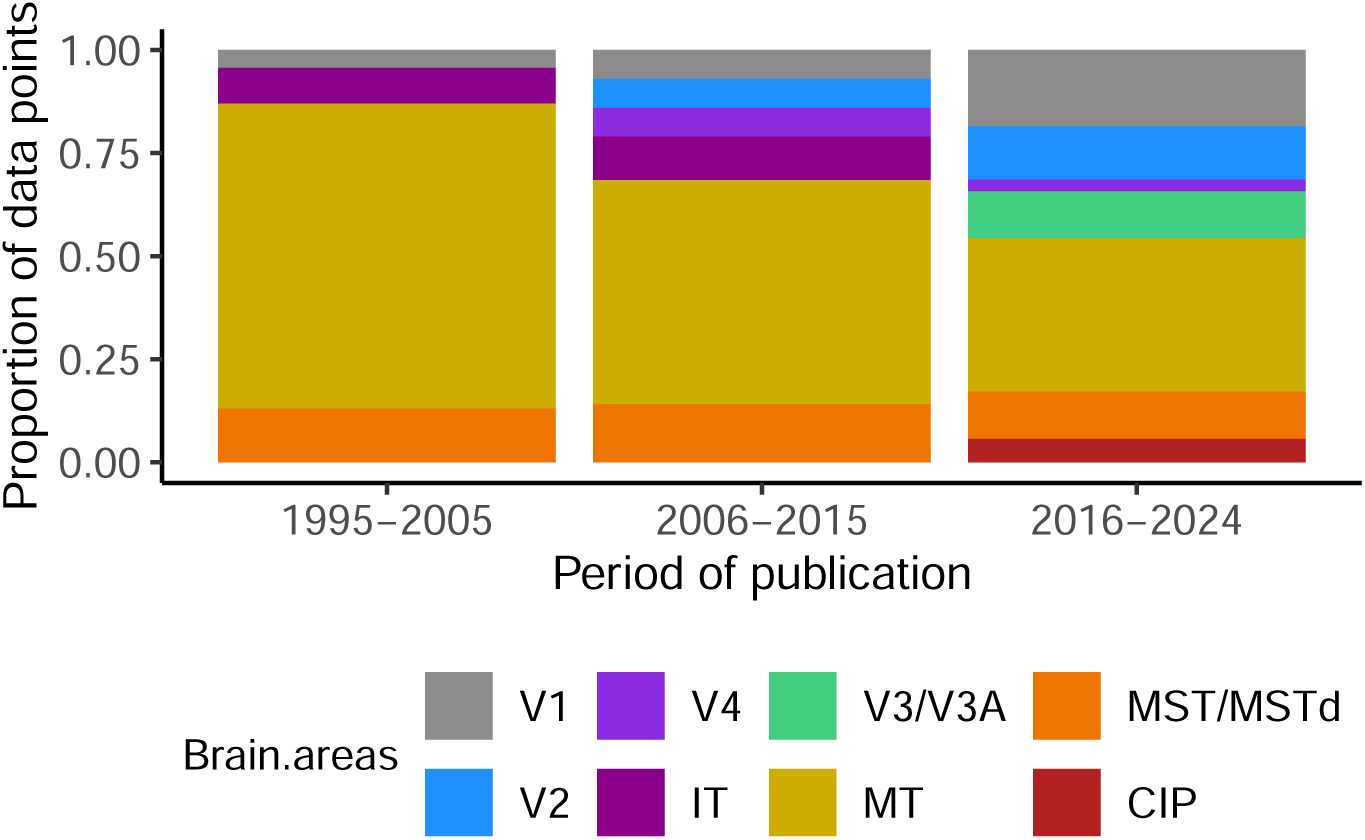
Changes in the distribution of brain areas studied over time. Proportion of data points from each brain area is shown across three publication periods, illustrating shifts in focus within the CP literature.

### 8.3 Variables for which no effect on mean CP was found

Both feedforward and feedback models predict that task exposure (the cumulative duration the animal has performed the task) and performance metrics should positively correlate with mean CP. Longer exposure to the task may increase CP by aligning the animal’s strategy with the experimental paradigm and inducing perceptual learning (Law and Gold, 2008; Uka et al., 2012; Sanayei et al., 2018). However, we found no significant effect of task exposure on mean CP – neither in a regression including only task exposure (*p* = 0.8, *N* = 36, Fig. S16) nor in models controlling for sensitivity, brain area, stimulus duration, and task type (inverse mean N/P ratio control: *p* = 0.2, *N* = 27; tailoring control: *p* = 0.97, *N* = 33). Performance metrics that capture how well the monkey engages with the task are also expected to correlate positively with CP. The lapse rate – defined as the error rate on the easiest trials – is widely used as a proxy for such engagement (Prins, 2012; McCarley and Yamani, 2021). A negative correlation with CP was expected (Uka et al., 2012), but no significant relationship was found before (Fig. S17; *p* = 0.5, *N* = 37) and after controlling for four experimental variables (inverse N/P ratio control: *p* = 0.5, *N* = 30; tailoring control: *p* = 0.6, *N* = 35).

Recent work suggests that CP may also be influenced by the spatial arrangement and predictability of saccade targets, through the effects of saccade planning (Laamerad et al., 2024; Zhang and Gu, 2026) (bottom right of Fig. S1). A necessary precondition for the saccade-planning effects is the spatial predictability of saccade targets (Zhang and Gu, 2026). We graded saccade-target predictability across studies on a five-level scale: (1) responses not made by saccades; (2) unpredictable targets; (3) targets varying within a recording day; (4) targets varying between days; and (5) fixed targets across the entire experiment. We found no evidence that target predictability modulated mean CP, but the small number of studies with low target predictability (no saccades: *N* = 8; unpredictable targets: *N* = 6) limits the strength of this conclusion (Fig. S18).

Researchers have argued that computing CP using trials with non-identical stimuli may introduce confounds that inflate CP estimates (Nienborg and Cumming, 2009; Kang and Maunsell, 2012; Wimmer et al., 2015) (bottom left of Fig. S1). Ideally, CP should be calculated using trials in which the stimulus is fixed and fully ambiguous to avoid contamination from stimulus-driven activity (Britten et al., 1996). In practice, however, most CP studies deviate from this ideal. First, instead of presenting the exact same stimulus across trials (same noise seed, “frozen noise”), researchers often use multiple random noise seeds to prevent monkeys from memorizing specific noise patterns in the stimulus. Following Britten et al. (1996), researchers assumed that the monkey’s choice and the neuron’s response are not systematically affected by differences in the noise seed. However, Nienborg and Cumming (2009) and Wimmer et al. (2015) showed that this practice can lead to a modest overestimation of CP. Second, most studies compute what is known as “grand CP”, in which neuronal responses are z-scored within each stimulus condition and then pooled across all conditions. This approach substantially increases the number of trials available for CP estimation compared to analyses restricted to ambiguous stimuli (“zero signal” in Fig. 2) alone. It rests on the assumption that z-scoring effectively removes all task-relevant stimulus information from the neural responses – otherwise, CP may be artificially inflated due to a stimulus-choice correlation. Conversely, Kang and Maunsell (2012) showed that when choice distributions are highly imbalanced across conditions, z-scoring can underestimate CP by subtracting away part of the choice-related signal. In our meta-analysis, we used grand CP with random seeds whenever possible, as it was the most commonly reported metric across studies (63% of data points, see Section 7). For the subset of studies that reported multiple CP metrics, we directly compared the different methods. We found no significant difference between frozen and random seed conditions (Wilcoxon signed-rank test: *p* = 0.2, *N*_pairs_ = 9; Fig. S19), nor between grand CP and CP calculated from ambiguous stimulus trials (*p* = 0.5, *N*_pairs_ = 22; Fig. S20). Overall, our results suggest that while CP calculation methods may introduce some variability, in practice they have little effect on mean CP compared to the much greater uncertainty arising from limited trials and neurons.

If a neuron’s stimulus tuning is partially driven by feedback signals, estimating its stimulus preference from task trials can increase CP estimates, particularly for low-sensitivity neurons (Zaidel et al., 2017). In CP calculations, the “positive” choice is defined relative to the neuron’s preferred stimulus category (see Fig.2). Zaidel et al. (2017) and Elmore et al. (2019) showed that choice-related signals during task performance can shift estimated preferences toward the direction of choice modulation and flip some CPs from below 0.5 to above 0.5, thereby increasing population mean CP. We found no evidence for such an effect; rather, the 26% of data points from studies that derived preferences from task trials exhibited slightly lower mean CP (*β* = −0.017, *p* = 0.05; Fig. S21), a trend that became non-significant when controlling for recording technique (*β* = −0.012, *p* = 0.16) or when adjusting for sensitivity, brain area, duration, and task type (inverse mean N/P ratio: *β* = 0.015, *p* = 0.19; tailoring: *β* = −0.004, *p* = 0.7).

### 8.4 Variables that do not have good theoretical explanation

The apparent effect of recording technique on mean CP is fully accounted for by differences in sensitivity, brain area, stimulus duration, and task type. Of the 70 data points reporting a mean N/P ratio, only one came from a study using multi-electrode recordings, making it infeasible to assess the recording technique effect using this sensitivity measure. When controlling for tailoring technique alone, the effect of multi-electrode recordings on mean CP was not statistically significant (*β*_multi-electrode_ = 0.004, *p* = 0.8), and it remained non-significant after additionally controlling for brain area, stimulus duration, and task type (*β*_multi-electrode_ = 0.007, *p* = 0.6). Our results suggest that the apparent effect of recording technique on mean CP is driven by tailoring limitations, as multi-electrode studies cannot fine-tune stimuli to individual neurons.

Similarly, the relationships of mean CP with year of publication and number of neurons – patterns suggestive of publication bias – can instead be explained by four key experimental factors. When controlling for neuronal sensitivity, brain area, stimulus duration, and task type, the effect of publication year disappears entirely (inverse mean N/P ratio: *β*_year_ = 0.0005, *p* = 0.55; tailoring-based sensitivity: *β*_year_ = − 0.0003, *p* = 0.68). This pattern reflects a temporal decline in both sensitivity and stimulus duration, along with a shift in brain areas studied – from MT to V1 in particular (Fig.3c, S26, S28). The negative association between mean CP and the number of neurons loses statistical significance when accounting for the four variables (single-electrode studies only; inverse mean N/P ratio: *p* = 0.15; tailoring: *p* = 0.12). This attenuation appears primarily driven by a decrease in neuronal sensitivity in studies that recorded from more neurons, as such studies often reduce tailoring efforts due to practical constraints (single-electrode studies only, Fig.S27).

Stimulus eccentricity – the distance between the stimulus and the fixation point – showed no significant relationship with mean CP in any analysis (Fig. S25; pairwise regression: *p* = 0.32, *N* = 57; regression controlling for sensitivity using the inverse N/P ratio: *p* = 0.5, *N* = 35; tailoring-based sensitivity control: *p* = 0.3, *N* = 57).

### 8.5 Studies with exceptionally high or low mean CP values not explained by experimental variables

We examined data points that appeared as outliers—defined as having the largest residu-als—in our main regression models that included neuronal sensitivity (either inverse mean N/P ratio or tailoring-based proxy), brain area, stimulus duration, and task type.

Several studies demonstrated pronounced heterogeneity between monkeys, with mean CP values for one animal showing extreme deviations from model predictions. Britten et al. (1996) reported a relatively low mean CP in MT for one of four monkeys (monkey J: mean CP = 0.51, prediction error in model with inverse mean N/P ratio = –2.2 SD), despite a large sample size (*N* = 77; CP values for the other monkeys: 0.53, 0.57, 0.58). An even more striking outlier comes from Goris et al. (2017), who reported a mean CP below 0.5 and a negative CP–sensitivity relationship in V2 for one monkey (monkey 1: mean CP = 0.46, *N* = 48, prediction error in model with inverse mean N/P ratio = –1.8 SD; model with tailoring = –1.9 SD), in contrast to the other monkey in the study (mean CP = 0.53). Uka and DeAngelis (2006) observed a similar pattern in area MT: one monkey (monkey B) had a noticeably low mean CP of 0.47 (*N* = 48, prediction error in model with inverse mean N/P ratio = –2.6 SD; in model with tailoring = –2.1 SD), whereas the other monkey exhibited a mean CP of 0.57, a value typical for this brain area. Such pronounced inter-animal variability suggests that individual task strategies or the sampled neuronal populations are more heterogeneous than is assumed by theoretical models.

Some studies using bistable stimuli reported either unexpectedly high or low mean CP values. Maier et al. (2007) reported high CPs in MT with binocular motion rivalry stimuli, despite using much shorter duration of 800 ms compared to the 2000 ms typical for rotating cylinder tasks and tailoring stimulus size for a multi-neuron population rather than individual units (two monkeys combined: mean CP = 0.68, prediction error in model with tailoring = +2.1 SD). In this study, monkeys did not directly report their percepts; instead, perception was inferred via binocular flash suppression—a technique where presenting a pattern to one eye and subsequently flashing a dissimilar pattern to the other reliably biases perception toward the newly flashed stimulus. By comparison, Williams et al. (2003) found no significant CP in MT using a bistable apparent motion stimulus where evenly spaced columns of dots abruptly jump by half their inter-column distance, creating an illusion of movement in one of two opposite directions (two monkeys combined: mean CP = 0.50, prediction error in model with tailoring as sensitivity control = –2.8 SD). The authors argued that, unlike the rotating cylinder, their stimulus was suboptimal for MT and may have engaged higher-order areas. Similarly, Grunewald et al. (2002) found that the rotating cylinder stimulus failed to elicit significant CP in V1, suggesting that, unlike MT, this area does not contribute to the perception of such complex stimuli (two monkeys combined: mean CP = 0.48, prediction error in model with tailoring as sensitivity control = –2.2 SD).

Ghose and Harrison (2009) used a very brief stimulus presentation (60–83 ms) in a detection task but nevertheless reported high CP values in MT (two monkeys combined: mean CP = 0.64, prediction error in model with tailoring = +2.5 SD). The study employed a non-standard stimulus for MT: an array of 31 small Gabor patches that began drifting coherently at a specific point in time. A second array was presented symmetrically outside the neurons’ receptive fields, so the monkey could not predict which array would be behaviorally relevant on a given trial. Another unusual feature was that the stimuli were retinally stabilized to correct for fixational eye movements. Additionally, CP was estimated with a non-standard method that adjusted latency and time windows individually for each neuron to maximize CP values. Interestingly, and in contrast to our meta-analysis findings, the authors attributed the elevated CPs to the short stimulus duration – arguing that the highly informative, spatially compact, and temporally precise stimulus elicited strong, low-latency responses from MT neurons.

Palmer et al. (2007) reported an unusually high mean CP in V1 during a detection task (three monkeys combined: mean CP = 0.62; prediction error in model with inverse mean N/P ratio = +2.5 SD, model with tailoring = +2.8 SD). Despite these elevated CP values, no significant correlation between CP and neuronal sensitivity was found. In this study, monkeys were required to detect a low-contrast Gabor patch. Interestingly, the elevated mean CP values were observed only for trials in which the stimulus was present (hits vs. misses), which we chose to use in our meta-analysis. In contrast, trials without a stimulus (false alarms vs. correct rejections) showed much lower CP (mean = 0.52, we did not used this value in our analysis). The authors partly attributed this discrepancy to premature saccades and “fast guesses” during false alarm trials and also suggested that in the absence of a stimulus and low activity in V1, decision variability arises primarily from downstream noise sources.

### 8.6 On the Difference Between Mean P/N Ratio and Inverse Mean N/P Ratio

To clarify the difference between the mean P/N ratio and the inverse mean N/P ratio used in our analyses, we consider a simplified notation. Let the psychometric threshold *P*, which tends to be relatively constant across neurons within an experiment (especially after training), be replaced with a constant *a*. In contrast, let the neurometric threshold *N*, which varies across neurons, be denoted by the random variable *x*. This substitution allows us to isolate the effect of nonlinearity when aggregating across neurons.

#### Nonlinearity of Expectation and Reciprocals

Consider the following two expressions:

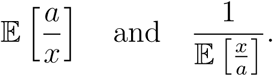

These two quantities are generally not equal and are not linearly related. This is a direct consequence of the convexity of the reciprocal function and the nonlinearity of expectation.

Specifically, Jensen’s inequality implies:

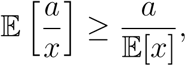

and similarly,

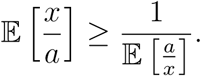

Therefore, it follows that:

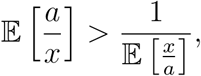

unless *x* is constant. Intuitively, when *x* takes small values (i.e., when neurons have high sensitivity), the reciprocal 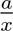 becomes large, which skews the expectation upward. This distinction is critical, as aggregating P/N vs. N/P ratios across neurons will yield systematically different results.

#### Implications for Regression Analysis

Let us now consider two linear regressions without intercept relating a dependent variable *y* to the two different versions of the sensitivity ratio:

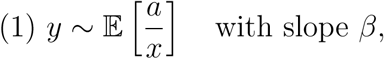

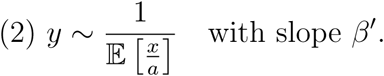

Because 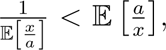 the independent variable in regression (2) tends to be smaller in magnitude. To achieve a similar fit to the dependent variable *y*, the model compensates with a larger slope. Thus, we typically find:

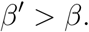

This means that using the inverse mean N/P ratio as a predictor (as done in some of our control analyses) tends to produce larger regression slopes compared to using the mean P/N ratio.

## Footnotes

1 To the best of our knowledge. Please contact us to report a study we might have overlooked.

2 Hereafter “single-electrode” denotes the physical recording technique, which should not be conflated with single-unit isolation.

3 Grouped under the “Learning and Engagement” category in Fig. 2.

